# SCENIC+: single-cell multiomic inference of enhancers and gene regulatory networks

**DOI:** 10.1101/2022.08.19.504505

**Authors:** Carmen Bravo González-Blas, Seppe De Winter, Gert Hulselmans, Nikolai Hecker, Irina Matetovici, Valerie Christiaens, Suresh Poovathingal, Jasper Wouters, Sara Aibar, Stein Aerts

**Affiliations:** VIB Center for Brain & Disease Research, Leuven, Belgium; Department of Human Genetics, KU Leuven, Leuven, Belgium; VIB Tech Watch, VIB Headquarters, Ghent, Belgium

## Abstract

Joint profiling of chromatin accessibility and gene expression of individual cells provides an opportunity to decipher enhancer-driven gene regulatory networks (eGRN). Here we present a new method for the inference of eGRNs, called SCENIC+. SCENIC+ predicts genomic enhancers along with candidate upstream transcription factors (TF) and links these enhancers to candidate target genes. Specific TFs for each cell type or cell state are predicted based on the concordance of TF binding site accessibility, TF expression, and target gene expression. To improve both recall and precision of TF identification, we curated and clustered more than 40,000 position weight matrices that we could associate with 1,553 human TFs. We validated and benchmarked each of the SCENIC+ components on diverse data sets from different species, including human peripheral blood mononuclear cell types, ENCODE cell lines, human melanoma cell states, and Drosophila retinal development. Next, we exploit SCENIC+ predictions to study conserved TFs, enhancers, and GRNs between human and mouse cell types in the cerebral cortex. Finally, we provide new capabilities that exploit the inferred eGRNs to study the dynamics of gene regulation along differentiation trajectories; to map regulatory activities onto tissues using spatial omics data; and to predict the effect of TF perturbations on cell state. SCENIC+ provides critical insight into gene regulation, starting from multiome atlases of scATAC-seq and scRNA-seq. The SCENIC+ suite is available as a set of Python modules at https://scenicplus.readthedocs.io.

## Introduction

Cell identity is encoded by gene regulatory networks (GRNs), in which transcription factors (TF) interact with sets of genomic regulatory regions to control the transcription of their target genes. These regulatory regions, called enhancers and promoters, contain specific combinations of TF binding sites, and thereby constitute *de facto* the GRN nodes that link the upstream TFs to their target genes. In depth knowledge of GRNs is important for the mechanistic understanding of biological aspects underlying development^1–3^ and evolution of cell types^4–7^ potentially providing insights into disease mechanisms^8^ and leading to better predictions of core regulatory complexes^9^ able to induce reprogramming. However, current knowledge of bona fide TF-target relations is still limited.

Experimental techniques including ChIP-seq have yielded a wealth of TF binding data sets, but this strategy works only on large amounts of homogeneous cells such as cell lines, or on bulk tissues that have a large proportion of one particular cell type, such as hepatocytes in the liver^10–14^. For tissues with a high diversity of cell types or cell states, such as the brain or a tumor, it remains challenging to experimentally map TF binding sites for each state. In addition, for most TFs, high-quality antibodies are lacking. Although alternative approaches have recently been described that have an increased cellular resolution (e.g., single-cell Cut&Tag^15^), or that rely on genetic tagging rather than antibodies (e.g., DamID^16^, nanoDam^17^), such methods are still difficult to scale to all TFs for all cell types. An alternative route for GRN mapping is to computationally model and predict TF-target gene relationships, for example using co-expression, a strategy that saw significant improvements during the last five years, thanks to scRNA-seq datasets. Large scRNA-seq atlases and time series data (e.g., trajectories) provide increased power to find TF-target associations based on co-variability of the TF and the candidate target gene^18^. However, such approaches have no notion of the genomic binding sites, and as such it cannot distinguish direct from indirect targets, and often yields high false positive prediction rates^18^. One way to circumvent this is to combine scRNA-seq co-expression networks with TF motif discovery, as is done in the SCENIC method^19,20^. This however requires searching for TF binding sites in a large sequence space around each co-expressed gene, and even when 10-20kb is included per gene, it represents a small portion of a gene’s putative regulatory space^21,22^.

With single-cell chromatin accessibility data available, for example obtained by scATAC-seq, the accuracy of TF binding site predictions can be improved significantly, even reaching similar precision and recall compared to ChIP-seq^23^. Genomic regions that are specifically accessible in a cell type often represent enhancers, and studies have shown that co-accessible regions are enriched for TF binding site combinations^3,22, 24–26^. Remaining challenges for this scenario are to accurately associate enriched motifs with candidate TFs; to expand the currently available motif databases so that more TFs can be identified; and to link predicted TF binding sites with candidate target genes, even if they are located far away from the target’s transcription start site^21,22^. To tackle these challenges, we developed three new Python packages that are combined into a computational eGRN prediction framework, called SCENIC+.

## Results

### SCENIC+ uses more than 30,000 TF motifs to predict eGRNs

SCENIC+ takes as input either paired or unpaired scRNA-seq and scATAC-seq data, as well as cell type annotations, and performs three analysis steps (Fig 1a, S1). Firstly, candidate enhancers that are co-accessible in certain cell types and/or states are identified by topic modelling. For this, we implemented a new Python package, called pycisTopic, that runs faster than the original cisTopic version^24^, while retaining comparable accuracy (Fig 1b,c, S2). Secondly, for each topic and all sets of differentially accessible regions (DARs, see Methods) for each cell type, the DNA sequences of its contributing regions are analyzed by motif discovery. To improve both precision and recall for motif enrichment and TF prediction, we created a new motif compendium and implemented a Python package for motif discovery called pycisTarget, that implements the cisTarget ranking-and-recovery based algorithm^19,27–29^, as well as two other motif discovery methods (Differential Enrichment of Motifs (DEM) and Homer^30^, see Methods). The motif compendium was generated by a meta-analysis of 29 motif collections, totaling 34,524 unique position weight matrices (Methods, S3a,b). A two-step clustering strategy (Methods) yielded 12,935 motif clusters, of which 1,985 contain more than 1 motif (with a mean of 5.8 motifs per cluster) (Fig 1d, S3c,d). By extrapolating TF annotations to similar motifs, we identified 1,553 human, 1,357 mouse and 467 Drosophila TFs with at least one motif cluster (with an average of 5, 6, and 2 motifs per TF respectively; Fig 1e, S3e). To our knowledge, this is the largest motif database, with the highest number of represented TFs. Given this resource, we asked how to optimally combine multiple motifs per TF. We found that, rather than defining an “archetype” (or average) motif per TF, maintaining all motif variations yielded a significantly higher recall and precision, on 309 ChIP-seq data sets, and this approach also outperforms previously existing methods such as Homer^30^ (Fig 1f, Fig S3f-h). Technically, pycisTarget exploits all motif variations by implanting each motif as a hidden state in a Hidden Markov Model (HMM), and each candidate sequence receives a log-likelihood ratio (LLR) score per motif cluster (Fig 1d, Methods). Significantly enriched motifs are identified for each set of co-accessible regions through an area under the recovery curve along the LLR-based ranking, and the optimal subset of sequences (called cistrome) is identified by a leading-edge analysis (Methods). DEM performs pairwise comparisons between region sets using these LLR scores as well (Methods, Fig 1d).

**Figure 1.**
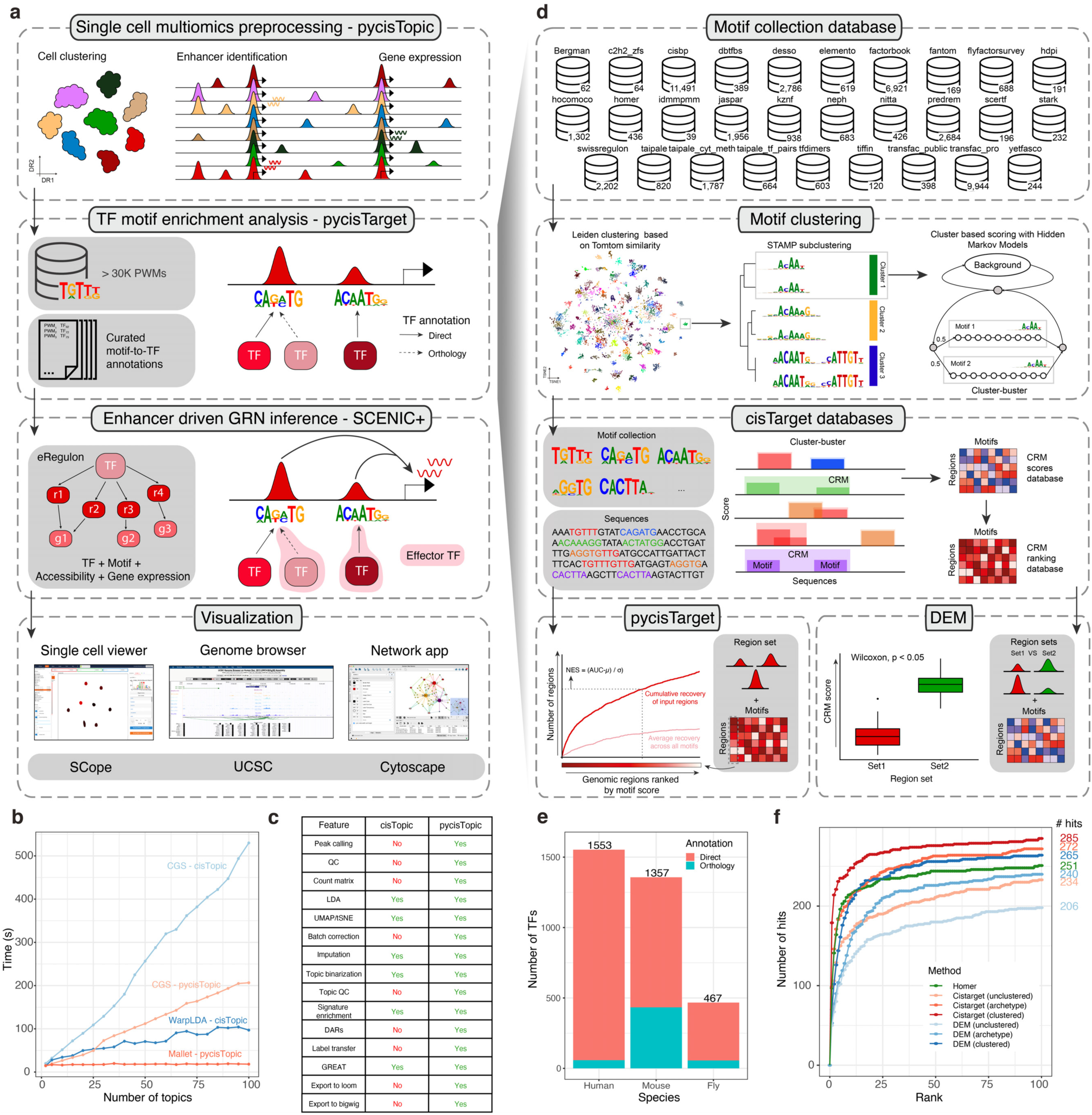
The SCENIC+ workflow and motif collection. **a**. In the SCENIC+ workflow, enhancer topics and DARs are first inferred using pycisTopic. pycisTarget then identifies modules for which the regulator’s binding motif is significantly enriched across regions, creating cistromes with only directly bound regions. SCENIC+ integrates region accessibility, TF and target genes expression and cistromes using GRNBoost to infer enhancer-GRNs, in which TFs are linked to their target regions, and these to their target genes. SCENIC+ can export results to loom files compatible with Scope^32^, bigwig and bigBed formats compatible with UCSC, and Cytoscape networks. **b.** Running time comparison per topic model using cisTopic with Collapsed Gibbs Sampling and WarpLDA (blue) and pycisTopic with Collapsed Gibbs Sampling and Mallet (red) for parameter optimization. **c.** Feature comparison between cisTopic and pycisTopic. **d.** The SCENIC+ motif collection includes 34,524 unique motifs gathered from 29 motif collections, which were clustered with a two-step strategy in which motifs are first clustered using Leiden clustering on the -log10 Tomtom q-value matrix, and these clusters are then refined using STAMP. Region scoring is performed using Hidden Markov Models (HMMs) models with an updated version of Cluster-Buster, in which clusters are scored using all the motifs as states instead of using a consensus motif. Ranked scores are used as input for pycisTarget, in which a recovery curve is built for each motif and region set. The Normalized Enrichment Score (NES) indicates how enriched a motif is in the region set. Scores are used for Differentially Enriched Motifs (DEM) analysis, in which a Wilcoxon test is performed between foreground and background region sets to assess enrichment. **e.** Number of TFs in the SCENIC+ motif collection annotated by direct evidence or orthology. **f.** Recovery of TFs from ENCODE ChIP-seq data using different motif enrichment methods, namely Homer, pycisTarget and DEM; and databases. The unclustered databases includes all annotated motifs before clustering (singlets), the archetype databases use the consensus motifs of the clusters based on STAMP, and the clustered databases uses the motif clusters, scoring regions using all motifs in the cluster. The x-axis shows the positions in which the TF targeted in the ChIP-seq experiment can be found, and the y-axis shows the cumulative number of TFs that are found in that position.

In the third step, SCENIC+ combines the gene expression values, the denoised region accessibility, and the cistromes to predict TF-region-gene triplets. Region-to-gene (default up to 150kb from the transcription start site) and TF-to-gene relationships are inferred using Pearson correlation and a tree-based regression approach, namely Gradient Boosting Machines (GBMs), to assess both linear and non-linear relationships^31^. These measurements are used in an enrichment analysis approach to recover the optimal set of target genes and regions for each TF (Methods). A TF with its set of predicted target enhancers and predicted target genes is called an enhancer-regulon (eRegulon), and all eRegulons are combined per cell type into an eGRN, and visualized in UCSC Genome Browser, SCope, and Cytoscape (Fig 1a).

### Illustration of SCENIC+ on PBMC multiome data

To showcase and validate SCENIC+ on a biologically complex data set, we analyzed a publicly available single cell multiomics data set containing 9,409 cells from human Peripheral Blood Mononuclear Cells (PBMCs) (Fig. 2a and Fig. S4a-c). SCENIC+ identified 53 activator TFs (Fig. 2b, S4d), targeting a total of 23,470 regions (out of 331,691 consensus peaks) and 6,142 genes (out of 16,351 genes). 5,469 genes (89%) have between 1 and 10 predicted enhancers and 9,347 (40 %) enhancers are predicted to regulate only the closest gene while some enhancers skip up to 20 genes (Fig. 2c).

**Figure 2.**
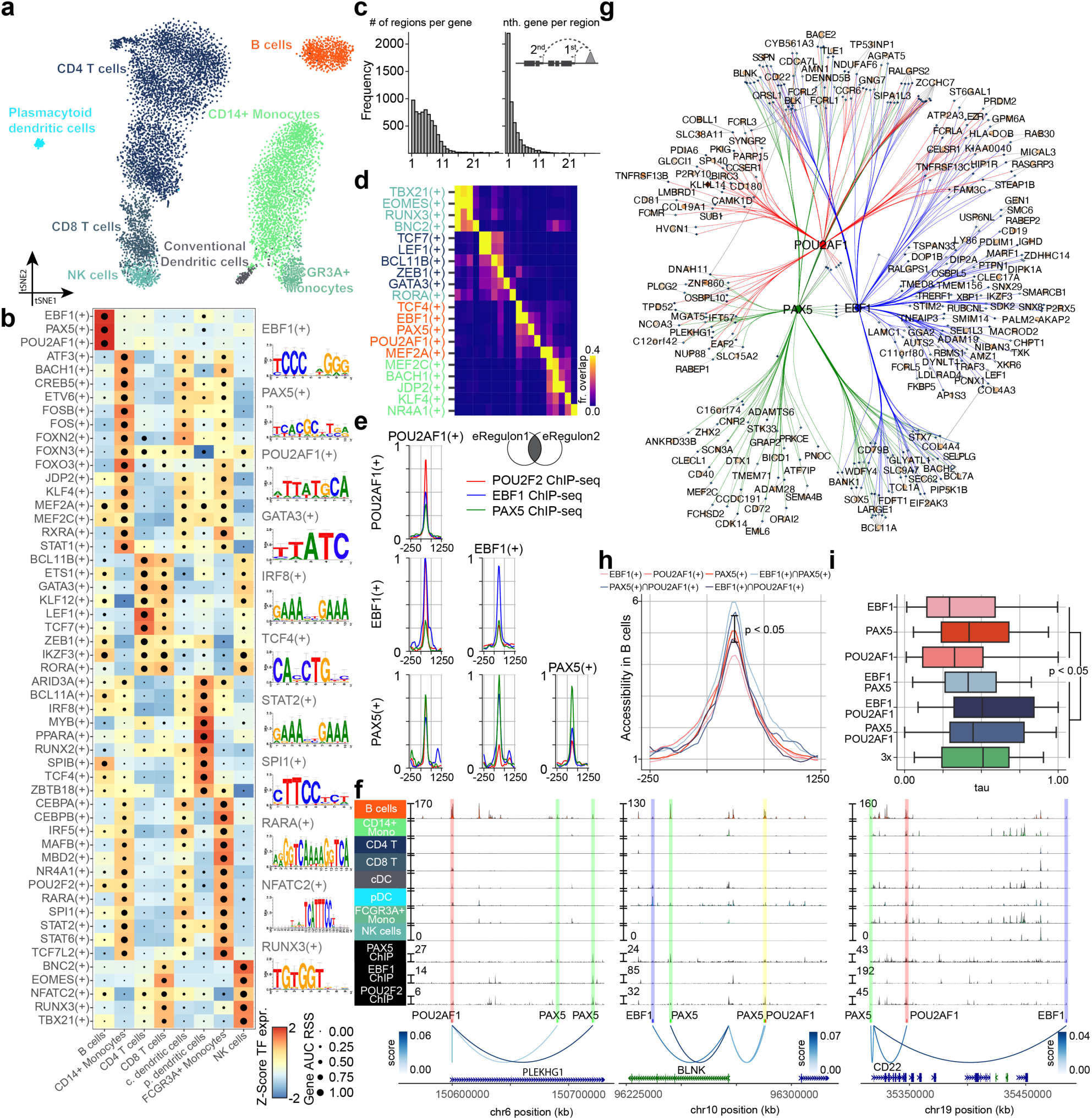
SCENIC+ analysis on peripheral blood monocyte multiome dataset. **a.** tSNE dimensionality reduction of 9,409 cells based on target gene and target region AUC scores of eRegulons. **b.** Heatmap-dotplot showing TF expression of the eRegulon on a color scale and cell type specificity (RSS) of the eRegulon on a size scale. **c.** Distribution of the number of regions linked to each gene and distribution of rank based on absolute distance between region and gene for each region. **d.** Overlap between target regions of eRegulons, overlap is divided by the number of target regions of the eRegulon in each row. **e.** Aggregated ChIP-seq signal of EBF1, PAX5 and POU2F2 ChIP-seq in GM12891 on target regions of either EBF1, PAX5 or POU2AF1 and combinations of two of these factors. **f.** Chromatin accessibility profiles across cell types and ChIP-seq signal of EBF1, PAX5 and POU2F2 ChIP-seq in GM12891 in the following regions: chr6-150562217-150730918, chr10-96226082-96316945 and chr19-35316521-35485142. Region-gene links are shown using arcs the color of which represents region-gene gradient boosting machine feature importance scores. Predicted target sites of eRegulons are shown using colored ticks and semi-transparent boxes. **g.** Visualization of eGRN formed by EBF1, PAX5, POU2AF1 and POU2F2 TF target nodes are restricted to highly variable genes and regions. **h.** Accessibility in B cells of target regions of either EBF1(+), PAX5(+) or POU2AF1(+) and combinations of two of these factors. Mann-Whitney U test (two-sided; p value 0.0336) between the average peak value of regions predicted to be targeted by one factor (n = 497) and regions predicted to be targeted by two factors (n = 70). Averages are indicated using line with bars. **i.** Distribution of cell type specificities, as measured by tau factor, of target genes of EBF1, PAX5 or POU2AF1 and combinations of two and three of these factors. Mann-Whitney U test (two-sided, p value 0.0062) between the average of genes predicted to be regulated by one factor (n = 127) and genes predicted to be regulated by two or three factors (n = 125). pDC: plasmacytoid dendritic cell, cDC: conventional dendritic cell.

SCENIC+ recovers well-known master regulators of B cells (EBF1, PAX5 and POU2F2/POU2AF1), T cells (TCF7, GATA3 and BCL11B), NK cells (EOMES, RUNX3 and TBX21), dendritic cells (SPIB and IRF8), and monocytes (SPI1 and CEBPA) (Fig 2b)^33–37^. For all these TFs, except EBF1, SCENIC+ predicted that they directly activate their own expression. Interestingly, for B cells SCENIC+ found both POU2F2 as well as its transcriptional co-activator POU2AF1^38^. We find that most of the top 5 specific TFs of each cell type show co-binding to shared enhancers. Such cooperativity is not observed for TFs that are not specific for the same cell type (Fig. 2d).

For B cells, SCENIC+ suggests cooperativity between EBF1(+), PAX5(+) and POU2F2/AF1(+) (Fig. 2d, g and Fig. S4i, j), with a strong overlap of their predicted target enhancers with EBF1, PAX5 and POU2F2 ChIP-seq data (Fig. 2e, f). We observed increased accessibility of enhancers predicted to be targeted by multiple of these TFs compared to enhancers with a single TF input (Fig. 2h), and increased cell type specificity of co-regulated target genes compared to genes regulated by a single TF (Fig. 2i).

In conclusion, SCENIC+ infers key regulators of different PBMC types and its accurate TF-region and TF-gene predictions can be exploited to infer enhancer and TF cooperativity.

### Validation and benchmark of SCENIC+ predictions using ENCODE data

To validate all components of SCENIC+ eGRNs (TFs, target regions, target genes and region-to-gene relationships) we used data from eight ENCODE Deeply Profiled Cell Lines^12,13^, namely MCF7 (breast cancer), HepG2 (hepatocellular carcinoma), PC3 (prostate cancer), GM12878 (B cell), K562 (leukemia), Panc1 (pancreatic cancer), IMR90 (lung fibroblast) and HCT116 (colon cancer). These data include paired bulk ATAC-seq and RNA-seq profiles for each cell line, 309 high quality ChIP-seq experiments, 157 TF knockdown experiments followed by bulk RNA-seq, and Hi-C data for 5 of the cell lines. We simulated 500 single-cell multiomics profiles by randomly sampling 50,000 reads and 20,000 fragments from each bulk RNA-seq and ATAC-seq profiles, respectively; resulting in a data set with 4,000 simulated single-cell cells. We also ran other methods that predict GRN/eGRNs using multiomics data and TF motifs as input, namely CellOracle^39^, Pando^40^, FigR^41^, and GRaNIE^42^; and as a baseline we included an updated version of SCENIC, which uses scRNA-seq and TF motifs but no chromatin accessibility^19,20^ (Fig 3a,b, Suppl Note 1).

**Figure 3.**
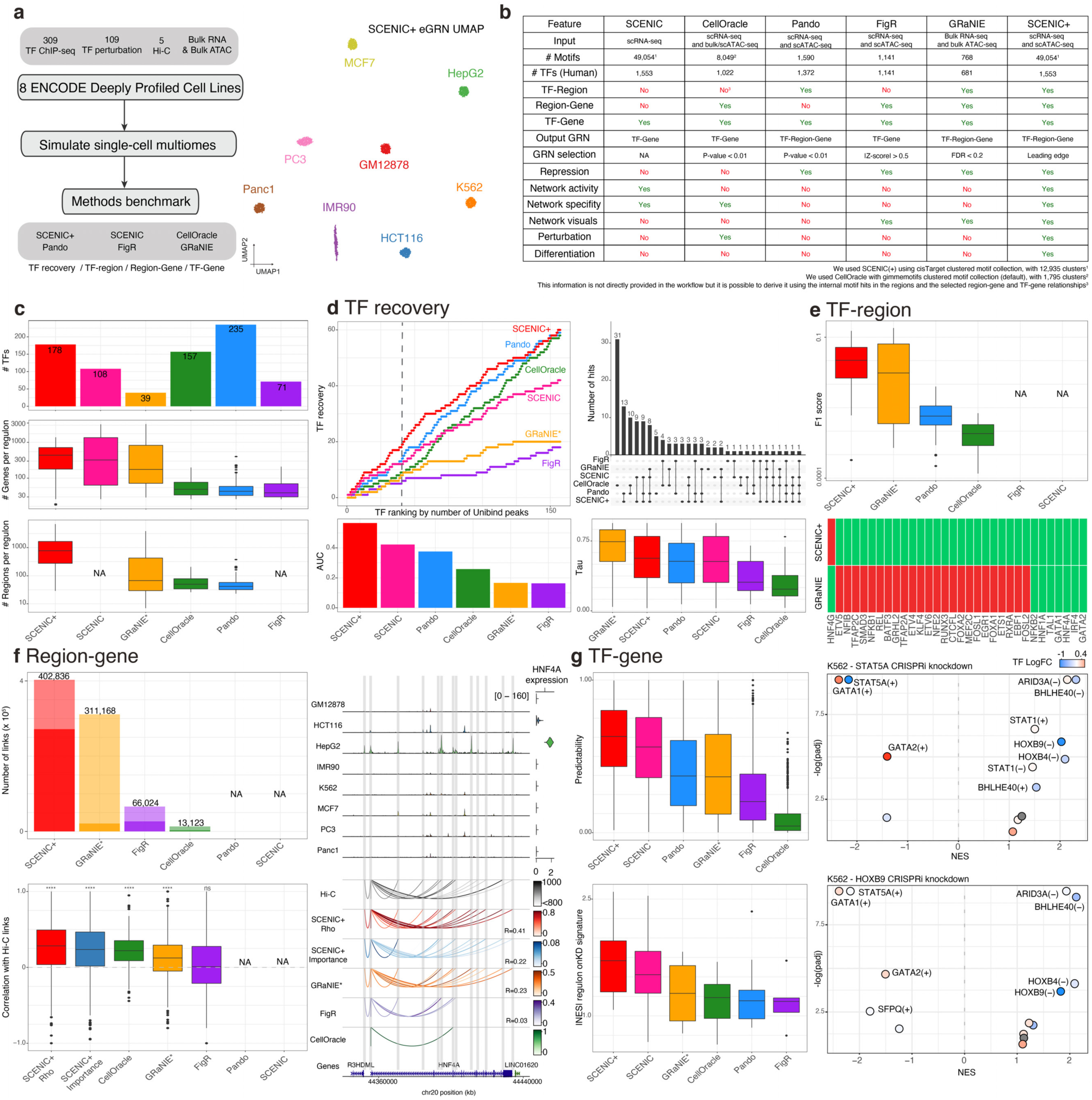
Benchmark of SCENIC+ and other single-cell multiomics GRN inference methods using ENCODE Deeply Profiled Cell Lines. **a**. Diagram showing benchmark data and strategy accompanied by the eGRN-based UMAP resulting from the analysis of the data with SCENIC+ (4,000 simulated cells) **b.** Table recapitulating main features of the different methods used in the benchmark **c.** Number of TFs recovered per method (top), and distributions showing the number of genes and regions found in each regulon per method (middle and bottom, respectively). **d.** Right: Cumulative TF recovery for each method using as x-axis TFs ranked based on the number of Unibind peaks and AUC values per method using the top 40 TFs. Top left: Overlap between the TFs recovered from the ranking for each method. Bottom left: Tau values distributions for the TFs recovered by each method. **e.** Top: F1 score distributions resulting from the comparison of the target regions found by the methods and Unibind. Bottom: Heatmap indicating whether a TF is found (green) or not (red) by each method (F1 score > 0.03). FigR and SCENIC are excluded from the comparison since they do not report TF-region relationships. **f.** Top left: Bar plot showing the number of region-gene links found by each method. Non-transparent bars show the number of links found in the eGRN, transparent bars show the number region-gene links before the eGRN construction. Bottom left: Correlation between Hi-C links for the top 100 markers genes for each of the cell lines where Hi-C is available (namely IMR90, GM12878, HCT116, HepG2 and K562) and the region-gene scores from different methods. Left: Example on the HNF4A locus depicting chromatin accessibility profiles and gene expression on the different cell lines and Hi-C and region-gene links for each method. Pando and SCENIC are excluded from the comparison since they do not report (unique) region-gene relationships. **g.** Top left: Box plot depicting the correlation between observed and predicted gene expression values using the eGRNs inferred from each method. Bottom left: NES distribution based on GSEA analysis using TF knockdown data as ranking and target genes derived by each method as gene sets. Left: Examples on K562 upon STAT5A (top) and HOXB9 (bottom) knockdowns showing GSEA -log10 adjusted p-value and NES for different eGRNs found by SCENIC+. NA is used in the panels when that information is not available when using that method. GRaNIE* indicates it was run with simulated single-cell data instead of bulk.

We first assessed how many TFs were recovered by each method, and how many target regions and genes were predicted per TF (Fig 3c, S5a-e). SCENIC+ identifies 178 TFs across the eight cell lines (Fig S5, S6). These include most of the known lineage TFs, such as GATA1/TAL1/MYB/LMO2 for K562^43–45^, or HNF1A/HNF4A/FOXA2/ CEBPB for HepG2^46^, or ESR1/GRHL2 for MCF7^47^. Vice versa, of the 228 marker gene TFs (Log FC > 3, padj < 0.05) in total 42% are found as SCENIC+ regulator. GRaNIE, FigR, and Pando identify fewer TFs (39, 71 and 157 respectively), while CellOracle identifies more TFs (235). The latter can be explained because CellOracle does not focus on cell type specific TFs, but also predicts broadly expressed TFs (e.g., GABPA, YY1, SP1), while the other methods, primarily SCENIC+ and GRaNIE, predominantly predict cell type specific TFs (showing higher tau values for specificity) (Fig 3d, S5e). To further quantify how many of the TFs identified by SCENIC+ are relevant, we tested how many of them have high quality ChIP-seq data with direct binding evidence (through UniBind^10,11^). SCENIC+ showed the best recovery of ‘directly bound TFs’ in the corresponding cell line, followed by SCENIC and Pando (Fig 3d, S5b-d).

Next, we assessed the number and quality of the predicted target regions per TF. Here, we used the predicted TF binding sites from UniBind as silver standard, which are derived from 309 ChIP-seq data sets in these cell lines. SCENIC+ recovers more regions (1,152 average) compared to other methods (ranging between 51-495) (Fig 1c), and these regions have a higher overlap with ChIP-seq peaks (Fig 1e, S5f). Note that only a small fraction (less than 8%) of ChIP-seq peaks can be recovered by any eGRN prediction method; this is likely due to the fact that many ChIP-seq peaks are not located close to a co-expressed target gene. In that sense, eGRN mapping selects only those TF binding sites that are predicted to play a functional role in regulating target genes. For some TFs, the predicted target regions overlap with multiple ChIP-seq datasets, which is due to shared/ common binding sites, as confirmed by overlapping ChIP-seq peaks (Fig S5g). Next, to assess the quality of the predicted region-gene associations we made use of deeply sequenced Hi-C data on five of the cell lines (see Methods). SCENIC+ predicts 402,838 links between a region and a putative target gene, having an average correlation of 0.25 with HiC interactions (Fig 3f, Fig S5h). The other methods identify fewer links (between 13,123 and 311,168) and show a lower correlation with HiC (Fig 3f, S5h).

Validation of the predicted target genes for each TF is more difficult compared to the assessment of TF relevance or TF binding sites, because only few experimentally validated target genes are available, for any TF. We reasoned that correct TF-gene predictions would result in accurate prediction of target gene expression, given only the expression of upstream TFs as input. To this end, we used 80% of the cells to train a regression model using Gradient Boosting Machines (GBM) and used the trained model to predict the remaining 20% of cells. For SCENIC+, predicted gene expression correlated on average 0.61 with the real expression vector, while this correlation was lower for Pando, GRaNIE, FigR and CellOracle (Fig 3g, S7a,b). An alternative approach to validate TF target genes is to examine changes in gene expression after the knock-down of a TF. Even though TF perturbations result in expression changes of both direct and indirect target genes (eGRNs only contain direct targets), one may expect the direct targets to show significant changes after a TF knockdown. Across 157 TF perturbation data sets with bulk RNA-seq data on our eight ENCODE cell lines, SCENIC+ regulons showed the highest enrichment by GSEA (average NES 1.66, Fig 3g, S7c-f).

In conclusion, SCENIC+ builds high quality GRNs with accurate enhancer and target gene predictions. In these tasks, SCENIC+ shows the best performance compared to state-of-the-art methods and provides the most insights in TF-region-gene regulation mechanisms.

### SCENIC+ simulates phenotype switching of cancer cell states

Gene regulatory network analysis of cancer cells holds promise to identify stable (attractor) cell states and their regulators. As a case study we generated a new scATAC-seq data set for nine melanoma cell lines that represent different melanoma states^48,49^ and combined this data set with previously published scRNA-seq data for the same lines^49^. SCENIC+ identified 42 high-quality activators (Fig. S8a), and cells clustered in three states based on the activity of these eRegulons (Methods, Fig. 4a and supplementary Fig. S8b, c). SCENIC+ was able to recover known regulators for the melanocytic (MEL) state (MITF, SOX10, TFAP2A and RUNX3), the mesenchymal (MES) state (JUN, NIFB and ZEB1) and finally, the intermediate sub-state of MEL which is governed by the MEL TFs, supplemented with SOX6, EGR3 and ETV4 activity (Fig. 4b and Fig. S8d)^49–51^. Because SCENIC+ uses information from both gene expression data and motif enrichment analysis it can aid with the discovery of the identity of the TF that binds to cell type specific enhancers. For example, it was previously suggested that RUNX motifs are part of the MEL enhancer code^25,49^ but which member of the RUNX family was unclear. Using SCENIC+, we predict that it is most likely RUNX3 (Fig. 4b).

**Figure 4.**
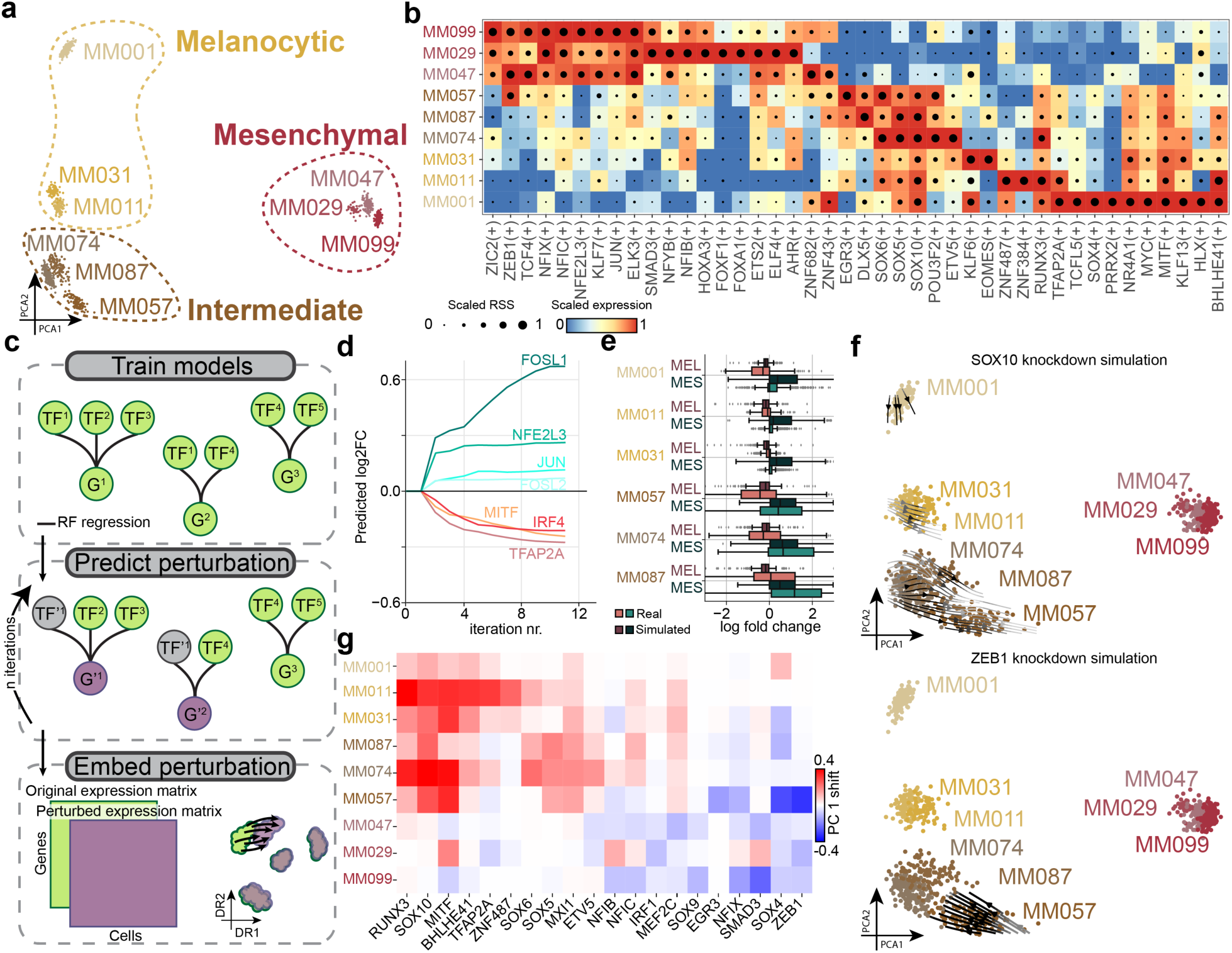
SCENIC+ analysis using separate scATAC-seq and scRNA-seq data on a mix of human melanoma lines. **a.** Principal Component Analysis (PCA) projection of 936 pseudo mutliome cells based on target gene and target region AUC scores of eRegulons. **b.** Heatmap-dotplot showing TF expression of the eRegulon on a color scale and cell type specificity (RSS) of the eRegulon on a size scale. **c.** Illustration on how predictions from SCENIC+ can be used to simulate TF perturbations; Top: SCENIC+ is used as a feature selection method and random forest (RF) regression models are fitted for each gene using TF expressions as predictors for gene expression; middle: the expression of (a) TF(s) is altered *in silico* and the effect on gene expression is predicted using the regression models, this is repeated for several iterations to simulate indirect effects; bottom: the original and simulated gene expression matrix are co-embedded in the same lower dimensional space to visualise the predicted effect of the perturbation on cell states. **d.** Predicted log2 fold change of mesenchymal (blue shades) and melanocytic (red shades) marker genes over several iterations of SOX10 knockdown simulation. **e.** Simulated and actual distribution of log2 fold changes of melanocytic and mesenchymal marker genes after SOX10 knockdown across several MM-lines. **f.** Simulated cellular shift after SOX10 and ZEB1 knockdown represented using arrows. Arrows are shaded based on the distance traveled by each cell after knockdown simulation. **g.** Heatmap representing the shift along the first principal component of each MM-line after simulated knockdown of several TFs.

Next, we examined TF cooperativity in the melanocytic state. It was previously shown that a core regulatory complex^9^ of SOX10, RUNX, MITF and TFAP2A regulates melanocytic-state-specific enhancers, and that most MEL enhancers have at least one SOX10 site^25^. Using SCENIC+ we recapitulate that many of the predicted SOX10 target regions are also predicted to be bound by either RUNX3, MITF or TFAP2A, or a combination thereof (Fig. S8e). Interestingly, while many enhancers are predicted to be bound by a combination of two of the factors, only very few are predicted to be bound by a combination of three or all four (Fig. S8e). We validated these findings using publicly available ChIP-seq data^52,53^ showing high and specific enrichment of ChIP-seq signal in enhancers predicted to be bound by either SOX10, MITF and TFAP2A or a combination of these factors (Fig. S8f).

It is known that melanoma cells can dynamically shift fate from melanocytic to mesenchymal and vice versa, driving metastasis and therapeutic resistance, a process called phenotype-switching^54^. Knockouts of specific TFs can drive this process^49^. To simulate this switch and prioritize TFs driving it using SCENIC+, we took inspiration from the recently published method CellOracle^39^. CellOracle exploits eGRN predictions to simulate the effect of perturbing a specific TF, using a linear model. We implemented an adapted version of this technique in SCENIC+. Basically, we use SCENIC+ for feature selection (i.e., the TFs per target gene) and train a random forest regression model for each gene to predict gene expression based on these predictors. After fitting the model, we use it to predict the effect of a TF perturbation on gene expression by simply setting the expression of the TF to zero. To include indirect effects, we predict new gene expression values for several iterations. Finally, we visualize the effect of the simulated perturbation by co-embedding the simulated gene expression matrix with the original one (Fig. 4c). As proof of principle, we simulated the effect of SOX10 KD in the MEL state. Interestingly, the simulated cells after SOX10 KD suggest that they switch to a more mesenchymal-like state, whereby mesenchymal genes are up-regulated, and melanocytic genes are down-regulated, and this effect stabilized after around 4 iterations of simulation (Fig. 4d). This predicted effect of SOX10 knockdown was strongest in the intermediate cell lines, and is recapitulated by experimental SOX10 KD, followed by RNA-seq^49^ (Fig. 4e and Fig. S8g). Encouraged by this result, we simulated cells for perturbations of all the identified TFs. Simulated knockdowns of RUNX3, SOX10 and MITF show the strongest potential to switch cells from MEL to MES; while knockdowns of ZEB1, SOX4 or SMAD3 may lead to the reverse switch from MES to MEL (Fig. 4 f, g). This strategy can thus be used to prioritize TFs regulating cell state and state transitions.

In conclusion, using eGRNs inferred by SCENIC+ we could recapitulate known melanoma cellular states, rediscovering many TFs which are known to underlie the identity of these states. Our analysis also highlights how SCENIC+ can be used in unpaired (i.e., non-multiome) scATAC-seq and scRNA-seq data while retaining accuracy. Furthermore, by using the networks inferred by SCENIC+ we could simulate TF perturbations, recapitulating known effects and suggesting new candidates for driving phenotypic switches.

### Conservation and divergence of eGRNs in the human and mouse cerebral cortex

The mammalian cortex contains a high diversity of excitatory (pyramidal) and inhibitory neurons, whereby the excitatory cell types are spatially organized in cortical layers^55,56^. Although several marker TFs have been described for certain cell types, little is known about how precise TF combinations, their binding sites, and their target genes underlie neuronal identity. Given the very high degree of conservation of cortical cell types between human and mouse^57^, we reasoned that two independent SCENIC+ analyses on human and mouse could reveal conserved, and thereby high-confidence, cortical eGRNs.

For the mouse cortex, we generated a new 10x single cell multiome data set with 19,485 cells, and for the human cortex we re-used a previously published multiome data set (SNARE-seq2) with 84,159 cells^57^ (Fig 5a-b). Basic quality metrics for both data sets are similar on the scRNA-seq part, with 6,392 and 5,747 UMIs (2,155 and 2,226 genes) detected per cell on average, for mouse and human, respectively. For the scATAC-seq part, we identified 568,403 and 697,721 accessible regions for mouse and human respectively (21,337 and 5,078 unique fragments per cell; median TSS enrichment of 8.9 and 3.5; 59% and 43% FRiP). Despite the differences in the number of cells and coverage, we were able to identify matching cell types in both species, including layer specific excitatory neurons, interneuron derived from the Medial and Caudal Ganglionic Eminescences (MGE and CGE, respectively), and non-neuronal populations (microglia, astrocytes, endothelial cells and oligodendrocytes and Oligodendrocyte Progenitor Cells (OPC)) (Fig 5a-b).

**Figure 5.**
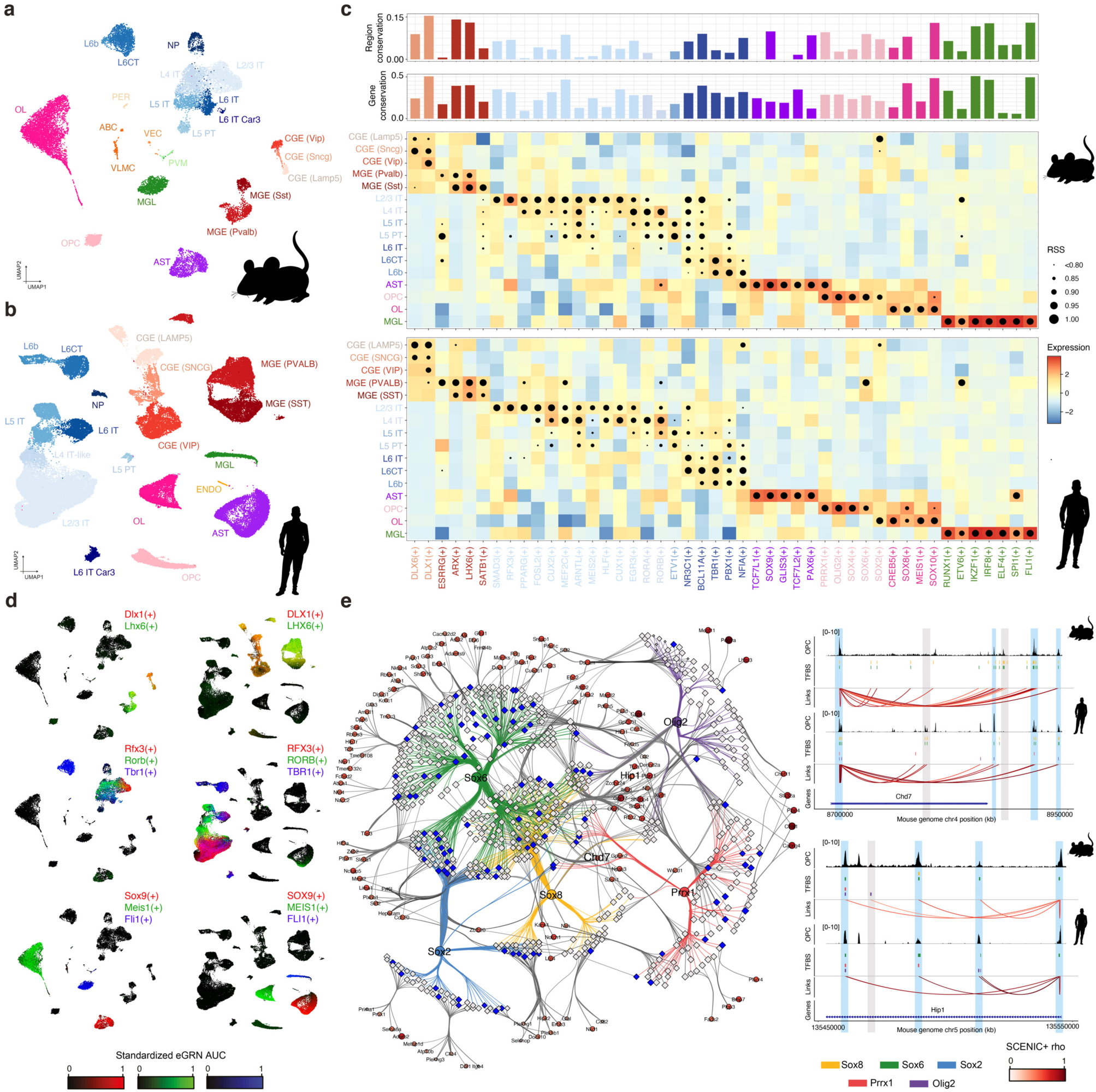
SCENIC+ reveals regulatory lexicon conservation across mammalian brains. **a.** SCENIC+ UMAP containing 19,485 cells from the mouse cortex processed with 10x multiomics. **b.** SCENIC+ UMAP containing 84,159 from the human motor cortex processed with SNARE-seq2. **c.** Dotplot for the mouse (top) and human (bottom) analyses showing the Regulon Specificity Scores (RSS) as dot size and TF expression as color (as background for each dot). The barplot above indicate the percentage of the regulon that is conserved in the other species, for predicted target regions (top) and target genes (bottom). **d.** Mouse and human UMAPs colored by AUCell enrichment of selected regulons using RGB encoding. **e.** Left: Mouse-based OPC (Oligodendrocyte Progrenitor Cells) eGRN with conserved TFs. Regions are shown with diamond shape, and their size represents the LogFC of the region accessibility in OPC compared to the rest of the cells. Regions conserved in the human brain are shown in blue, regions only found in the mouse analysis are shown in grey. Genes are shown with circular shape, and their color and size represent the LogFC of the gene expression in OPC compared to the rest of the cells. TF-region links are colored by TF, and region-gene links are colored by SCENIC+ correlation scores between the region accessibility and the target gene expression. Right: OPC coverage, TF binding sites and region-gene links in two loci, Chd7 and Hip1. Data is shown in the mouse genome (mm10), and human data has been liftovered (mm10). Peaks found in both human and mouse are highlighted in blue, while peaks only accessible in one of the species are highlighted in grey. ABC/VLMC: Vascular leptomeningeal cells, AST: Astrocytes, CGE: Caudal Ganglionic Eminescence, CT: Cortico-thalamic, ENDO: Endothelial cells, IT: Intratelencephalic, MGE: Medial Ganglionic Eminescence, MGL: Microglia, NP: Near-projecting, PER: Pericytes, PVM: Perivascular macrophages, PT: Pyramidal-tract, OL: Oligodendrocyte, OPC: Oligodendrocyte Precursor Cells, VEC: Vascular endothelial cells.

Using SCENIC+ we found 125 and 142 high quality regulons for mouse and human, respectively; out of which 60 are found in both species (Fig 5c,d, S9a,b). Importantly, there was a very high correlation between the matching cell types based on the enrichment of these shared regulons, showing that cell identity can be compressed by these 60 conserved TFs (Fig S9c). Similarly, regulons are enriched in the same cell types, with a median correlation of 0.80 when comparing their enrichment across cell types (with 53 regulons with a correlation above 0.6). In both species -and in agreement with literature-, we distinguished Dlx1/DLX1 and Dlx6/DLX6 as master regulators in CGE interneurons; Lhx6/LHX6, in MGE interneurons; Rfx3/RFX3, Cux1/CUX1, Cux2/CUX2, Mef2c/MEF2C, Egr3/EGR3, in excitatory neurons located in the upper cortical layers (L2/3 to L4); Rorb/RORB and Rora/RORA in L4 excitatory neurons; Etv1/ETV1 in L5 intratelencephalic (IT) and pyramidal-tract (PT) excitatory neurons; Tbr1/ TBR1 in L6 excitatory neurons; Sox9/SOX9 and Glis3/GLIS3 in astrocytes; Sox10/SOX10 in oligodendrocytes; Olig2/OLIG2 and Sox2/SOX2 in OPC and Spi1/SPI1 and Irf8/IRF8 in microglia, among others^56,58,59^ (Fig 5c,d, S9a,b). To assess whether species-specific TFs are not conserved or missed in one of the two analyses (i.e. due to data sparsity and/or the applied filters along the workflow) we converted the predicted target genes in the mouse regulons to human gene names and scored these regulons in the human data set. We found a high correlation between the mouse and the human enrichment values across cell types, with a median correlation of 0.79 (with 51 regulons with a correlation above 0.6), indicating that these regulators are likely also conserved. For example, while Pou3f1/POU3F1 and Fezf2/FEZF2, described regulators of L5 PT and L5/6 neurons respectively^56^, were only found in the mouse analysis; the human-based mouse regulons are enriched in the corresponding cell types in the human data set, matching with the TFs expression (Fig S9d).

Next, we asked whether the predicted target genes and regions are conserved between the two species. To do this, we mapped the human target genes to mouse orthologs (based on ENSEMBL); and the human regions to the mouse genome using LiftOver (see Methods). Out of the 102,746 regions found within the human eGRNs, 84,861 could be lifted over (82%), while only 69% of all accessible regions (697,721) could be lifted over. Out of these 84,861 orthologous mouse regions, 61,973 were accessible in the mouse cortex. In addition, 312,591 (out of 379,749) region-gene links from the human cortex could be lifted over, of which 283,900 corresponded to the same region-gene pair in the mouse cortex eGRN. On average, 28% of the target genes and only 6% of the target regions of a certain TF were conserved between the two species, with a strong correlation (0.68) between the conservation values across the region and gene based regulons. Among the most conserved regulons are Dlx1/DLX1 (51% and 15% of the target genes and regions, respectively), Lhx6/LHX6 (41% and 13%), and Arx/ARX (39% and 14%), which are key regulators of interneurons^60^. Other TFs with conserved regions and genes include Mef2c/MEF2C (47% and 9%), Cux1/CUX1 (36% and 4%), Smad3/SMAD3 (36% and 6%), Rfx3/RFX3 (32% and 7%) and Cux2/ CUX2 (32% and 3%); which are TFs involved in upper layer excitatory neurons^58^. In the deep layer neurons, TFs with conserved targets include Tbr1/TBR1 (31% and 3%) and Nfia/NFIA (33% and 8%). Slightly higher conservation was found for Fli1/FLI1 (51% and 13%) and SOX10 (54% and 13%) in microglia and oligodendrocytes, respectively^56,58,59^ (Fig 5c, SB1e-f). Thus, despite the high conservation of TFs per cell type, the target genes -and even more so the target regions-are less conserved. This has also been observed in previous studies, for example Bakken et al. reported 25% and 5% conservation of DEGs and DARs, respectively, across the cell types in the human and marmoset cortex^57^. Stergachis et al. performed a DNase I footprinting study across 25 mouse tissues and cell types, finding that only ∼20% of the TF footprints are conserved in comparison to human, while 95% of the TF code is shared^61^. Genomic relocation and evolution of TF binding sites and enhancers may partly explain these observations; however, the sparsity of the single cell data sets may also contribute to these findings, as we are only capturing a small fraction of the transcriptome and epigenome in each cell.

We further studied eGRN conservation in oligodendrocyte precursor cells, which includes Sox2/SOX2, Sox6/ SOX6, Sox8/SOX8, Olig2/OLIG2 and Prrx1/PRRX1 (Fig 5e). These TFs have been extensively described in literature as key drivers of OPC proliferation, migration, specification, quiescence, and differentiation^62,63^. Out of the 636 regions targeted by at least 1 of these TFs in mouse and linked to at least one of the 127 conserved target genes in both mouse and human OPCs, only 102 are conserved TF binding sites between the two species (16%). To further examine the relationship between target region conservation and TFBS presence, we zoomed in on two different loci, Chd7 and Hip1^64,65^. We distinguished 3 different patterns: 1) The peak and TFBS are present in one of the species, while in the other species there is no accessibility and no TFBS (2 cases in the Chd7 loci), 2) the peak and the same TFBSs are found in both species (2 cases in the Hip1 loci) and 3) the peak and at least one TFBS is shared between the two species, but additional not shared TFBSs can be found. For the latter case, we also observed variation in the peak (after liftover of the human profile to the human genome) when other different

TFBSs are found in the same peak between species (e.g., more accessibility in the species where additional TFBSs are found, or a different peak shape); while peaks with the same TFBSs have a similar shape (Fig 5e).

In conclusion, comparative analysis with SCENIC+ reveals TF lexicon conservation across mammalian brains, but divergence on their target genes and regions.

### GRN velocity

Single-cell RNA-seq data is often used to sample cells during a dynamic biological process, such as cellular differentiation. Reconstructing the most likely cellular trajectory from such data is challenging^66^. RNA velocity-based methods exploit cell-intrinsic features to model dynamic changes, namely the relative fraction of spliced and unspliced reads^67,68^. Recently, an approach has been described to use the temporal relationship between chromatin accessibility and gene expression based on single cell multiomics read outs to infer dynamics^69^. However, none of these approaches include gene regulatory information in their model. We reasoned that also regulatory relationships derived by SCENIC+ could provide additional intrinsic cues to predict cellular dynamics. For example, the expression of a TF may precede accessibility of its binding sites; and chromatin accessibility in turn may precede target gene expression^70^ (Fig 6a). The procedure we apply to quantify the putative differentiation force of a TF is described in the Methods section. Briefly, after ordering the cells by pseudotime, a standardized generalized additive model (GAM) is fit along the pseudotime axis for the TF expression, as well as for the activity of its regulon (joint expression of target genes; or joint accessibility of its target regions). Next, each cell is projected from its quantile in the GAM TF expression model to the same quantile in the GAM regulon curve (posterior in the pseudotime axis). The differentiation potential for this TF in a cell is then defined as the distance from the TF expression curve to its matching cell in the regulon AUC curve. Differentiation potential can be plotted as an arrow grid in any cell embedding (Fig 6a,b). Furthermore, since these arrows are calculated for each TF, they can be used to identify TFs with a certain direction, or with a particularly high potential. To test this method, we first applied it to a linear differentiation trajectory from OPC to mature oligodendrocytes in the mouse brain. This revealed a set of TFs (Olig2, Bcl6 and Prrx1) with arrows pointing “inwards” into the OPC state. Another set of TFs (Tcf7l2 and Sox10) showed a delay between TF expression and target gene, with arrows pointing towards early differentiating oligodendrocytes (also called Newly Forming Oligodendrocytes (NFOL)). A third set of TFs (Zeb2, Meis1 and Tcf12) was identified as potential drivers of the maturation process from NFOL to oligodendrocytes (Fig 6b-d). The predicted role of many of these TFs agrees with literature^62,63, 71–74^.

**Figure 6.**
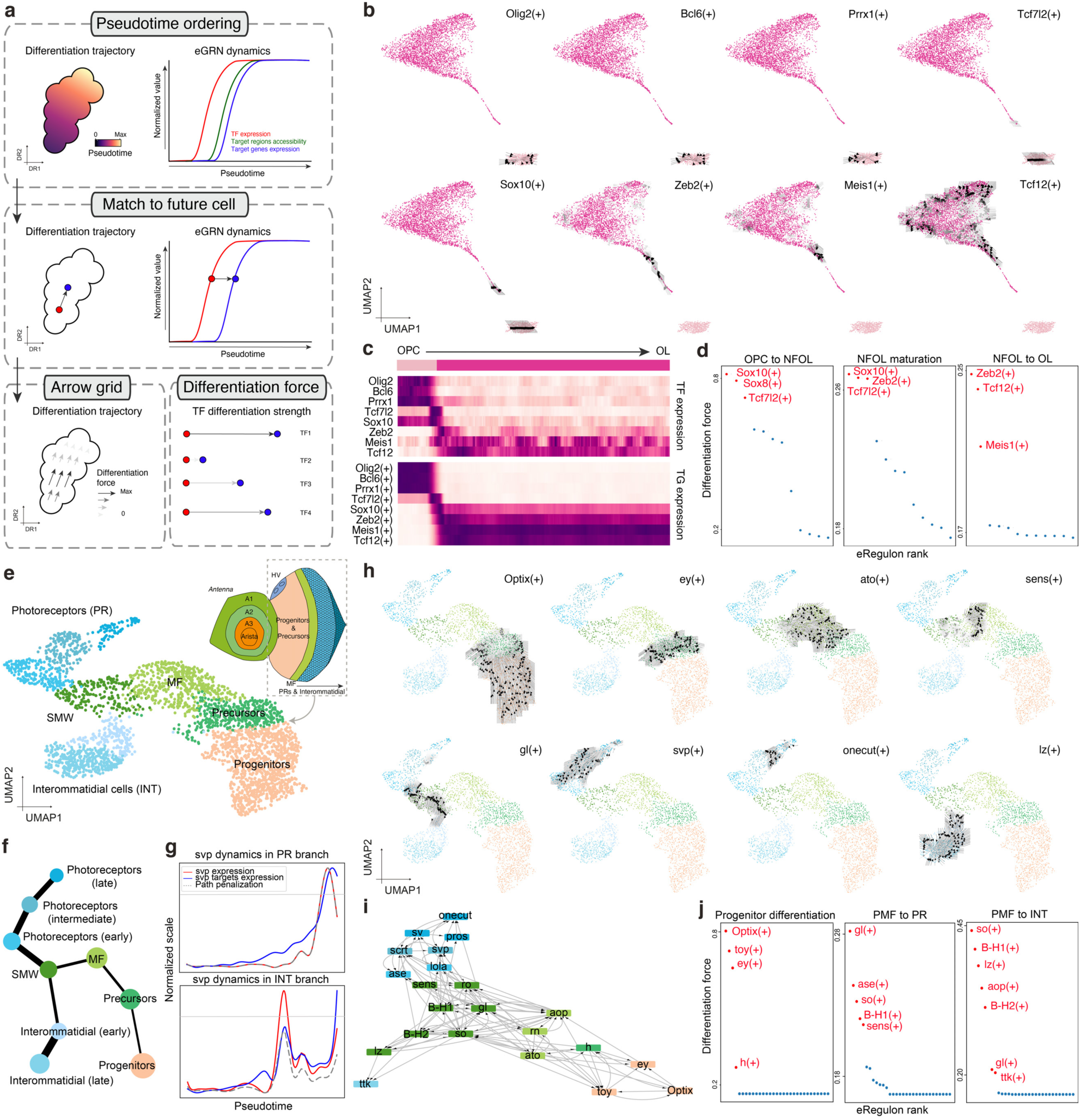
Identification of differentiation drivers from SCENIC+ enhancer-driven gene regulatory networks. **a.** Computational approach to infer differentiation drivers from a SCENIC+ analysis. First, differentiating cells are ordered by pseudotime. Second, for each TF, we fit a standardized generalized additive model (GAM) along the pseudotime axis for its expression and its target genes (or regions) AUC values and then map each cell in a certain quantile of the GAM TF expression curve with the one in the same quantile in the GAM regulon AUC curve. Finally, we define the differentiation force of a cell as the distance from the TF expression curve to its matching cell in the regulon AUC curve. Differentiation forces can also be plotted as an arrow grid in any dimensionality reduction of the data. **b.** Arrow grid representation along the differentiation of Oligodendrocyte Precursor Cells (OPC) to mature oligodendrocytes in the mouse cortex (4,435 cells). **c.** Heatmaps showing standardized TF expression (top) and regulon AUC values (bottom) along the oligodendrocyte differentiation pseudotime. **d.** Prioritization of differentiation forces different state transitions along the oligodendrocyte differentiation. **e.** SCENIC+ regulon UMAP with 3,104 pseudocells from the fly eye disc, together with a schematic representation of the fly eye-antennal disc. **f.** PAGA graph based on SCENIC+ regulons representing showing the different paths along the eye disc differentiation, from progenitors to photoreceptors or interommatidial cells, respectively. The width of the links represents the confidence in the presence of the connections. **g.** Representation of svp dynamics along the two paths in the eye disc differentiation. The grey horizontal line represents the TF expression threshold for arrows to be drawn. **h.** Arrow grid representation along the eye disc differentiation. **i.** GRN summarizing key drivers of differentiation in the eye disc along different states**. j.** Prioritization of differentiation forces in progenitors, and driving to differentiation into photoreceptors or interommatidial cells, respectively.

As a second application we analyzed a branched differentiation trajectory from progenitor cells to photoreceptors or interommatidial cells in the developing fly retina. As data set we re-analyzed our previously published scRNA-seq and scATAC-seq atlases of this developing tissue^22^, and we added an additional scATAC-seq run with 7,953 cells from the eye field only. We then used ScoMAP^22^ to spatially integrate the independent scRNA-seq and scATAC-seq data sets into an eye-antennal disc template with 5,058 pseudocells, each with single-cell multiomics-like information (Fig 6e). SCENIC+ detected 153 regulons, out of which 105 are active in the eye part (Fig S10a). TFs were also identified for the different antennal rings (e.g. Lim1 in A1, Hth in A1 and A2, Dll in A2, A3 and arista, and ss in A3 and arista^75,76^), for glia (e.g. repo^77^), the head vertex (e.g. oc^78^) and hemocytes (e.g. pnr^79^). Interestingly, the Cut (Ct) transcription factor, expressed in the antennae where it acts as repressor of the eye field, was indeed identified as candidate repressor, targeting 13 other TF (e.g., ss, ey, toy and Optix, in agreement with literature^80^) (Fig S10b-c).

We inferred a trajectory in the eye field based on the eRegulon activities using PAGA^81^, which runs from progenitor cells to precursors, then to the morphogenetic furrow (MF), the Second Mitotic Wave, and a branch point leading to either photoreceptor cells or interommatidial cells (Fig 6f). Strong differentiation arrows are found for Optix, Twin of Eyeless and Eyeless in progenitors, followed by Hairy, and Aop, Rotund and Atonal in the morphogenetic furrow. Bar-H1, Bar-H2, Sine Oculis, and Glass were found to trigger the differentiation process after the MF, both towards photoreceptors and interommatidial cells. Lozenge was found as the key force towards the interommatidial cell identity, and Tramtrack as key driver in their maturation. In the photoreceptor branch, Senseless and Rough were identified as key regulators of the early photoreceptor differentiation, followed by Asense, Lola, Seven-up, Scratch, and afterwards Shaven, and Onecut in mature photoreceptors (Fig 6g-j, S10d). Importantly, these results recapitulate the previously described differentiation cascade in the eye disc^22^.

### Spatial GRN analysis

Spatial omics enable single-cell profiling while maintaining the positional context of cells in a tissue. Ultimately, we could envision eGRN mapping by directly using spatial multiomics data, but currently available high throughput technologies are limited to spatial transcriptome profiling^82^. Nevertheless, with a SCENIC+ analysis of a matching single-cell multiomics data set of the same tissue, we were able to map eGRN activity onto the tissue, via the spatial transcriptome. To illustrate this, we used a publicly available sc-multiome data set of the human cerebellum with matched 10x Visium data from 10x Genomics. From the scRNA-seq data of the multiome we annotated nine main cell types, namely OPC, oligodendrocytes, Purkinje cells, granule cells, inhibitory neurons (Vip+, Sst+, Pvalb+ and Sncg+), and Bergman glia (Fig 7a). Despite the small number of cells in this data set

**Figure 7.**
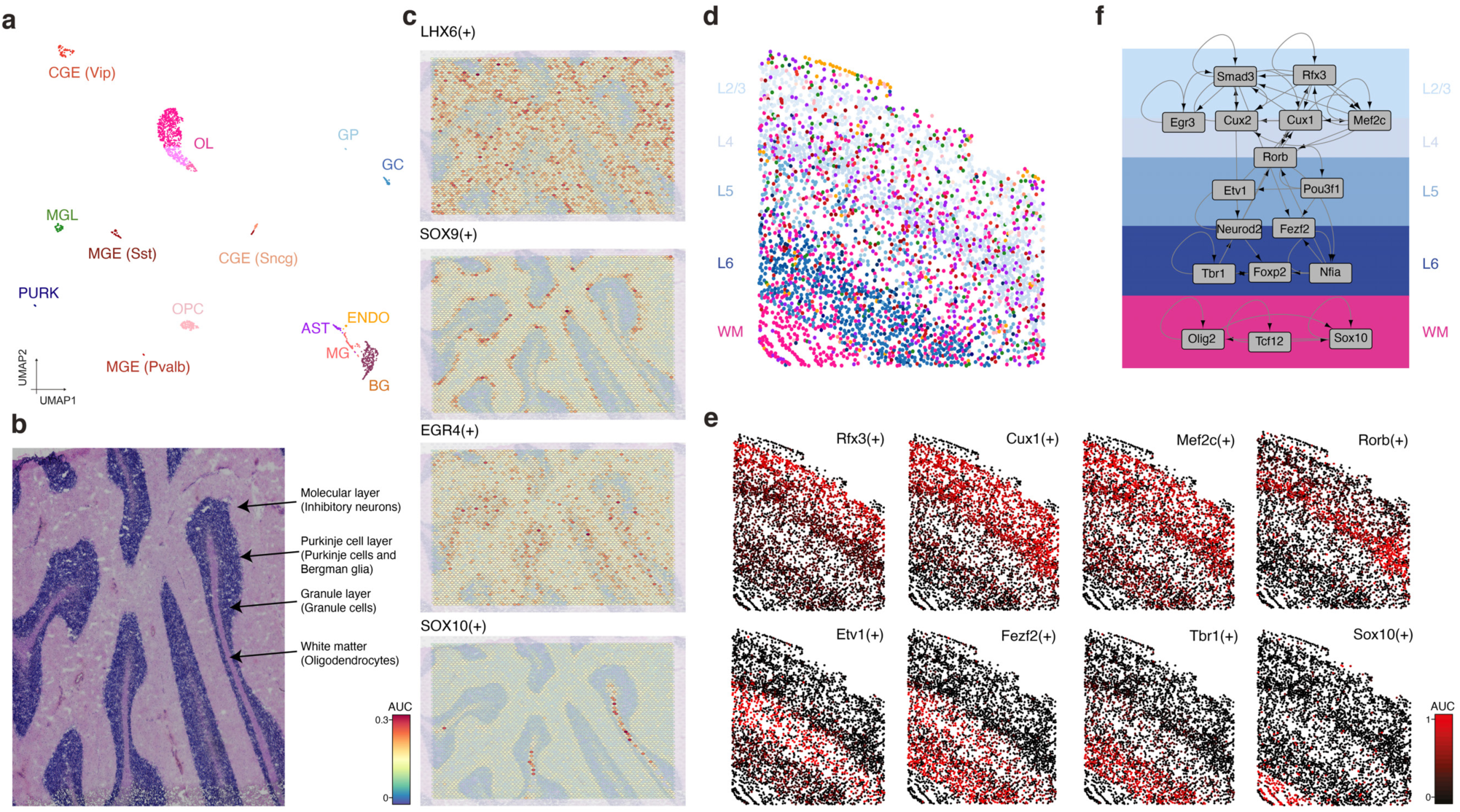
Spatial visualization of enhancer-GRNs in the mammalian brain. **a.** SCENIC+ UMAP containing 1,736 cells from the human cerebellum. **b.** Human cerebellum 10x Visium slide annotated with anatomical regions in the cerebellum. **c.** Visualization of regulons AUC enrichment on the 10x Visium data. **d.** Mapping of cell types in the mouse cortex into our smFISH map using Tangram^87^. **e.** Visualization of regulons AUC enrichment in our smFISH map of the mouse cortex. **f.** Representative layer specific gene regulatory network. The network depicted from L2/3 to L6 corresponds to excitatory neurons, while in the white matter corresponds to oligodendrocytes. AST: Astrocytes, BG: Bergman Glia, CGE: Caudal Ganglionic Eminescence, ENDO: Endothelial cells, GC: Granule Cell, GP: Granule cell Progenitor, MGE: Medial Ganglionic Eminescence, MGL: Microglia, MG: Muller Glia, OL: Oligodendrocyte, OPC: Oligodendrocyte Precursor Cells, PURK: Purkinje cells, WM: White Matter.

(1,736), SCENIC+ identified 111 regulons, including DLX1 and DLX6 in CGE interneurons (80 cells), LHX6 in MGE interneurons (134 cells), NEUROD2 and EGR4 in granule cells (76 cells), NEUROD1 in Purkinje cells (20 cells), PRRX1 and OLIG2 in OPC (157 cells), SOX10 and TCF12 in mature oligodendrocytes (823 cells), SPI1 and RUNX1 in microglia (67 cells), and NFIA and SOX9 in the Bergman glia (276 cells)^56,60, 83–85^. These results show that, even in small data sets and for rare populations, SCENIC+ can infer bona fide gene regulatory networks (Fig S11). We then scored SCENIC+ regulons onto the 10X visium spots using AUCell. This shows the LHX6 regulon enriched in the molecular layer of the human cerebellum, where interneurons reside; SOX9 on the Purkinje cell layer, where the Bergman glia are found; EGR4 in the granule cell layer, and SOX10 in the white matter, which is populated by oligodendrocytes (Fig 7b-c), in agreement with literature^85,86^.

As second case study we spatially analyzed the eGRNs of the mouse cortex multiome data that we identified above. To this end, we generated a smFISH atlas using 100 genes (based on marker genes per cell types and literature (see Methods)). We then used Tangram^87^ to transfer cell type annotations and gene expression to the segmented cells in the smFISH map (Fig 7d), and then scored the SCENIC+ regulons using AUCell on the imputed complete transcriptome. Congruently with our previous analyses, this shows Rfx3 activity in the L2/3 area, Cux1 and Mef2c from L2/3 to L4, Rorb in L4, Etv1 in L5, Fezf2 in L5 and L6, Tbr1 in L6, and Sox10 in the white matter (Fig7e).

## Discussion

Mapping a gene regulatory network (GRN) underlying a biological process can be useful for a variety of reasons: to identify candidate TF combinations defining a cell type or cell lineage^9^; to predict genomic binding sites of these TFs and gain insight into their cis-regulatory logic; to predict downstream batteries of differentiation genes and reveal their phenotypic consequences; to study gene regulatory mechanisms such as activation, repression, and feedback; or to study changes in any of these phenomena during development, evolution, or disease. A classical GRN contains by definition the ‘upstream’ TFs, their genomic binding sites, and their target genes^1,88^. However, only few GRNs have been characterized to the level of detail where they include cis-regulatory elements as nodes, so that the TF is linked to its target gene via its binding site in a regulatory region^3,22,42^.

That we lack such GRNs is mainly due to the challenges associated with the experimental identification and validation of TF binding sites in high-throughput and at genome-wide scale; for example, due the requirement for large amounts of homogenous input material, of high-quality antibodies, in the case of ChIP-seq, and of low-throughput genetic perturbations. Computational methods to predict GRNs have therefore often relied on gene expression data (bulk or single-cell), to search for putative causal relationships between a TF and target genes based on their expression correlation^18^. With our SCENIC method^19^ we made a first attempt to bring back cis-regulatory support for such correlation-based TF-gene interactions, basically by pruning co-expression edges to retain only the interactions that also show evidence for the correct TF motif. Next, a significant leap occurred, namely to predict high-confidence genome-wide TF binding sites by combining sequence analysis and chromatin accessibility profiling^89^. Bulk DHS atlases lead to TF foot-printing and regulatory grammars^90^; while scATAC-seq combined with sequence analysis allowed for TF binding site predictions across tissue atlases^91^, reaching similar levels of recall and precision compared to ChIP-seq; this approach was recently coined as ‘computational ChIP’^23^. Not surprisingly, this strategy to predict TF binding sites raised high expectations for eGRN mapping, particularly when it is combined with co-expression network inference. We and others have investigated different ways to bring all these data layers together, and as first proof-of-principle we spatially mapped a comprehensive eGRN (combining scATAC-seq, scRNA-seq, and sequence analysis) for Drosophila retinal development^22^, followed by brain-wide eGRNs for more than 40 cell types^3^.

In the current work we comprehensively analyzed and improved computational strategies to map eGRNs from single cell multiome data (or from unpaired scATAC and scRNA), with a strong basis in regulatory sequence analysis. We implemented a new pipeline, optimized the performance of each step in the pipeline, and compared each step with alternative approaches. This led to a modular software suite, called SCENIC+, that we make publicly available, alongside example data sets and tutorials. A first improvement is the analysis of scATAC-seq data, for which we re-implemented cisTopic, now allowing for topic modeling that scales to hundreds of thousands of cells, while maintaining its high accuracy compared to alternative scATAC-seq analysis methods^92–95^. Topic modelling also provides imputation to resolve drop-outs, which we found to result in more accurate region-gene links in comparison with other methods that use raw binary data.

Another novelty in SCENIC+ is a new Python package called pycisTarget, that performs sequence analysis and identifies candidate TF motifs per cell type. pycisTarget re-addresses the classical problem of TF motif enrichment, which has seen gradual improvements over the last 20 years, particularly because of the increase in high-quality position weight matrices (PWM) for many TFs. However, the increase in PWMs also poses challenges of redundancy. We created, to our knowledge, the largest motif collection to date, with more than 40,000 PWMs (32,000 unique motifs), corresponding to ∼1,5K human TFs, which represents an average increase of 48% compared to other motif collections. To address motif redundancy, we found that maintaining the diversity of motifs achieves a significantly higher accuracy compared to defining one archetype motif per TF. The use of motif discovery in eGRN inference is unique to SCENIC+; other methods rely on motif scanning to identify TFBSs, but this comes with the risk of high false positive prediction rates^96,97^. Technically, pycisTarget provides three different motif discovery methods, including the widely used Homer^30^, although the cisTarget approach with motif clustering showed the highest TF recall, while maintaining high precision.

We created a new benchmark dataset to compare eGRN inference methods using the ENCODE core cell lines and foresee that future methods can be easily benchmarked on the same data. We compared SCENIC+ with other methods, including CellOracle^39^, FigR^41^, GRaNIE^42^, and Pando^40^. All these methods solve certain aspects of the eGRN inference problem and focus on particular sub-problems. For example, CellOracle is excellent at predicting broadly active TFs and focuses on TF-Gene networks during differentiation, exploiting trajectories and predicting the consequence of TF perturbations, while it focuses less on TF binding sites and enhancers^39^. FigR proposes a new concept of co-regulated chromatin hubs, called TF-DORCs (Domains of Regulatory Chromatin), thereby focusing more on chromatin interactions rather than individual TF binding sites and enhancers^41^. Pando^40^ and GRaNIE^42^ have more similar goals compared to SCENIC+, but GRaNIE was originally designed for bulk RNA/ ATAC data (although we have shown here that for small single-cell data sets it performs accurately). We believe that having multiple methods available to solve eGRN mapping is an advantage for the community, and since each method has its own strengths, their combination may yield further improvements. For example, inspired by CellOracle, we implemented a regression model to predict gene expression from their TF inputs, which allows to predict the effect of a TF perturbation on the transcriptome; and consequently, this technique allows identifying the most ‘potent’ TFs for each cell state.

We applied SCENIC+ to a series of biological systems, in Drosophila, mouse and human, covering retinal development, immune cell diversity, cancer cell heterogeneity, and neuronal diversity in the brain. These applications covered continuous and steady-state data sets, of both large and small sizes (ranging from 84K to 1,7K cells). Across all the systems we studied, SCENIC+ revealed high-confidence master regulators and their targets, many of which are supported by literature, and allowed us to explore TF cooperativity (e.g. SOX10-TFAP2A-MITF in melanoma cell lines and PAX5-EBF1-POU2F2 in B cells).

A particular novelty in our work is the comparative study of eGRNs between the human and mouse cortex, where we exploited the high conservation of cell types between these species^56^ to validate TF, motif, and target gene predictions. For each cell type in the brain, SCENIC+ identifies several TFs that are either uniquely active, or strongly biased to that cell type. Interestingly, while TF combinations and predicted target genes are overall conserved, TF binding sites show high turnover. This finding corroborates earlier, experimental findings based on DNase I footprinting across species^61^.

Another novelty we present is the spatial mapping of eGRNs onto tissues, making use of newly generated and previously published spatial transcriptomics data. We found that both sequencing-based methods, such as 10x Visium, and probe-based imaging methods, in our case Molecular Cartography, provide a good basis to study spatial GRN activities. This proof-of-concept may be exploited in the future for cross-species analysis, or even for further benchmarking challenges. Using cross-species conservation and spatial activity, we thus confirmed the quality of the eGRNs inferred by SCENIC+.

Two other new approaches we present to SCENIC+ users are the use of eGRNs to predict trajectories (both linear and branched), and perturbations. Using the cell-intrinsic properties of TF-enhancer and enhancer-gene relationships provides a regulatory scaffold (or model) to predict the future state in a dynamic process, and to predict the future state after a TF knockout. We foresee that the application of SCENIC+ to single-cell perturbation studies (e.g., CROP-seq) with multiome readout, as will likely appear in the future, will be a powerful combination. By applying such simulations for all TF knockouts *in silico*, we found that well known driver TFs can be prioritized to modulate melanoma phenotype switching, followed by several new promising candidates^49,98^ .

There are still several challenges that remain to be solved to improve eGRN prediction accuracies. One of the current caveats is the use of thresholds at different steps in all methods including SCENIC+. We envision that the integration of sequence-based convolutional neural networks with ATAC/RNA auto-encoders could circumvent the ‘heuristics’ that current methods use. Another caveat is that cell type identity is often controlled by reciprocal activation and repression of different TFs from the same family. A classic example is the duality provided by SOXE family members SOX10 and SOX9 in melanoma, which are expressed dis-jointly, but have the same TF recognition sequence. Correlations and anti-correlations between motif enrichment, chromatin accessibility, and gene expression can lead to alternative interpretations of activation by one TF, and repression by the other, and such predictions are impossible to disentangle without experimental TF binding data (Supplementary Note 2, Fig S12). Nevertheless, SCENIC+ predicted several well-known repressor TFs as repressor, where the TF is expressed in those cells where its predicted target regions are not accessible, and its predicted target genes are not expressed. These include Ct in the Drosophila eye antennal disc (expressed in outer antennal rings and late photoreceptors) and HES1 in melanoma for example (Supplementary Note 2, Fig S12). As repression is an important feature of GRN biology, these findings are promising, but may require further computational innovations. Currently, the power for detecting negative correlations is intrinsically lower given that it relies on the absence of data which is aggravated by dropouts in single cell data.

In conclusion, we presented a new computational toolkit for the inference of eGRNs called SCENIC+. SCENIC+ is available as an open-source python package at https://github.com/aertslab/scenicplus. We also provide extensive documentation and tutorials at https://scenicplus.readthedocs.io. Results of the analysis presented in this manuscript can be further explored using Scope (https://scope.aertslab.org/#/scenic-v2) and UCSC.

## Methods

### SCENIC+ workflow

The SCENIC+ workflow consists of three main steps, performed with three linked Python modules: 1) unsupervised identification of enhancers with shared accessibility patterns from the scATAC-seq data (pycisTopic), 2) prediction of TF binding sites via motif enrichment analysis (pycisTarget) and 3) prediction of eGRNs combining TF expression, TF binding sites, region accessibility and gene expression (SCENIC+) (Fig 1a, S1). The minimal input for SCENIC+ is a gene expression matrix with cell type annotation and a corresponding scATAC-seq fragments file. The two latter can be replaced by a precompiled matrix with fragment counts and precomputed peak coordinates.

The three Python modules include detailed tutorials to facilitate their use for standalone analyses and within the SCENIC+ pipeline. Links to the tools, SCENIC+ code and tutorials are available at: https://scenicplus.readthedocs.io.

#### PycisTopic

Pycistopic is an improved Python-based version of our Bayesian framework cisTopic^24^, which exploits a topic modelling technique called Latent Dirichlet Allocation (LDA)^99^. This unsupervised approach simultaneously clusters cells and co-accessible regions into regulatory topics, and it is among the top performers among numerous independent single-cell epigenomics benchmarks^92–95^. Outside of the SCENIC+ framework, pycisTopic can be used to analyze independent scATAC-seq data as well. PycisTopic is available at https://github.com/aertslab/pycisTopic, with full documentation and tutorials available at https://pycistopic.readthedocs.io The full pycisTopic pipeline consists of the following steps (*indicates those required for the SCENIC+ workflow, ** indicates those recommended for the SCENIC+ workflow):

- **Consensus peak calling**\*: PycisTopic will first create a set of consensus peaks across all cells by calling and merging peaks on pseudobulk ATAC-seq profiles per cell type. First, utilising the fragments file and barcode-cell type annotations provided by the user we generate pseudobulk fragments bed files and coverage bigwig files per cell type. In a second step, peaks are called in each of these pseudobulks using MACS2, using –format BEDPE (as we are providing fragments bed files as input) and –keep-dup all --shift 73 --ext_size 146, as recommended for scATAC-seq data. To derive a set of consensus peaks, we use the iterative overlap peak merging procedure describe in Corces et al. (2018)^100^. First, each summit is extended a ‘*peak_half_width*’ (by default, 250bp) in each direction and then we iteratively filter out less significant peaks that overlap with a more significant one. During this procedure peaks will be merged and depending on the number of peaks included into them, different processes will happen: 1) 1 peak: The original peak will be kept, 2) 2 peaks: The original peak region with the highest score will be kept and 3) 3 or more peaks: The original region with the most significant score will be taken, and all the original peak regions in this merged peak region that overlap with the significant peak region will be removed. The process is repeated with the next most significant peak (if it was not removed already) until all peaks are processed. This procedure will happen twice, first in each pseudobulk peaks, and after peak score normalization to process all peaks together. We recommend using this set of regions downstream, as we (and others) have observed that using pseudobulk peaks improves signal compared to using bulk peaks across the whole population (specially for rare cell types, whose signal may be confused by noise while using the merged ATAC-seq profile of the whole population)^92^. In case of independent scATAC-seq data, the cell annotation can also be obtained from alternative methods, such as a preliminary clustering analysis using a predefined set of genome-wide regions/peaks (e.g. SCREEN^101^) as input to identify cell populations.
- **QC analysis and cell selection**\*\*: PycisTopic computes QC metrics at the sample-level and the barcode-level. Sample-level statistics can be used to assess the overall quality of the sample, while barcode level statistics can be used to differentiate good quality cells versus the rest. These metrics include: - *Barcode rank plot:* The barcode rank plot shows the distribution of non-duplicate reads and which barcodes were inferred to be associated with cells. A steep drop-off (’knee’) is indicative of good separation between the cell-associated barcodes and the barcodes associated with empty partitions. - *Insertion size:* ATAC-seq requires a proper pair of Tn5 transposase cutting events at the ends of DNA. In the nucleosome-free open chromatin regions, many molecules of Tn5 can kick in and chop the DNA into small pieces; around nucleosome-occupied regions, and Tn5 can only access the linker regions. Therefore, in a good ATAC-seq library, you should expect to see a sharp peak at the <100 bp region (open chromatin), and a peak at ∼200bp region (mono-nucleosome), and other larger peaks (multi-nucleosomes). A clear nucleosome pattern indicates a good quality of the experiment. - *Sample TSS enrichment:* The TSS enrichment calculation is a signal to noise calculation. The reads around a reference set of TSSs are collected to form an aggregate distribution of reads centered on the TSSs and extending to 1000 bp in either direction (for a total of 2000bp). This distribution is then normalized by taking the average read depth in the 100bp at each of the end flanks of the distribution (for a total of 200bp of averaged data) and calculating a fold change at each position over that average read depth. This means that the flanks should start at 1, and if there is high read signal at transcription start sites (highly open regions of the genome) there should be an increase in signal up to a peak in the middle. - *FRiP distribution:* Fraction of all mapped reads that fall into the called peak regions, i.e. usable reads in significantly enriched peaks divided by all usable reads. A low FRIP indicates that many reads form part of the background, and so that the data is noisy. - *Duplication rate:* A fragment is considered “usable” if it uniquely maps to the genome and remains after removing PCR duplicates (defined as two fragments that map to the same genomic position and have the same unique molecular identifier). The duplication rate serves to estimate the amount of usable reads per barcode. High duplication rates may indicate over-sequencing or lack of fragments after transposition and encapsulation. On the other hand, barcode-level statistics can be used to select high-quality cells. Typical measurements that can be used are: - *Total number of (unique) fragments* - *TSS enrichment:* The normalized coverage at the TSS position for each barcode (average +-10bp from the TSS). Noisy cells will have a low TSS enrichment. - *FRiP:* The fraction of reads in peaks for each barcode. Noisy cells have low FRIP values. However, this filter should be used with nuance, as it depends on the quality of the original peaks. For example, if there is a rare population in the sample, its specific peaks may be missed by peak calling algorithms, causing a decrease in their FRIP values.
- **Count matrix generation*:** PycisTopic can generate a fragment count matrix from the fragments files, a set of regions (preferably, consensus regions as previously explained) and a list of high quality cells. Alternatively, a precomputed count matrix can also be used as input. In this step a cisTopic object will be created, including the fragment counts, path/s to the fragments files (if used to generate the count matrix) and cell/region metadata.
- **Doublet identification:** The fragment count matrix can be used as input for Scrublet^102^ (v0.2.3), By default, and when dealing with 10x data sets, we set the expected doublet rate to 10%.
- **Topic modelling algorithms and model selection*:** PycisTopic implements two algorithms for topic modelling, serial LDA with a Collapsed Gibbs Sampler (as implemented in the lda module) and Mallet, which allows to parallelize the LDA model estimation. We use the same default parameters as in cisTopic^24^. In addition, we have included additional metrics for model selection: - *Minmo_2011:* Uses the average model coherence as calculated by Mimno et al (2011)^103^. To reduce the impact of the number of topics, we calculate the average coherence based on the top selected average values. The better the model, the higher coherence. - *Log-likelihood:* Uses the log-likelihood in the last iteration as calculated by Griffiths and Steyvers (2004)^104^, as used in cisTopic. The better the model, the higher the log-likelihood. - *Arun_2010:* Uses a density-based metric as in Arun et al (2010)^105^ using the topic-region distribution, the cell-topic distribution and the cell coverage. The better the model, the lower the metric. - *Cao_Juan_2009:* Uses a divergence-based metric as in Cao Juan et al (2009)^106^ using the topic-region distribution. The better the model, the lower the metric.
- **Dimensionality reduction and batch effect correction**:** We can cluster the cells (or regions) using the leiden algorithm and perform dimensionality reduction with UMAP and TSNE using the cell-topic (or topic region distributions). In addition, harmonypy (v0.0.5) can be used on scaled cell-topic distributions to correct for batch effect between samples (see mouse cortex analysis). When working with single-cell multiome data, it is possible to co-cluster and reduce dimensionality using both the scRNA-seq and scATAC-seq data by using UMAP to build fuzzy simplicial sets (similar to KNN graphs).
- **Topic binarization and QC*:** To perform motif analysis (and other downstream steps) we need to have topics as region sets rather than continuous distributions. We have included several binarization methods (applicable for topic-region and cell-topic distributions): ’*otsu*’ (Otsu, 1979)^107^, ’*yen*’ (Yen et al., 1995)^108^, ’*li*’ (Li & Lee, 1993)^109^, ’*aucell*’ (Van de Sande et al., 2020)^20^ or ’*ntop*’ (Taking the top n regions per topic). Otsu and Yen’s methods work well in topic-region distributions; however, for some downstream analyses it may be convenient to use ’*ntop*’ to have balanced region sets. By default, pycisTopic uses Otsu for binarization. For cell-topic distributions, we recommend using the AUCell method. In addition, pycisTopic includes new metrics to assess topic quality: - *Number of assignments and regions/cells per binarized topic*. - *Topic coherence (Mimno et al., 2011)*^103^: Measures to which extent high scoring regions in the topic are co-accessible in the original data. If it is low, it indicates that the topic is rather random. The higher, the better is a topic. - *The marginal topic distribution:* Indicates how much each topic contributes to the model. The higher, the better is a topic. - *The gini index:* Value between 0 and 1, that indicates the specificity of topics (0: General, 1: Specific)
- **Drop-out imputation*:** Thanks to the probabilistic nature of topic modelling, drop-outs can be imputed by multiplying the cell-topic and topic-region distributions, resulting in a matrix with the probabilities of each region in each cell. This approach was already available in cisTopic and has been extensively validated in external works^92–95^.
- **Calculation of Differentially Accessible Regions (DARs)*:** Using the imputed fragment matrix we can identify Differentially Accessible Regions, or DARs, between cell types. Briefly, we perform a Wilcoxon rank-sum test between each group in the specified variable and the rest. Alternatively, specified contrast can be provided as a list with foreground and background groups. By default, we identify a region as a DAR if padj < 0.05 and LogFC > 0.5.
- **Gene activity and Differentially Accessible Genes (DAGs):** The gene activity recapitulates the overall accessibility values in a space around the gene. Differentially Accessible Genes (DAGs) can be derived from this matrix. The user can select among different options: - *Search space:* The user can choose whether the search space should include other genes or not (*use_gene_boundaries*), and the minimum and maximum distance it should have (upstream and downstream). Promoters can be excluded from the calculations, as they are usually ubiquitously accessible. - *Distance weight:* The parameters related to the distance weight measure the impact of distance when inferring region to gene weights as an exponential function. The user can control whether this weight should be used (*distance_weight*) and the effect of the distance (*decay_rate*). In addition, the user can choose from which distance to the gene body this weight i s applied (*extend_gene_body_upstream* and *extend_gene_body_downsstream*) - *Gene size weight:* Large genes may have more peaks by chance. The user can optionally apply a weight based on the size of each gene (*gene_size_weight*), which by default is dividing the size of each gene by the median gene size in the genome. Alternatively, the user can also use *average_scores* which will calculate the gene activity as the mean weighted region accessibility of all regions linked to the gene. - *Gini weight* : This weight will give more importance to specifically accessible regions (*gini_weight*).
- **Label transfer:** PycisTopic includes wrappers for several label transfer methods using annotated scRNA-seq and the gene activity matrix. The methods available for label transferring are: ‘*ingest*’^110^, ’*harmony*’^111^, ’*bbknn*’^112^, ’*scanorama*’^113^ and ’*cca*’. Except for ingest, these methods return a common co-embedding and labels are inferred using the distances between query and reference cells as weights.
- **pyGREAT:** pycisTopic makes GREAT (Genomic Regions Enrichment of Annotations Tool)^114^ analysis automatic by constructing a HTTP POST request according to an input region set and automatically retrieving results from the GREAT web server, analogously to the rGREAT package in R.
- **Signature enrichment:** Given epigenomic signatures are intersected with the regulatory regions in the dataset and summarized into region sets. Using the imputed fragment matrix, all regions in each cell are ranked and the cell-specific rankings are used as input for AUCell. By default, we use 5% of the total number of regions in the dataset as a threshold to calculate the AUC.
- **Export to loom files**:** PycisTopic allows to export cisTopic object to loom files compatible with Scope for visualization^32^ and SCopeLoomR, for importing pycisTopic analyses into R.

#### PycisTarget

PycisTopic unsupervisedly identifies groups of co-accessible regions (cis-regulatory topics) as well as cell type specific enhancers (DARs). These regulatory programs are used in the second step of SCENIC+, in which TFs and their potential target regions (i.e. cistromes) are identified using motif enrichment analysis. For this purpose, we have developed pycisTarget, a motif enrichment suite that combines different motif enrichment approaches such as cisTarget^27–29^ and Homer^30^; and a novel approach to compute Differentially Enriched Motifs between sets of regions called DEM. Pycistarget is available at https://github.com/aertslab/pycistarget, with full documentation and tutorials available at https://pycistarget.readthedocs.io.

- **Homer:** PycisTarget includes a wrapper to run Homer’s *findMotifsGenome.pl*, allowing the identification of known and *de novo* motifs (by default using default Homer parameters). For identifying cistromes for each motif, found motifs are used as input for homer2 find. Known motifs are annotated according to the motifs annotation in the SCENIC+ motif collection. To annotate *de novo* motifs, tomtom is run with the SCENIC+ motif collection to identify the closest match, allowing to transfer its annotation to the *de novo* motif when specified. To form TF cistromes, motif-based cistromes are combined based on the TF annotations.
- **Generation of cisTarget databases:** cisTarget and DEM require ranking and score-based databases, respectively, with regions as rows, motifs (or motif clusters) as columns, and motif enrichment scores, or ranking values of these scores, as values. For this, we have developed an adapted version of Cluster-Buster^115^, which is now 2 times faster. Cluster-Buster uses Hidden Markov Models (HMMs) to score clusters of motifs (i.e., Cis-Regulatory Modules (CRM)) given a set of regions. This implementation is available at https://resources.aertslab.org/cistarget/programs/cbust. Briefly, given a motif collection (in cb format) and a set of regions, we run Cluster-Buster using each motif across all regions. When dealing with motif clusters, Cluster-Buster uses all motif variations by implanting each motif as a hidden state in a HMM, and each candidate sequence receives a log-likelihood ratio (LLR) score per motif cluster (i.e. CRM score). A scores database is first generated by taking the highest CRM score per sequence. A ranking database is then generated by ranking, for each motif, all the regions by decreasing motif score. The code and documentation to generate these databases is available at https://github.com/aertslab/create_cisTarget_databases. In addition, we provide precomputed databases using predefined sets of regulatory regions for hg38, mm10 (using SCREEN regions) and dm6 (using cisTarget regions, defined by partitioning the entire non-coding Drosophila genome based on cross-species conservation) at: https://resources.aertslab.org/cistarget/databases/https://resources.aertslab.org/cistarget/databases/.
- **cisTarget:** pycisTarget implements our ranking based motif enrichment method cisTarget^27–29^. Briefly, we first score genomic regions (i.e. consensus peaks, or predefined regions from SCREEN) using a motif collection with Cluster-Buster, as previously described. These regions will be ranked based on their motif score for each motif. The input regions are intersected with regions in the database (with at least 40% overlap). cisTarget uses a recovery curve approach (for each motif), in which a step is taken in the y-axis when as region in the motif ranking (x-axis) is found in the region set. The Area Under the Curve for each motif is normalized based on the average AUC for all motifs and their standard deviation, resulting in a Normalized Enrichment Score (NES) that is used to quantify the enrichment of a motif in a set of regions:

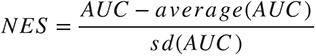

By default, motif that obtain a NES above 3.0 are kept. To obtain the target regions for each TF, the motif-based cistromes of motifs annotated to that TF are combined.
- **DEM:** DEM performs a Wilcoxon test between scores in foreground and background region sets to assess motif enrichment. Briefly, we first score genomic regions (i.e. consensus peaks, or predefined regions from SCREEN101) using a motif collection with Cluster-Buster, as previously described. Regions in the input region sets are intersected with regions in the database (with at least 40% overlap), and the Wilcoxon test is performed between CRM score distributions in the two groups. By default, motifs with adjusted p-val < 0.05 (Bonferroni) and Log2FC > 0.5 are kept. A cistrome for each motif is generated by simultaneously optimizing precision and recall to separate foreground from background regions or using a predefined CRM threshold. To obtain the target regions for each TF, the motif-based cistromes of motifs annotated to that TF are combined.

Within the SCENIC+ workflow, motif enrichment is performed by default in binarized topics and DARs calculated by pycisTopic, using cisTarget and DEM (including and excluding promoters from the region sets). By default, DEM is run using topics or DARs as foreground and 500 regions in other topics/DARs as background (with the same proportion of promoters as in the foreground). Additional region sets (e.g. DARs derived from specific contrasts instead of using all populations as background) can be easily added. Cistromes derived from all the motif enrichment analysis are merged by TF to generate a final set of TF-region cistromes.

#### SCENIC+

After having identified potential enhancers using pycisTopic and their potential TF inputs using pycisTarget eGRNs can be predicted in the final step of SCENIC+. This step requires as input: the gene expression matrix, the imputed accessibility matrix (from pycisTopic) and the TF cistromes (from pycisTarget). Input data can be single-cell multiome or paired scRNA-seq and scATAC-seq in matching populations (see below). This final step consists of the following steps:

- **Generating pseudo multiome data (in case of non-multiome data):** To generate pseudo multiome data, cells must be annotated by common labels in both data modalities (single-cell chromatin accessibility and gene expression). Pseudo multiome cells are generated by sampling a predefined number of cells from each data modality, within the same cell type annotation label, and averaging the raw gene expression and imputed chromatin accessibility data across these cells to create a multiome meta cell containing data of both modalities. By default, the number of meta cells generated for each cell type annotation label is set as such that each cell is included in a meta cell on average twice.
- **Calculating TF-to-gene and region-to-gene relationships:** TF-to-gene relationships are calculated as described in^19,20^. Briefly, the Arboreto python package (v0.1.6) is used to calculate TF-to-gene importance scores for each TF and each gene, given a list of known TFs and the raw gene expression matrix. By default, Gradient Boosting Machine regression is used. Pearson correlation is used to separate positively correlating from negatively correlating relationships (resp. correlation coefficient above 0.03 or below –0.03 by default). To infer autoregulation the TF-to-TF importance is set to the maximum importance score across genes for that TF added with the value of 1E-5. Region-to-gene relationships are calculated for each gene by considering all regions within a search space surrounding that gene. By default, a search space of a minimum of 1kb and a maximum of 150kb upstream/downstream of the start/end of the gene or the promoter of the nearest upstream/ downstream gene. By default, the promoter of the gene is considered as the TSS of the gene +/- 10 bps. For each consensus peak in the search space of each gene region-to-gene importance scores are calculated using the Arboreto python package (v0.1.6) using the imputed accessibility of the regions as predictors for the raw gene expression counts and Gradient Boosting Machine regression (by default). Spearman rank correlation (SciPy v1.8.1) is used to separate positively correlating from negatively correlating relationships (resp. correlation coefficient above 0.03 or below –0.03 by default). Before eGRN building region-to-gene relationships are binarized using multiple methods. By default, the 85th, the 90th and the 95th quantile of the region-to-gene importance scores, the top 5, 10, and 15 regions per gene based on the region-to-gene importance scores and a custom implementation of the BASC^116^ method on the region-to-gene importance scores is used.
- **eGRN compilation:** First TF-to-region-to-gene modules are generated by intersecting the binarized region-to-gene relationships with TF-to-region cistromes by region name. Next these modules are pruned by running GSEA, *gsea_compute* function from the GSEApy python package (v0.10.8), using the TF-to-gene importance scores, for all TFs and genes in the module, as ranking and genes of the module as gene set. The leading-edge genes along with the binarized regions linked to those genes are retained, generating eRegulons. This analysis is run separately for TF-to-gene and region-to-gene relationships with positive and negative correlation coefficients, for a total of four GSEA runs. By default, eRegulons with less than 10 predicted target genes or obtained from region-to-gene relationships with a negative correlation coefficient are discarded.
- **eRegulon enrichment:** Target regions and genes of each eRegulon are used separately together with regions and genes ranked based on imputed accessibility and raw gene expression counts in each cell as input for AUCell^20^, using the ctxcore python package (v. 0.1.2.dev2+g1ffcf0f). By default, 5% of the total number of regions or genes in the dataset are used as threshold to calculate AUC values. High quality regulons are then selected based on the correlation between region based and gene based AUC values (by default 0.4) and/or AUC values and TF expression.
- **eRegulon dimensionality reduction:** The eRegulon enrichment scores for regions and genes are normalized for each cell and used as input into the UMAP, tSNE or PCA from the python package umap (v0.5.2), fitsne (v1.2.1) or sklearn (v0.24.2) respectively.
- **eRegulon specificity scores:** eRegulon specificity scores are calculated using the RSS algorithm as described in^20,117^ using target region or target gene eRegulon enrichment scores as input.
- **Export to loom files:** To visualize SCENIC+ results in SCope^32^ a chromatin accessibility and gene expression based loom file containing the eRegulons with target regions/genes and eRegulon enrichment scores for target regions/genes is generated, using the LoomXpy python package (v0.4.1; https://github.com/aertslab/LoomXpy). In addition, loom files can also be used to import data into R via the SCopeLoomR package.
- **Visualization in the UCSC genome browser:** To visualize SCENIC+ results in the UCSC genome browser a UCSC interaction file, containing region-to-gene links color coded by region-to-gene importance scores or correlation coefficients, and a bed file, containing genomic coordinates of eRegulon target regions is generated. The UCSC interact file and the bed file are converted to the bigInteract and bigBed format using the bedtobigbed program (v2.7) from UCSC. These can be uploaded to the UCSC genome browser along with pseudobulk bigWig files.
- **Network visualization:** Enhancer GRNs can be visualized using networkx (v2.7.1) concentrical and Kamada-Kawai layouts, with customized features for nodes (size, shape, color, transparency) and edges (stroke, color, transparency). Figures can be generated with networkx (v2.7.1) or interactively with pyvis (v0.1.3.1). In addition, SCENIC+ also allows to export networks to Cytoscape (v3.9.0). The SCENIC+ style for Cytoscape is available at: https://github.com/aertslab/scenicplus/tree/main/cytoscape_styles.

### SCENIC+ motif collection and motif enrichment benchmark

The SCENIC+ motif collection includes more than 49,504 motifs from 29 motif collections (Table S1), with curated TF motif annotations based on direct evidence and orthology between species for human, mouse and fly. In order to account for motif redundancy (i.e. the same or a very similar version of the same motif can be found in more than one of these collections), we implemented a new approach to create non-redundant (or clustered) motif collections using a two-step clustering strategy. First, identical PWMs across collection (after rescaling) were merged, resulting in 34,524 motifs. A matrix with motif-to-motif similarity values was computed using Tomtom (MEME v5.4.1)^118^, and motifs with equal length and q-value < 10-40 were merged, resulting in 32,766 motifs (unclustered motif collection). For clustering, we used motifs that are similar to at least another motif with q-value < 10-5 (11,526), while the remaining were kept as unique motifs, or singlets (9,685). Dimer motifs (1,265) were excluded from the clustering, as well as motifs from factorbook and desso, as they do not have direct annotations since they are derived from AI models. Motifs with an Information Content below 5 were also excluded. We converted the motif similarity value matrix to -log10(Tomtom similiarity)+10-45. We then used Seurat (v4.0.3) to normalize, scale and perform PCA in this matrix. Using 100 PCs, we performed Leiden clustering with resolution 25, resulting in 199 clusters. Clusters were refined by running STAMP (v1.3; using the -cc -sd –chp option)^119^ resulting in 1,985 clusters. For each cluster, we use STAMP’s consensus motif to generate the motif logo. The TF annotation of a cluster was inferred by merging the TF annotations (direct and orthology) of all its members. Overall, the clustered motif collection contains 9,685 singlets, 1,265 dimers and 1,985 clusters (with a mean of 5.8 motifs per cluster).

The SCENIC+ motif collection contains 8,384, 8,045, 958 annotated clusters for 1,553, 1,357 and 467 TFs (with an average of 5, 6, 2 motifs per TF) for human, mouse, and fly; respectively. Importantly, motifs are not only annotated based on direct TF-motif experimental evidence; but also based on orthology (by downloading gene orthologs for the selected species from Ensembl Biomart 105), which permits the incorporation of annotations based on experiments in different species. In fact, 433 mouse TFs are only found via orthology, augmenting TF-annotations by 47%, as more experiments have been performed in human systems than in mouse.

To benchmark the different motif enrichment techniques included in pycisTarget and approaches to build databases, we assessed the recovery of target TFs using 309 ChIP-seq data sets from the Deeply Profiled Cell Lines collection from ENCODE^12,13^ (Table S2). These tracks have been also included in Unibind^10,11^ and present the TF motif, showing that their quality is good enough to find the target TF motif back. Motif enrichment was performed using Homer, cisTarget and DEM. For the latter two, three different approaches for creating the motif databases were used: 1) generating a database without clustering the motif collection and only retaining annotated motifs (24,309 motifs), 2) generating a database using a single consensus motif (or archetype) for each of the STAMP clusters and 3) generating a database by scoring regions using all the motifs in a cluster (as described above). Our results show that using cisTarget (or DEM) with the clustered collection (using all PWMs for scoring) recovers the most target TF motifs and in the top positions, followed by Homer. Note that DEM is meant to be used to compare foreground and background region sets, and in this analysis, we used regions in other tracks as background. We found that scoring with archetypes performs worse compared to using all PWMs in the cluster, especially for clusters where there is more variability. In addition, since our motif collection contains licensed motifs from Transfac Pro, we also benchmarked cisTarget and DEM using a publicly shareable clustered collection (using all PWMs for scoring but removing these protected motifs, finding equal TF recovery and comparable scores in regions). Finally, we also compared the target regions (or cistromes) recovered by each approach with those reported by Unibind^10,11^ as ground truth, finding that both cisTarget and DEM have higher precision and recall compared to Homer (Fig S3).

We provide all PWMs and clusters in the SCENIC+ motif collection for the creation of custom databases (with the exemption of the licensed Transfac Pro PWMs) at https://motifcollections.aertslab.org/ precomputed databases using genomic regions (SCREEN for mouse and human, and cisTarget regions for fly) and updated gene-based databases for SCENIC at https://resources.aertslab.org/cistarget/databases/.

### Comparison of cisTopic and pycisTopic

To assess the improvements on the topic modelling strategy, we benchmarked cisTopic and pycisTopic using a simulated single cell epigenomics data set from 5 melanoma cell lines (3 melanocytic and 2 mesenchymal) with 100 cells^24^, downloaded from https://github.com/aertslab/cisTopic. We ran cisTopic (v2.1.0) using Collapsed Gibbs Sampling and WarpLDA modelling, and pycisTopic (v1.0.1.dev21+g8aa75d8) using Collapsed Gibbs Sampling and Mallet, using 150 iterations and 21 cores for 21 models (starting from 2 topics and from 5 to 100 increasing by 5). For all models we set alpha to 50/number of topics and beta to 0.1, as previously described2^4,104^.

### PBMC analysis

#### Data

Filtered feature barcode matrix and fragments files were downloaded from the 10x website (https://cf.10xgenomics.com/samples/cell-arc/1.0.0/pbmcgranulocyte_sorted_10k/pbmc_granulocyte_sorted_10k_filtered_feature_bc_matrix.h5 and https://cf.10xgenomics.com/samples/cell-arc/1.0.0/pbmc_granulocyte_sorted_10k/pbmc_granulocyte_sorted_10k_atac_fragments.tsv.gz).

#### Quality control of scRNA-seq and cell type annotation

The scRNA-seq part of the multi-ome dataset was preprocessed using Scanpy (v1.8.2)^110^. Briefly, genes expressed in less than 3 cells were removed. Cells which expressed less than 200 genes, more than 6,000 genes or had more than 30 % counts in mitochondrial genes were removed. Doublets were detected and removed using Scrublet (v0.2.3)^102^ with a doublet score threshold of 0.17. This resulted in 11,101 high quality cells. Cells were annotated using ingest label transfer, using the *sc.tl.ingest* function included in Scanpy (v1.8.2)^110^, and by matching transferred labels to leiden clusters (resolution 0.8) based on maximum overlap using the annotated pbmc dataset included in the Scanpy package as a reference (*sc.datasets.pbmc3k_processed()*).

#### Quality control of scATAC-seq and topic modeling

The scATAC-seq part of the multiome dataset was preprocessed using pycisTopic (v1.0.1.dev21+g8aa75d8). Briefly, consensus peaks (342,044) were called as described above using the downloaded fragments file and cell type labels from the scRNA-seq side. Cells with less than 1E33 total number of unique fragments, FRIP below 0.45 and TSS enrichment below 5 were removed. Doublets were detected and removed using Scrublet (v0.2.3)^102^ with a doublet score threshold of 0.33. This resulted in 10,955 high quality cells. Topic modeling was performed as described above using LDA with the collapsed Gibbs sampler. A model of 16 topics was selected based on the stabilization of *Arun_2010*, *Cao_Juan_2009*, *Minmo_2011* and *loglikelihood* quality metrics.

#### Motif enrichment analysis

Motif enrichment analysis was performed using pycisTarget (v1.0.1.dev17+gd2571bf) as described above. For this, a custom score and ranking database was generated using *create_cisTarget_databases* python package using the DNA sequences of consensus peaks and all annotated motifs as input. Motif enrichment was performed using both the cisTarget and DEM algorithm on cell type based DARs (log2 fold change above 1.5), top 3,000 regions per topic and topics binarized using the otsu method. The motif enrichment analysis was run both once including promoters and excluding them. Promoters were defined as regions within 500 bp up- or downstream of the TSS of each gene. TSS’s for each gene were downloaded from Biomart (http://sep2019.archive.ensembl.org), using the pybiomart package (v0.2.0).

#### SCENIC+ analysis

The raw gene expression count matrix, imputed accessibility (product of topic-region and cell-topic probabilities), and motif enrichment results were used as input into the SCENIC+ workflow, keeping 9,409 cells having both high quality ATAC-seq and RNA-seq profiles. The SCENIC+ workflow was run using default parameters and as described above. Briefly, a search space of a maximum between either the boundary of the closest gene or 150 kb and a minimum of 1 kb upstream of the TSS or downstream of the end of the gene was considered for calculating region-to-gene relationships using gradient boosting machine regression. TF-to-gene relationships were calculated using gradient boosting machine regression between all TFs and all genes. Genes were considered as TFs if they were included in the TF list available on http://humantfs.ccbr.utoronto.ca/. 120. Final eRegulons were constructed using the GSEA approach in which region-to-gene relationships were binarized based on gradient boosting machine regression importance scores using the 85th, 90th and 95th quantile; the top 5, 10 and 15 regions per gene and using the BASC method^116^ for binarization. Only eRegulons with a minimum of 10 target genes were retained. For each eRegulon cellular enrichment scores (AUC) of target genes and regions were calculated using the AUCell algorithm^20^. eRegulons for which the correlation coefficient between semi pseudo bulked per cell type (100 meta cells per cell type and 5 cells per meta cell) TF expression and region AUC scores was above 0.7 or below –0.8 were considered as high quality and used for downstream analysis. This resulted in 63 regulons, with a median of 296 genes and 528 regions per regulon. Analyses can be explored in Scope at https://scope.aertslab.org/#/scenic-v2 and UCSC at https://genome-euro.ucsc.edu/s/Seppe%20De%20Winter/scenicplus_pbmc.

#### ChIP-seq enrichment in eRegulon target regions

PAX5, EBF1 and POU2F2 ChIP-seq BigWig and summit bed files were downloaded from Encode (https://www.encodeproject.org/)^12,13^ using the following accession numbers ENCFF702MTT and ENCSR000BHD for PAX5; ENCFF107LDM and ENCSR000BGU for EBF1; ENCFF025CFR and ENCFF922DNK for POU2F2 for BigWig and summit bed files respectively. The target regions of PAX5(+), EBF1(+) and POU2AF1(+) and regions targeted by any combination of two eRegulons were intersected with the ChIP-seq summits, the original region was kept if a region did not intersect with the summit. ChIP-seq coverage was calculated on these regions using the PyBigWig package (v0.3.18) using 50 bins. Coverage was min-max normalized by using the minimum and maximum across all regions per ChIP-seq dataset. ChIP-seq data along with pseudo-bulk accessibility was visualized using the Signac R package (v1.3.0)^121^.

### Benchmark of GRN inference methods

#### Simulated single-cell multiomics data

Data from ENCODE Deeply Profiled Cell Lines was downloaded from https://www.encodeproject.org/^12,13^. Single cell multiome data was generated using bulk RNA-seq and ATAC-seq data from 8 ENCODE Deeply Profiled Cell Lines, namely MCF7 (breast cancer, ENCFF136ANW and ENCFF772EFK, for RNA-seq and ATAC-seq, respectively), HepG2 (hepatocellular carcinoma, ENCFF660EXG and ENCFF239RGZ), PC3 (prostate cancer, ENCFF874CFD and ENCFF516GDK), GM12878 (B cell, ENCFF626GVO and ENCFF415FEC), K562 (leukemia, ENCFF833WFD and ENCFF512VEZ), Panc1 (pancreatic cancer, ENCFF602HCV and ENCFF836WDC), IMR90 (lung fibroblast, ENCFF027FUC and ENCFF848XMR) and HCT116 (colon cancer, ENCFF766TYC and ENCFF724QHH). Briefly, we simulated 500 single-cell multiomics profiles by randomly sampling 50,000 reads and 20,000 fragments from each bulk RNA-seq and ATAC-seq profiles, respectively; resulting in a data set with 4,000 simulated single-cell cells. The scRNA-seq count matrix was generated using featureCounts (Subread v1.6.3) and the GRCh38.86 genome annotation. After calling peaks with MACS2 (v2.1.2.1) on the bulk ATAC-seq samples, we generated a set of 642,982 consensus peaks that was used to generate the scATAC-seq matrix, as previously described. Analyses can be explored in Scope at https://scope.aertslab.org/#/scenic-v2 and UCSC at https://genome.ucsc.edu/s/cbravo/SCENIC%2B_DPCL.

#### Methods

The simulated data set was analyzed with different state-of-the-art methods. For all methods we used a search space of 150kb for inferring region-gene relationships and kept regulons with at least 25 target genes:

- **SCENIC+:** pycisTopic was run with default parameters, using Mallet with 500 iterations for topic modelling and generating 21 topics (2 topics and from 5 to 100 increasing by 5), selecting the model with 40 topics based on the model selection metrics. To assess the effect of the regions in the motif database, we ran SCENIC+ using consensus peaks (642,982 peaks) and SCREEN regions^101^, that overlap with ∼80% of the consensus peaks in this case, finding that results between the variants for each method give similar results. SCENIC+ was run with default parameters, using http://oct2016.archive.ensembl.org/ as Biomart host. High quality regulons were selected based on the correlation between gene-based regulon AUC and region-based regulon AUC (> 0.7). This resulted in 178 regulons, with a median of 437 genes and 774 regions per regulon.
- **CellOracle**^39^: CellOracle was run as described at https://morris-lab.github.io/CellOracle.documentation/. Briefly, scRNA-seq data was analyzed using Scanpy (v1.8.2), using *flavor=‘cell_ranger’* and *n_top_genes=3000* to identify highly variable genes, and 7 PCs for dimensionality reduction. scATAC-seq data was processed with Cicero (v1.6.2) with *window=150000*, which inferred 13,123 region-gene connections with coaccess >= 0.8. Next, gimmemotifs (v0.17.1) was used to identify TFBS using the gimmemotifs motif collection with default parameters. After inferring the links, we filtered links with the *filter_links* function using p=0.001, *weight=’coef_abs’*, *threshold_number=2000*. This resulted in 157 regulons, with a median of 49 genes and 50 regions per regulon.
- **Pando**^40^: Pando was run as described at https://quadbiolab.github.io/Pando/. Briefly, the data was processed using Signac (v1.3.0) and SCREEN regions were used in the *initiate_grn* function. MotifMatchR (v1.10.0) was used with motifs from Jaspar and CisBP. To infer the grn (*infer_grn*), we used *peak_to_gene_method=’Signac*’, *method=’glm’*, *upstream=150000*, *downstream=150000*. This resulted in 235 regulons, with a median of 43 genes and 42 regions per regulon.
- **FigR**^41^: FigR was run as described at https://buenrostrolab.github.io/FigR/. Briefly, we used the pycisTopic cell-topic matrix (with 40 topics, as derived in the SCENIC+ workflow) to infer the kNN matrix for smoothing the data. We kept peak-gene correlations with p-value < 0.05 and kept TF-DORC associations with abs(score) > 1, resulting in 10,757 TF-gene pairs. This resulted in 71 regulons, with a median of 39 genes per regulon.
- **GRaNIE**^42^: GRaNIE was run as described at https://grp-zaugg.embl-community.io/GRaNIE/, initially using the bulk data. However, its performance was very poor (likely to the reduced size of the data set), only finding 26 TFs and 11,106 TF-region-gene links. Interestingly, when we applied it to our simulated single-cell data set (with adaptations), its recovery increased (finding 39 TFs and 44,666 TF-region-gene links); hence, we report the latter results. Briefly, we used the binary matrix (with a pseudocount of 1) as input, normalizing the scATAC-seq data using “*Deseq_sizeFactor*” and the scRNA-seq data with “*quantile*”. Motif scanning was performed using the default motif collection (Hocomoco). In *addConnections_peak_gene* we used *promoterRange=150000*. To filter the GRN, we used 0.2 as fdr threshold. This resulted in 39 regulons, with a median of 176 genes and 68 regions per regulon.
- **SCENIC**^20^: SCENIC was run as described at https://pyscenic.readthedocs.io/en/latest/. To assess the effect of clustering in the motif collection, we have also benchmarked SCENIC using the non-clustered and the clustered databases, and both at the same time, obtaining similar results. For each collection, we generated two gene-motif rankings (10kb around the TSS or 500bp upstream the TSS). Using both collections at the same time, we found 108 regulons with a median of 322 genes.

#### TF recovery

To assess whether the methods recover relevant TFs for the cell lines, we ranked the TFs included in Unibind for these cell lines (https://unibind.uio.no/) based on the number of target regions in the database, under the assumption that relevant TFs will be more bound to chromatin compared to residual or inactive TFs. We then built a cumulative recovery curve for each method, in which a unit is added if the TF in that ranking position is recovered by the method; and computed the AUC values using the first 40 positions of the ranking. Tau values for each TF were calculated using tispec (v0.99.0) and overlap plots were generated using UpsetR (v1.4.0).

#### TF-region

To assess the quality of the TF-region relationships inferred by the methods, we used as silver standard the predicted TF target regions from Unibind inferred from 309 ChIP-seq data sets (https://unibind.uio.no/) (Table S2) in these cell lines and F1 score as quality metric (i.e., the harmonic mean of precision and recall). FigR and SCENIC are excluded from the comparison since they do not report TF-region relationships.

#### Region-gene

In this comparison, Pando and SCENIC are excluded. Pando reports a score per TF-region-gene triplet, and because generally several TFs can bind to the same region, this results in several scores for the same region-gene pair. SCENIC does not calculate region-gene relationships. Among the methods, we find that SCENIC+ recovers the most links (402,838), followed by GRaNIE (311,168), FigR (66,024) and CellOracle/Cicero (13,123). For GRaNIE, FigR and CellOracle, these numbers are drastically reduced if we only include links that remain in the final GRN, resulting in 20,982, 26,586 and 2,867 region-gene links, respectively, while 271,718 are kept for SCENIC+. To assess the quality of the region-gene relationships inferred by the methods, we used with Hi-C data on 5 of the cell lines (namely IMR90 (ENCFF685BLG), GM12878 (ENCFF053VBX), HCT116 (ENCFF750AOC), HepG2 (ENCFF020DPP) and K562(ENCFF080DPJ)) available at https://www.encodeproject.org. Briefly, for each data set, we extracted SCALE normalized scores across bins of 5kb using Juicer Tools (v2.13.05), keeping only links with score > 10 and involving a bin that overlaps at least one of the consensus peaks and a TSS (+/-1000bp), resulting in 4,076,222 region-gene links on average. Finally, for each cell line, we computed the correlations between the scores given by the methods and the Hi-C scores for the top 100 marker genes in that cell line.

#### TF-gene

To assess the quality of the predicted target genes for each TF, two different approaches were used. First, we tested the predictability capacity of the methods, in other words, how well the regulons predicted by the methods could predict the transcriptome. Briefly, for each method and for each gene we trained a GBM regression model (sklearn v0.24.2) using as features the expression of the TFs that are predicted to regulate the gene, using 80% of the data. Then, we used these models to predict gene expression in the remaining 20% of the cells. As a quality metric we used the correlation between the observed and predicted values. Second, we assessed whether gene expression changes upon TF knockdown coincide with predicted target genes of the different methods, using 157 TF perturbation data sets from ENCODE (https://www.encodeproject.org/) (Table S3). Log Fold Changes between control and perturbed samples were calculated with DESeq2 (v1.28.1). We estimated the effect of these perturbation on all regulons of the different methods by performing a gene set enrichment analysis (GSEA) where genes are ranked based on the Log Fold Change compared to control data and predicted target genes for the TF are used as gene sets.

### Melanoma cell line analysis

#### scATAC-seq (10X genomics)

Two rounds of scATAC-seq (10X genomics) were performed on a mix of MM050, MM099, MM116, MM001, MM011, MM057 and MM087 (run 1) and a mix of MM031, MM074, MM047 and MM029 (run 2).

#### Cell culture

Patient-derived melanoma cell lines used in this study were obtained from the laboratory of Pr. Ghanem-Elias Ghanem (Istitut Jules Bordet, ULB, Belgium). The identity of each line has been determined using RNA-seq and ATAC-seq. Cell cultures used for experiments providing data to this study were tested for myoplasm contamination and were found to be negative. Cells were cultured in Ham’s F10 nutrient mux (Thermo Fisher Scientific, Waltham, MA) supplemented with 10% fetal bovine serum (FBS; Thermo Fisher Scientific) and 100 µg ml-1 penicillin–streptomycin. Cell cultures were kept at 37°C and 5% CO2. Prior to nuclei isolation cells were washed with phosphate buffered saline (1x PBS; Thermo Fisher Scienific), detached using trypsin (Thermo Fisher Scientific) and centrifuged at 1,000 r.p.m. for 5 min to remove the medium.

#### Nuclei isolation

To isolate nuclei from the different melanoma cell lines, protocol CG000169 (10x Genomics) was followed. Briefly, cells were washed with PBS + 0.04% BSA and cell concentration was determined with the LUNA-FL Dual Fluorescence Cell Counter. For each cell line, 500k cells were resuspended in 100 µl nuclei lysis buffer (10 mM Tris-HCl pH 7.4; 10 mM NaCl; 3 mM MgCl2; 0.1% Tween-20; 0.1% NP40; 0.01% Digitonin and 1% BSA in nuclease-free water). After 5 min incubation on ice, 1 ml chilled wash buffer was added to the lysed cells (10 mM Tris-HCl pH 7.4; 10 mM NaCl; 3 mM MgCl2; 0.1% Tween-20 and 1% BSA in nuclease-free water). The lysed cell suspension was centrifuged at 500 rcf for 5 min at 4°C and the pellet was resuspended in 1x nuclei buffer.

#### Library preparation

Single-cell libraries were generated using the GemCode Single-Cell Instrument and Single Cell ATAC Library & Gel Bead Kit v1-v1.1 and ChIP Kit (10x Genomics). Briefly, single nuclei suspended in 1x nuclei buffer were incubated for 60 min at 37°C with a transposase that fragments the DNA in open regions of the chromatin and adds adapter sequences to the ends of the DNA fragments. After generation of nanoliter-scale gel-bead-in-emulsions (GEMs), GEMs were incubated in a C1000 Touch Thermal Cycler (Bio-Rad) under the following program: 72°C for 5 min; 98°C for 30 s; 12 cycles of 98°C for 10 s, 59°C for 30 s, 72°C for 1 min; and hold at 15°C. After incubation, single-cell droplets were broken, and the single-strand DNA was isolated and cleaned using Cleanup Mix containing Silane Dynabeads. Illumina P7 sequence and a sample index were added to the single-strand DNA during library construction via PCR: 98°C for 45s; 9-13 cycles of 98°C for 20 s, 67°C for 30 s, 72°C for 20 s; 72°C for 1 min; and hold at 15°C. The sequencing-ready library was cleaned up with SPRIselect beads.

#### Sequencing

Before sequencing, the fragment size of every library was analyzed using the Bioanalyzer high-sensitivity chip. All 10x scATAC libraries of run 1 were sequenced on NextSeq2000 instrument (Illumina) with the following sequencing parameters: 51 bp read 1 – 8 bp index 1 – 24 bp index 2 – 51 bp read 2 and those of run 2 on NovaSeq6000 instrument (Illumina) with the following sequencing parameters: 50 bp read 1 – 8 bp index 1 – 16 bp index 2 – 49 bp read 2. For both scATAC-seq runs reads were mapped to GRCh38 and fragments files generated using cellranger-atac count command (v2.0.0) using default parameters.

#### Cell line annotation of scATAC-seq and scRNA-seq cells

To determine the identity of the scRNA-seq cells demuxlet was performed as described in^49^. To determine the identity of scATAC-seq cells demuxlet was performed using sample specific mutations obtained from bulk ATAC-seq data on individual cell lines. Genotypes of the individual cell lines were called using the bcftools mpileup command (v.1.11; options, *--max-depth 8000* and *--skip-indels*) and bcftools call command (v1.11; *options, -- multiallelic-caller*, *-variants-only*, *--skip-variants indels* and *--output-type b*). Only variants which were SNPs and not homozygous across samples were kept. To run demuxlet the scATAC-seq bam files were filtered to only contains reads covering SNPs and having a cell barcode (based on CB tag), reads were piledup using the popscle dsc-pileup command using default parameters and demuxlet was run using the popscle demuxlet command (using the option *–field GT*).

#### Quality control of scRNA-seq

The scRNA-seq data of baseline MM-lines, as available on GEO using accession number GSE134432 was preprocessed using Scanpy (v1.8.2)^110^. Briefly, genes expressed in less than 3 cells were removed and cells which expressed less than 200 genes, more than 4000 genes or had more than 15% counts in mitochondrial genes were removed. Doublets were detected and removed using Scrublet (v0.2.3)^102^ with default parameters. This resulted in 3,557 high quality cells.

#### Quality control of scATAC-seq and topic modeling

The scATAC-seq data was preprocessed using pycisTopic (v1.0.1.dev21+g8aa75d8). Briefly, based on the cell-to-cell line annotation consensus peaks (total of 360,302 peaks) were called as described above. Cells with less than 3.25 and 3.8 log number of unique fragements per cell, FRIP below 0.5 and TSS enrichment below 4 and 5, respectively for scATAC-seq run 1 and run 2, and doublets based on Scrublet (v0.2.3)102 calls with a threshold of 0.25 were removed. This resulted in 5,509 high quality cells. Topic modeling was performed as described above using LDA with the collapsed Gibbs sampler. A model of 30 topics was selected based on the stabilization of *Arun_2010*, *Cao_Juan_2009*, *Minmo_2011* and *loglikelihood* quality metrics.

#### Motif enrichment analysis

Motif enrichment analysis was performed using pycisTarget (v1.0.1.dev17+gd2571bf) as described above. For this, a custom score and ranking database was generated using *create_cisTarget_databases* python package using the DNA sequences of consensus peaks and all annotated motifs as input. Motif enrichment was performed using both the cisTarget and DEM algorithm on cell line and cell state (melanocytic, mesenchymal or intermedate) based DARs (log2 fold change above 1.5) and top 3,000 regions per topic. The motif enrichment analysis was run both once including promoters and excluding them. Promoters were defined as regions within 500 bp up- or downstream of the TSS of each gene. TSS’s for each gene were downloaded from Biomart (http://www.ensembl.org), using the pybiomart package (v0.2.0).

#### SCENIC+ analysis

Pseudo multiome data was generated for cell lines present in both scRNA-seq and scATAC-seq data, as described above using 5 cells per meta cell for a total of 936 meta cells. The SCENIC+ workflow was run using default parameters and as described above. Briefly, a search space of a maximum between either the boundary of the closest gene or 150 kb and a minimum of 1 kb upstream of the TSS or downstream of the end of the gene was considered for calculating region-to-gene relationships using gradient boosting machine regression. TF-to-gene relationships were calculated using gradient boosting machine regression between all TFs and all genes. Genes were considered as TFs if they were included in the TF list available on http://humantfs.ccbr.utoronto.ca/^120^. Final eRegulons were constructed using the GSEA approach in which region-to-gene relationships were binarized based gradient boosting machine regression importance scores using the 85th, 90th and 95th quantile; the top 5, 10 and 15 regions per gene and using the BASC method116 for binarization. Only eRegulons with a minimum of 10 target genes were retained. For each eRegulon cellular enrichment scores (AUC) of target genes and regions were calculated using the AUCell algorithm^20^. eRegulons for which the correlation coefficient between semi pseudo bulked per cell type (100 meta cells per cell type and 5 cells per meta cell) TF expression and region AUC scores was above 0.65 or below –0.75 were considered as high quality and used for downstream analysis. This resulted in 51 regulons, with a median of 176 genes and 183 regions per regulon. Analyses can be explored in Scope at https://scope.aertslab.org/#/scenic-v2 and UCSC at http://genome-euro.ucsc.edu/s/Seppe%20De%20Winter/scenicplus_mix_melanoma.

#### ChIP-seq enrichment in eRegulon target regions

MITF, SOX10 and TFAP2A ChIP-seq fastq files were downloaded from GEO using the respective accession numbers: GSE61965 (MITF and SOX10) and GSE67555 (TFAP2A). Reads were mapped to Grch38 using Bowtie2 (v2.4.4) using default parameters. Genomic coverage was calculated and stored as BigWig files using the bamCoverage function from deepTools (v3.5.0; options *-binSize 10*, -*-effectiveGenomeSize 2913022398* and *--normalizeUsing RPGC*). ChIP-seq coverage was calculated on the target regions of MITF(+), SOX10(+) and TFAP2A(+) and the regions targeted by any combination of two eRegulons using the pyBigWig package (v0.3.18) using 50 bins. Coverage was min-max normalized by using the minimum and maximum across all regions per ChIP-seq dataset.

#### Perturbation simulation

To simulate TF perturbations first a random forest regression model is trained to predict gene expression from TF expression, using the GradientBoostingRegressor fit function from the sklearn python package (v0.24.2) for each gene using the TFs predicted to regulate that gene as predictors for the gene. TFs which are predicted to regulate their own expression are excluded from the list of predictors before training the model. To simulate the effect of a TF knockout a simulated gene expression matrix is generated by predicting the expression of each gene using the expression of the predictor TFs while setting the expression of the TF of interest to zero. This simulation is repeated over several iterations, always using the newly simulated TF expression values as new predictor values for each gene. To visualize the effect of the perturbation in an embedding first the shift of the cells in the original embedding (delta embedding), calculated using eRegulon gene based AUC values, is estimated as described in^39,67^ based on eRegulon AUC values calculated using the simulated gene expression matrix. The delta embedding is used to draw arrows on the original embedding, using the streamplot or quiverplot function from the matplotlib (v3.5.2) python package. To prioritize TFs for their effect of driving melanocytic-to-mesenchymal transitions or vice versa average shifts along the first principal component, based on the delta embedding after 5 iterations of simulation, were calculated.

#### SOX10 KD RNA-seq analysis

Fastq files of bulk RNA-seq of MM001, MM011, MM031, MM057, MM074 and MM087 after SOX10 knockdown and non-targeting controls^49^ was downloaded from GEO using accession number GSE134432. After trimming sequencing adaptors from reads using fastq-mcf (v1.05; default parameters) reads were mapped to GRCh38 using STAR (v2.7.9a; options *--alignIntronMax 1*, *--alignIntronMin 2*), only reads with a mapping quality of minimal 4 were kept and a count matrix was generated using htseq-count (v0.9.1; options *-a 0*, *-m union* and *-t exon*). Log2 fold changes of SOX10 knockdown over non targeting control were calculated using the rlogTransformation command from DESeq2 (v1.34.0).

### Comparative analysis in the mammalian brain using SCENIC+

#### Human cortex data

Human motor cortex data was downloaded from https://data.nemoarchive.org/publication_release/Lein_2020_M1_study_analysis/Multimodal/sncell/SNARE/human/processed/counts/counts/M1/ (scRNA-seq count matrix and cell metadata) and https://data.nemoarchive.org/biccn/grant/u01_zhangk/zhang/multimodal/sncell/SNARE_ATACseq/human/processed/align/BICCN-H_20190523_SNARE2-AC_snapTools_190808/ (scATAC-seq fragment files).

#### Mouse cortex data

##### Mouse cortex dissection

All animal experiments were conducted according to the KU Leuven ethical guidelines and approved by the KU Leuven Ethical Committee for Animal Experimentation (approved protocol numbers ECD P007/2021). Mice were maintained in a specific pathogen-free facility under standard housing conditions with continuous access to food and water. Mice used in the study were 57 days old and were maintained on 14 hr light, 10 hr dark light cycle from 7 to 21 hr. In this study, cortical brain tissue from male P57 BL/6Jax was used. Animals were anesthetized with isoflurane, and decapitated. Cortices were collected and immediately snap frozen in liquid nitrogen.

#### Sample preparation

Five multiome experiments were performed, with small variations in sample preparation. For the sample labelled as “10x_complex” we used a modified protocol from the Nuclei Isolation from Complex Tissues for Single Cell Multiome ATAC + Gene Expression Sequencing Protocol (CG000375) from 10x Genomics. Briefly, a ∼1 cm3 frozen piece of mouse cortex tissue was transferred to 0.5 mL of ice-cold homogenization buffer (Salt-tris solution - 10 mM NaCl, 10 mM Tris 7.4, 3 mM MgCl2, 0,1% IGEPAL CA-63, 1 mM DTT, 1 U/µl of Protector RNase inhibitor (Sigma)) in a Dounce homogenizer mortar and thawed for 2 min. The tissue was homogenized with 10 strokes of pestle A and 10 strokes of pestle B until a homogeneous nuclei suspension was achieved. The resulting homogenate was filtered through a 70-µm cell strainer (Corning). The homogenizer and the filter are rinsed with an additional 0.5 µL of homogenization buffer. The tissue material was pelleted at 500 x rcf and the supernatant was discarded. The tissue pellet was resuspended in wash buffer (1% BSA in PBS + 1 U/µl of Protector RNase inhibitor (Sigma)). Nuclei were stained with 7AAD and viability sorted on a BD FACS Fusion into 5 mL low bind Eppendorf tube containing BSA with RNase inhibitor. The sorted nuclei were pelleted at 500 x rcf and the supernatant was discarded. Next, the nuclei were permeabilized by resuspending the pellet in 0.1x lysis buffer (Salt-tris solution - 10 mM NaCl, 10 mM Tris 7.4, 3 mM MgCl2, 0,1% IGEPAL CA-63, 0.01% Digitonin, 1% BSA, 1 mM DTT, 1 U/µl of Protector RNase inhibitor (Sigma)) and incubated on ice for 2 min. 1 mL Wash Buffer (Salt-tris solution - 10 mM NaCl, 10 mM Tris 7.4, 3 mM MgCl2, 0,1% Tween-20, 1% BSA, 1 mM DTT, 1 U/µl of Protector RNase inhibitor (Sigma)) was added and the nuclei were pelleted at 500 x rcf and the supernatant was discarded. The nuclei pellet was resuspended in diluted nuclei buffer (1x Nuclei buffer Multiome kit (10x Genomics)), 1 mM DTT, 1 U/µl of Protector RNase inhibitor (Sigma)).

For the sample labelled as “10x_no_perm” we used the above-described modified protocol from the Nuclei Isolation from Complex Tissues for Single Cell Multiome ATAC + Gene Expression Sequencing Protocol (CG000375) from 10x Genomics, omitting the nuclei permeabilization step. After sorting the nuclei were pelleted at 500 x rcf and the supernatant was discarded. The nuclei pellet was resuspended in diluted nuclei buffer (1x Nuclei buffer Multiome kit (10x Genomics)), 1 mM DTT, 1 U/µl of Protector RNase inhibitor (Sigma)).

For the sample labelled as “TST” nuclei isolation we used a modified protocol from the De Rop et al, 2022^122^. Briefly, a ∼1 cm3 frozen piece of mouse cortex tissue was transferred to 0.5 mL of ice-cold homogenization buffer (Salt-tris solution - 146 mM NaCl, 10 mM Tris 7.5, 1 mM CaCl2, 21 mM MgCl2, 250 mM sucrose, 0.03% Tween-20, 1% BSA, 25 mM KCl, 1x complete protease inhibitor, 1 mM DTT, 1 U/µl of Protector RNase inhibitor (Sigma)) in a Dounce homogenizer mortar and thawed for 2 min. The tissue was homogenized with 10 strokes of pestle A and 10 strokes of pestle B until a homogeneous nuclei suspension was achieved. The resulting homogenate was filtered through a 70-µm cell strainer (Corning). The homogenizer and the filter are rinsed with an additional 0.5 mL of homogenization buffer. The tissue material was pelleted at 500 x rcf and the supernatant was discarded. The tissue pellet was resuspended in a homogenization buffer without Tween-20. An addition 1.65 mL of homogenization buffer was topped up and mixed with 2.65 mL of gradient medium (75 mM sucrose, 1 mM CaCl2, 50% Optiprep, 5 mM MgCl2, 10 mM Tris 7.5, 0.5x complete protease inhibitor, 1 mM DTT, 1 U/µl of Protector RNase inhibitor (Sigma)). 4 mL of 29% iodoxanol cushion was prepared with a diluent medium (250 mM sucrose, 150 mM KCl, 30 mM MgCl2, 60 mM Tris 8) and was loaded into an ultracentrifuge tube. 5.3 mL of sample in homogenization buffer + gradient medium was gently layered on top of the 29% iodoxanol cushion. The sample was centrifuged at 7,700 x rcf, 4°C for 30 min and the supernatant was gently removed without disturbing the nuclei pellet. The nuclei pellet was resuspended in diluted nuclei buffer (1x Nuclei buffer Multiome kit (10x Genomics)), 1 mM DTT, 1 U/µl of Protector RNase inhibitor (Sigma)).

For the sample labelled as “10x_complex_UC” we designed a new protocol where we combined the “10x_complex” with “TST”. Briefly, a ∼1 cm3 frozen piece of mouse cortex tissue was transferred to 0.35 mL of ice-cold homogenization buffer (Salt-tris solution - 10 mM NaCl, 10 mM Tris 7.5, 3 mM MgCl2, 0,1% IGEPAL CA-63, 1 mM DTT, 1 U/µl of Protector RNase inhibitor (Sigma)) in a Dounce homogenizer mortar and thawed for 2 min. The tissue was homogenized with 10 strokes of pestle A and 10 strokes of pestle B until a homogeneous nuclei suspension was achieved. The resulting homogenate was filtered through a 70-µm cell strainer (Corning). The homogenizer and the filter are rinsed with an additional 0.65 mL of homogenization buffer. The homogenate was incubated on ice for 5 min, pelleted at 500 x rcf and the supernatant was discarded. 1 mL permeabilization buffer (PBS 1x, BSA 1%, 0,1% IGEPAL CA-63, 0.01% Digitonin, 1 U/µl of Protector RNase inhibitor (Sigma)) was added and incubated on ice for 2 min. Next, the pellet was resuspended, incubated on ice for extra 5 min and pelleted at 500 x rcf. The pelleted nuclei were resuspended in 1 mL wash buffer (PBS 1x, BSA 1%, 0.5 U/µl of Protector RNase inhibitor (Sigma)). An additional 1.65 mL of wash buffer was topped up and mixed with 2.65 mL of gradient medium (75 mM sucrose, 1 mM CaCl2, 50% Optiprep, 5 mM MgCl2, 10 mM Tris 7.5, 1 mM DTT). 4 mL of 29% iodoxanol cushion was prepared with a diluent medium (250 mM sucrose, 150 mM KCl, 30 mM MgCl2, 60 mM Tris 8) and was loaded into an ultracentrifuge tube. 5.3 mL of sample in wash buffer + gradient medium was gently layered on top of the 29% iodoxanol cushion. The sample was centrifuged at 7,700 x rcf, 4°C for 30 min and the supernatant was gently removed without disturbing the nuclei pellet. The nuclei pellet was resuspended in diluted nuclei buffer (1x Nuclei buffer Multiome kit (10x Genomics)), 1 mM DTT, 1 U/µl of Protector RNase inhibitor (Sigma)). Nuclei concentration was assessed by the LUNA-FL Dual Fluorescence Cell Counter.

For the sample labelled as “TST_NP40_004” we designed a new protocol starting from “TST”. Briefly, a ∼1 cm3 frozen piece of mouse cortex tissue was transferred to 0.5 mL of ice-cold homogenization buffer (Salt-tris solution - 146 mM NaCl, 10 mM Tris 7.5, 1 mM CaCl2, 21 mM MgCl2, 250 mM sucrose, 0.03% Tween-20, 0.01% BSA, 25 mM KCl, 1x complete protease inhibitor, 1 mM DTT, 1 U/µl of Protector RNase inhibitor (Sigma)) in a Dounce homogenizer mortar and thawed for 2 min. The tissue was homogenized with 10 strokes of pestle A and 10 strokes of pestle B until a homogeneous nuclei suspension was achieved. The resulting homogenate was filtered through a 70-µm cell strainer (Corning). The homogenizer and the filter are rinsed with an additional 500 mL of homogenization buffer. The homogenate was pelleted at 500 x rcf and the supernatant was discarded. 0.3 mL permeabilization buffer (Salt-tris solution - 10 mM NaCl, 10 mM Tris 7.5, 3 mM MgCl2, 1%, BSA, 25mM KCl, 250 mM Sucrose, 0,04 % IGEPAL CA-63, 0.01% Digitonin, 1 mM DTT, 1x complete protease inhibitor, 0.5 U/µl of Protector RNase inhibitor (Sigma)) was added and incubated on ice for 5 min. The homogenate was pelleted at 500 x rcf and supernatant discarded. The tissue pellet was resuspended in 1 mL Wash buffer (Salt-tris solution - 10 mM NaCl, 10 mM Tris 7.5, 3 mM MgCl2, 1%, BSA, 25mM KCl, 250 mM Sucrose,1 mM DTT, 1x complete protease inhibitor, 0.5 U/µl of Protector RNase inhibitor (Sigma)). An additional 1.65 mL of Wash buffer was topped up and mixed with 2.65 mL of gradient medium (75 mM sucrose, 1 mM CaCl2, 50% Optiprep, 5 mM MgCl2, 10 mM Tris 7.5, 1 mM DTT). 4 mL of 29% iodoxanol cushion was prepared with a diluent medium (250 mM sucrose, 150 mM KCl, 30 mM MgCl2, 60 mM Tris 8) and was loaded into an ultracentrifuge tube. 5.3 mL of sample in wash buffer + gradient medium was gently layered on top of the 29% iodoxanol cushion. The sample was centrifuged at 7,700 x rcf, 4°C for 30 min and the supernatant was gently removed without disturbing the nuclei pellet. The nuclei pellet was resuspended in diluted nuclei buffer (1x Nuclei buffer Multiome kit (10x Genomics)), 1 mM DTT, 1 U/µl of Protector RNase inhibitor (Sigma)). Nuclei concentration was assessed by the LUNA-FL Dual Fluorescence Cell Counter.

#### Library preparation

Single-nuclei libraries were generated using the 10x Chromium Single-Cell Instrument and NextGEM Single Cell Multiome ATAC + Gene Expression kit (10x Genomics) according to the manufacturer’s protocol. Briefly, the single mouse brain nuclei were incubated for 60 min at 37°C with a transposase that fragments the DNA in open regions of the chromatin and adds adapter sequences to the ends of the DNA fragments. After generation of nanolitre-scale gel bead-in-emulsions (GEMs), GEMs were incubated in a C1000 Touch Thermal Cycler (Bio-Rad) under the following program: 37°C for 45 min, 25°C for 30 min and hold at 4°C. Incubation of the GEMs produced 10x barcoded DNA from the transposed DNA (for ATAC) and 10x barcoded, full-length cDNA from poly-adenylated mRNA (for GEX). Next quenching reagent (Multiome 10x kit) was used to stop the reaction. After quenching, single-cell droplets were dissolved, and the transposed DNA and full-length cDNA were isolated using Cleanup Mix containing Silane Dynabeads. To fill gaps and generate sufficient mass for library construction, the transposed DNA and cDNA were amplified via PCR: 72°C for 5 min; 98°C for 3 min; 7 cycles of 98°C for 20 s, 63°C for 30 s, 72°C for 1 min; 72°C for 1 min; and hold at 4°C. The pre-amplified product was used as input for both ATAC library construction and cDNA amplification for gene expression library construction. Illumina P7 sequence and a sample index were added to the single-strand DNA during ATAC library construction via PCR: 98°C for 45 s; 7-9 cycles of 98°C for 20 s, 67°C for 30 s, 72°C for 20 s; 72°C for 1 min; and hold at 4°C. The sequencing-ready ATAC library was cleaned up with SPRIselect beads (Beckman Coulter). Barcoded, full-length pre-amplified cDNA was further amplified via PCR: 98°C for 3 min; 6-9 cycles of 98°C for 15 s, 63°C for 20 s, 72°C for 1 min; 72°C for 1 min; and hold at 4°C. Subsequently, the amplified cDNA was fragmented, end-repaired, A-tailed and index adaptor ligated, with SPRIselect cleanup in between steps. The final gene expression library was amplified by PCR: 98°C for 45 s; 5-16 cycles of 98°C for 20 s, 54°C for 30 s, 72°C for 20 s. 72°C for 1 min; and hold at 4°C. The sequencing-ready GEX library was cleaned up with SPRIselect beads.

#### Sequencing

Before sequencing, the fragment size of every library was analyzed using the Bioanalyzer high-sensitivity chip. All 10x Multiome ATAC libraries were sequenced on NovaSeq6000 instruments (Illumina) with the following sequencing parameters: 50 bp read 1 – 8 bp index 1 (i7) – 16 bp index 2 (i5) - 49 bp read 2. All 10x Multiome GEX libraries were sequenced on NovaSeq6000 instruments with the following sequencing parameters: 28 bp read 1 – 10 bp index 1 (i7) – 10 bp index 2 (i5) – 75 bp read 2. The generated fastq files were processed with cellranger-arc (v2.0.0) count function, with include *introns=True* option. Reads were aligned to Mus musculus reference genome (*ata-cellranger-arc-mm10-2020-A-2.0.0*).

#### Human cortex data analysis

High quality cells (84,159) selected by Bakken et al. were used for the analysis. scRNA-seq data was analyzed using Seurat (v4.0.3), using 47 PCs for dimensionality reduction and leiden clustering (with resolution 0.6). This resulted in 30 clusters (corresponding to 19 cell types) that were manually annotated based on marker gene expression. These cell type labels were used to create pseudobulks from which peaks were called with MACS2 (v2.1.2.1) and consensus peaks were derived using the iterative filtering approach (as previously described), resulting in 697,721 regions. Topic modelling was performed using Mallet, using 500 iterations and models with 10 topics and from 25 to 500 by an increase of 25, selecting the model with 50 topics based on the model selection metrics. pycisTarget (v1.0.1.dev17+gd2571bf) was run using a custom database with the consensus regions, on DARs and binarized topics (with Otsu thresholding), with and without promoters, and using pycisTarget and DEM. SCENIC+ was run with default parameters, using http://jul2018.archive.ensembl.org/ as Biomart host. High quality regulons were selected based on the correlation between gene-based regulon AUC and region-based regulon AUC (>0.4), and on the number of target genes (>30). This resulted in 142 regulons, with a mean of 315 genes and 768 regions per regulon.

#### Mouse cortex data analysis

scRNA-seq data was first analyzed using VSN (v0.27.0). Briefly, cells with at least 100 genes expressed and less than 1% of mitochondrial reads were kept. Doublets were removed using Scrublet (v0.2.3), with default parameters. 50 PCs were used as input for harmony, which was used to correct batch effects due to the sample preparation protocol, and the corrected PCs were used for dimensionality reduction and leiden clustering (resolution 1). This resulted in 41 clusters, that were annotated based on marker gene expression. Cell types not belonging to the cortex (such as medium spiny neurons from the striatum) were removed, and the data set was reanalyzed using Seurat (v4.0.3), using 52 PCs as input for harmony, as previously described, which were used for dimensionality reduction. This resulted in a data set with 21,969 high quality cells (based on scRNA-seq). These cell type labels were used to create pseudobulks from which peaks were called with MACS2 (v2.1.2.1) and consensus peaks were derived using the iterative filtering approach (as previously described), resulting in 568,403 regions. We further filtered the data set based on the scATAC-seq quality as well, keeping cells with at least 1,000 fragments, FRiP > 0.4 and TSS > 4, resulting in 19,485 cells. Topic modelling was performed using Mallet, using 500 iterations and models with 2 topics and from 5 to 100 by an increase of 5, selecting the model with 60 topics based on the model selection metrics. PycisTarget (v1.0.1.dev17+gd2571bf) was run using a custom database with the consensus regions, on DARs and binarized topics (with Otsu thresholding), with and without promoters, and using pycisTarget and DEM. SCENIC+ was run with default parameters, using http://nov2020.archive.ensembl.org/ as Biomart host. High quality regulons were selected based on the correlation between gene-based regulon AUC and region-based regulon AUC (>0.4), and on the number of target genes (>30). This resulted in 125 regulons, with a mean of 295 genes and 694 regions per regulon.

#### Cross-species comparison

Mouse gene names were converted to their orthologous human gene names based on the orthology table at http://www.informatics.jax.org/downloads/reports/HOM_MouseHumanSequence.rpt. Gene based human and mouse-to-human regulons were intersected, calculating the percentage of agreement as the number of overlapping genes divided by the size of the smallest regulon. Human region based regulons were liftovered to mm10 using UCSC Liftover (https://genome.ucsc.edu/cgi-bin/hgLiftOver) with default parameters, and regions overlapping mouse cortex consensus peaks and linked to the same (orthologous) gene in the two species were kept. Region based human-to-mouse and mouse regulons were intersected, calculating the percentage of agreement as the number of overlapping regions divided by the size of the smallest regulon. Human bigwig files were liftovered to the mm10 genome using CrossMap (v0.6.0). Analyses can be explored in Scope at https://scope.aertslab.org/#/scenic-v2 and UCSC at https://genome-euro.ucsc.edu/s/cbravo/SCENIC%2B_Cortex.

### GRN velocity

#### GRN velocity calculation

Briefly, we first order the differentiating cells by pseudotime. For each TF, we fit a standardized generalized additive model (GAM) along the pseudotime axis for its expression and its target genes (or regions) AUC values, using pyGAM (v0.8.0). We then map each cell in a certain quantile of the GAM TF expression model with the one in the same quantile in the GAM regulon AUC curve (posterior in the pseudotime axis). If there is no posterior cell in that quantile, the cell is mapped to itself. We define the differentiation force of a cell as the distance from the TF expression curve to its matching cell in the regulon AUC curve. When having multiple differentiation paths, we apply the same strategy in each path, and then average the two (or more) arrows available for cells found in more than one path. In addition, since we standardize the data prior to fitting the GAM model, we introduce a penalization curve standardized on the whole data set. This will prevent that if a TF (or its target) are not present in the branch, false arrows will not be drawn if the TF is not expressed compared to the rest of the data set. Differentiation forces can be plotted as an arrow grid in any dimensionality reduction of the data and prioritized per group of cells to identify key drivers in differentiation transitions. To visualize a regulon differentiation force, the distance in the embedding between matching cells is calculated (delta embedding). The delta embedding is used to draw arrows between the cells, using the streamplot or quiverplot function from the matplotlib (v3.5.2) Python package.

#### GRN velocity along oligodendrocyte differentiation in the mouse cortex

Oligodendrocyte cells (Oligodendrocytes Precursor Cells and oligodendrocytes) were subsetted from our in-house mouse cortex data set, resulting in 4,435 cells. The eRegulon AUC matrix was processed using Scanpy (v1.8.2) and the embedding-based pseudotime was derived using the diffmap and dpt functions. Differentiation arrows were inferred for cells above the 70% quantile in TF expression and default parameters. Prioritization of differentiation forces was done using the Regulon Specificity Score (RSS) metric using arrow lengths in each cell for each regulon as input values.

#### GRN velocity along the fly retina differentiation Data

We used scRNA-seq and scATAC-seq from the third instar larvae eye-antennal disc from Bravo et al. 2020^22^. In addition, we performed an additional scATAC-seq run using only eye discs (by cutting the antenna out of the eye-antennal disc), using the same protocol as described in Bravo et al. 2020. The analysis can be explored in Scope at https://scope.aertslab.org/#/scenic-v2 and UCSC at http://genome.ucsc.edu/s/cbravo/SCENIC%2B_EAD.

#### SCENIC+ analysis

scATAC-seq annotated cells by Bravo et al. 2020 were used to create pseudobulks from which peaks were called with MACS2 (v2.1.2.1) and consensus peaks were derived using the iterative filtering approach (as previously described), resulting in 39,732 regions. In addition, we added additional Drosophila cisTarget regions that did not overlap with these peaks, resulting in a data set with 127,711 regions. cisTarget regions are defined by partitioning the entire non-coding Drosophila genome based on cross-species conservation, resulting in more than 136,000 bins with an average size of 790 bp; and we found that using this region set increases the resolution for rare cell types, in which peak calling is difficult due to low amounts of cells.

Combining high quality cells from all the runs (based on cellranger), we obtained a data set with 23,317 cells. Topic modelling was performed using Mallet, using 500 iterations and models with 2 topics and from 5 to 100 by an increase of 5, selecting the model with 80 topics based on the model selection metrics. pycisTarget (v1.0.1.dev17+gd2571bf) was run using a custom database with the consensus regions, on DARs and binarized topics (with Otsu thresholding), with and without promoters, and using pycisTarget and DEM. We next mapped our new scATAC-seq atlas and our previously published scRNA-seq data into a virtual template of the eye-antennal disc using ScoMAP (v0.1.0), as described in Bravo et al., 2020. This resulted in a data set with 5,058 multiome pseudocells, for which both scRNA-seq and scATAC-seq measurements are available. This data was used as input for SCENIC+. SCENIC+ was run with default parameters, using http://dec2017.archive.ensembl.org as Biomart host and a 50kb window for the inference of region-gene links (instead of 150kb). High quality regulons were selected based on the correlation between gene-based regulon AUC and region-based regulon AUC (>0.4), and on the number of target genes (>10). This resulted in 153 regulons, with a mean of 216 genes and 323 regions per regulon.

#### GRN velocity

Differentiating cells in the eye disc were subsetted, resulting in a data set with 3,104 cells. The eRegulon AUC matrix was processed using Scanpy (v1.8.2). Cell annotations were refined based on leiden clustering on the eRegulon AUC matrix, resulting in 9 clusters along the fly retina differentiation. To identify branching points, we used PAGA (included in Scanpy), using 0.24 as threshold for *paga_compare*. The UMAP representation was redone using *init_pos=’paga’*. The embedding-based pseudotime was derived using the diffmap and dpt functions. Differentiation arrows were inferred for cells above the 70% quantile in TF expression, 0.2 as branch penalization and default parameters along the two differentiation paths (from progenitors to photoreceptors and from progenitors to interommatidial cells). Prioritization of differentiation forces was done using the Regulon Specificity Score (RSS) metric using arrow lengths in each cell for each regulon as input values. Prioritization of differentiation forces was done using the Regulon Specificity Score (RSS) metric using arrow lengths in each cell for each regulon as input values.

### Spatial GRN mapping

#### Human cerebellum 10x Visium Data

10x Visium data from the human cerebellum was downloaded from https://www.10xgenomics.com/resources/datasets/human-cerebellum-whole-transcriptome-analysis-1-standard-1-2-0. 10x single cell multiome data from the human cerebellum was downloaded from https://www.10xgenomics.com/resources/datasets/frozen-human-healthy-brain-tissue-3-k-1-standard-1-0-0. The analysis can be explored in Scope at https://scope.aertslab.org/#/scenic-v2 and UCSC at https://genome-euro.ucsc.edu/s/cbravo/SCENIC%2B_cerebellum.

#### SCENIC+ analysis and regulon mapping

Cells with at least 500 scRNA-seq reads and less than 5% of mitochondrial reads were kept, and doublets were removed using Scrublet (v0.2.3) with default parameters. scRNA-seq data was analyzed using Seurat (v4.0.3), using 37 PCs for dimensionality reduction and leiden clustereing (with resolution 0.6). This resulted in 15 clusters (corresponding to 13 cell types) that were manually annotated based on marker gene expression. These cell type labels were used to create pseudobulks from which peaks were called with MACS2 (v2.1.2.1) and consensus peaks were derived using the iterative filtering approach (as previously described), resulting in 435,834 regions. We further filtered the data set based on the scATAC-seq quality as well, keeping cells with at least log(unique fragments) > 3.5, FRiP > 0.2 and TSS > 4, resulting in 1,736 cells. Topic modelling was performed using Mallet, using 500 iterations and models with 2 topics and from 5 to 50 by an increase of 5, selecting the model with 40 topics based on the model selection metrics. pycisTarget (v1.0.1.dev17+gd2571bf) was run using a custom database with the consensus regions, on DARs and binarized topics (with Otsu thresholding), with and without promoters, and using pycisTarget and DEM. SCENIC+ was run with default parameters, using http://www.ensembl.org as Biomart host. High quality regulons were selected based on the correlation between gene-based regulon AUC and region-based regulon AUC (>0.6), and on the number of target genes (>10). This resulted in 111 regulons, with a mean of 100 genes and 171 regions per regulon. 10x Visium data was processed using Seurat (v4.0.3). SCENIC+ regulons were scored in the spots using AUCell (with the spot-gene matrix as input), with default parameters.

#### Molecular Cartography in the mouse cortex

##### Gene panel selection

100 genes were selected based on their gene expression patterns (marker genes for a cell type or group of cell types) on our in-house mouse cortex data set and literature (Table S4). In addition, we performed dimensionality reduction only using these 100 genes to ensure that all cell types could be distinguished with this gene panel.

#### Tissue sections

Mouse brain samples were fixed with PAXgene Tissue FIX solution (Resolve Biosciences) for 24 hours at room temperature followed by two hours in PAXgene Tissue Stabilizer (Resolve Biosciences) at room temperature. Samples were cryoprotected in a 30% sucrose solution (w/v) overnight at 4°C and frozen in 2-methylbutane (Sigma-Aldrich 106056) on dry ice. Frozen samples were sectioned with a cryostat (Leica CM3050) and 10µm thick sections were placed within the capture areas of cold Resolve Biosciences slides. Samples were then sent to Resolve BioSciences on dry ice for analysis. Upon arrival, tissue sections were thawed and rehydrated with isopropanol, followed by one min washes in 95% Ethanol and 70% Ethanol at room temperature. The samples were used for Molecular CartographyTM (100-plex combinatorial single molecule fluorescence in-situ hybridization) according to the manufacturer’s instructions (protocol v1.3; available for registered users), starting with the aspiration of ethanol and the addition of buffer DST1 followed by tissue priming and hybridization. Briefly, tissues were primed for 30 minutes at 37°C followed by overnight hybridization of all probes specific for the target genes (see below for probe design details and target list). Samples were washed the next day to remove excess probes and fluorescently tagged in a two-step color development process. Regions of interest were imaged as described below and fluorescent signals removed during decolorization. Color development, imaging and decolorization were repeated for multiple cycles to build a unique combinatorial code for every target gene that was derived from raw images as described below.

#### Probe Design

The probes for the 100 selected genes were designed using Resolve’s proprietary design algorithm. Briefly, the probe-design was performed at the gene-level. For every targeted gene all full-length protein coding transcript sequences from the ENSEMBL database were used as design targets if the isoform had the GENCODE annotation tag ‘basic’^123^. To speed up the process, the calculation of computationally expensive parts, especially the off-target searches, the selection of probe sequences was not performed randomly, but limited to sequences with high success rates. To filter highly repetitive regions, the abundance of k-mers was obtained from the background transcriptome using Jellyfish^124^ Every target sequence was scanned once for all k-mers, and those regions with rare k-mers were preferred as seeds for full probe design. A probe candidate was generated by extending a seed sequence until a certain target stability was reached. A set of simple rules was applied to discard sequences that were found experimentally to cause problems. After these fast screens, every kept probe candidate was mapped to the background transcriptome using ThermonucleotideBLAST^125^ and probes with stable off-target hits were discarded. Specific probes were then scored based on the number of on-target matches (isoforms), which were weighted by their associated APPRIS level^126^, favoring principal isoforms over others. A bonus was added if the binding-site was inside the protein-coding region. From the pool of accepted probes, the final set was composed by greedily picking the highest scoring probes. Gene names and Catalogue numbers for the specific probes designed by Resolve BioSciences are included in Table S4.

#### Imaging

Samples were imaged on a Zeiss Celldiscoverer 7, using the 50x Plan Apochromat water immersion objective with an NA of 1.2 and the 0.5x magnification changer, resulting in a 25x final magnification. Standard CD7 LED excitation light source, filters, and dichroic mirrors were used together with customized emission filters optimized for detecting specific signals. Excitation time per image was 1000 ms for each channel (DAPI was 20 ms). A z-stack was taken at each region with a distance per z-slice according to the Nyquist-Shannon sampling theorem. The custom CD7 CMOS camera (Zeiss Axiocam Mono 712, 3.45 µm pixel size) was used. For each region, a z-stack per fluorescent color (two colors) was imaged per imaging round. A total of 8 imaging rounds were done for each position, resulting in 16 z-stacks per region. The completely automated imaging process per round (including water immersion generation and precise relocation of regions to image in all three dimensions) was realized by a custom python script using the scripting API of the Zeiss ZEN software (Open application development).

#### Spot Segmentation

The algorithms for spot segmentation were written in Java and are based on the ImageJ library functionalities. Only the iterative closest point algorithm is written in C++ based on the libpointmatcher library (https://github.com/ethz-asl/libpointmatcher).

#### Preprocessing

As a first step all images were corrected for background fluorescence. A target value for the allowed number of maxima was determined based upon the area of the slice in µm² multiplied by the factor 0.5. This factor was empirically optimized. The brightest maxima per plane were determined, based upon an empirically optimized threshold. The number and location of the respective maxima was stored. This procedure was done for every image slice independently. Maxima that did not have a neighboring maximum in an adjacent slice (called z-group) were excluded. The resulting maxima list was further filtered in an iterative loop by adjusting the allowed thresholds for (Babs-Bback) and (Bperi-Bback) to reach a feature target value (Babs: absolute brightness, Bback: local background, Bperi: background of periphery within 1 pixel). This feature target values were based upon the volume of the 3D-image. Only maxima still in a zgroup of at least 2 after filtering were passing the filter step. Each z-group was counted as one hit. The members of the z-groups with the highest absolute brightness were used as features and written to a file. They resemble a 3D-point cloud. Final signal segmentation and decoding: To align the raw data images from different imaging rounds, images had to be corrected. To do so the extracted feature point clouds were used to find the transformation matrices. For this purpose, an iterative closest point cloud algorithm was used to minimize the error between two point-clouds. The point clouds of each round were aligned to the point cloud of round one (reference point cloud). The corresponding point clouds were stored for downstream processes. Based upon the transformation matrices the corresponding images were processed by a rigid transformation using trilinear interpolation. The aligned images were used to create a profile for each pixel consisting of 16 values (16 images from two color channels in 8 imaging rounds). The pixel profiles were filtered for variance from zero normalized by total brightness of all pixels in the profile. Matched pixel profiles with the highest score were assigned as an ID to the pixel. Pixels with neighbors having the same ID were grouped. The pixel groups were filtered by group size, number of direct adjacent pixels in group, number of dimensions with size of two pixels. The local 3D-maxima of the groups were determined as potential final transcript locations. Maxima were filtered by number of maxima in the raw data images where a maximum was expected. Remaining maxima were further evaluated by the fit to the corresponding code. The remaining maxima were written to the results file and considered to resemble transcripts of the corresponding gene. The ratio of signals matching to codes used in the experiment and signals matching to codes not used in the experiment were used as estimation for specificity (false positives).

#### Downstream Analysis

Final image analysis was performed in ImageJ using the Polylux tool plugin from Resolve BioSciences to examine specific Molecular CartographyTM signals. Nuclei segmentation was performed using CellProfiler (v4.2.1) based on the DAPI signal, using 30 and 100 as minimum and maximum diameter of objects, an adaptative threshold strategy and Otsu as thresholding method. Nuclei were expanded by 50 pixels. We mapped cell type labels and whole transcriptome gene expression from our in-house mouse cortex atlas using Tangram (v1.0.2), after library size correction, log normalization and correction of the gene expression values by sample using Combat (Scanpy v1.8.2). SCENIC+ regulons were scored in the nuclei using AUCell (with the nuclei-gene matrix as input), with default parameters.

## Data availability

Data generated in this manuscript, namely scATAC-seq in melanoma cell lines, 10x multiome in the mouse cortex and scATAC-seq in the Drosophila eye disc are available in GEO (GSE210749). Data from ENCODE Deeply Profiled Cell Lines was downloaded from https://www.encodeproject.org/, including bulk RNA-seq and ATAC-seq for 8 cell lines, namely MCF7 (ENCFF136ANW and ENCFF772EFK, for RNA-seq and ATAC-seq, respectively), HepG2 (ENCFF660EXG and ENCFF239RGZ), PC3 (ENCFF874CFD and ENCFF516GDK), GM12878 (ENCFF626GVO and ENCFF415FEC), K562 (ENCFF833WFD and ENCFF512VEZ), Panc1 (ENCFF602HCV and ENCFF836WDC), IMR90 (ENCFF027FUC and ENCFF848XMR) and HCT116 (ENCFF766TYC and ENCFF724QHH); and Hi-C data on 5 of the cell lines (IMR90 (ENCFF685BLG), GM12878 (ENCFF053VBX), HCT116 (ENCFF750AOC), HepG2 (ENCFF020DPP) and K562(ENCFF080DPJ)). ChIP-seq data sets and bulk RNA-seq experiments upon perturbation in these cell lines are described in Table S2 and Table S3, respectively. 10x multiome data on PBMC was downloaded from the 10x website (https://www.10xgenomics.com/resources/datasets/pbmc-from-a-healthy-donor-granulocytes-removed-through-cell-sorting-10-k-1-standard-1-0-0). scRNA-seq data of baseline MM-lines and bulk RNA-seq data after SOX10 knockdown were downloaded from GEO (GSE134432). MITF, SOX10 and TFAP2A ChIP-seq data were downloaded from GEO (GSE61965 (MITF and SOX10) and GSE67555 (TFAP2A)). SNARE-seq2 data on the mouse cortex was downloaded from https://data.nemoarchive.org/publication_release/Lein_2020_M1_study_analysis/Multimodal/sncell/SNARE/human/processed/counts/counts/M1/ (scRNA-seq count matrix and cell metadata) and https://data.nemoarchive.org/biccn/grant/u01_zhangk/zhang/multimodal/sncell/SNARE_ATACseq/human/processed/align/BICCN-H_20190523_SNARE2-AC_snapTools_190808/ (scATAC-seq fragment files). scATAC-seq and scRNA-seq data from the Drosophila eye-antennal disc were downloaded from GEO (GSE115476). 10x Visium data from the human cerebellum was downloaded from https://www.10xgenomics.com/resources/datasets/human-cerebellum-whole-transcriptome-analysis-1-standard-1-2-0 and 10x single cell multiome data from the human cerebellum was downloaded from https://www.10xgenomics.com/resources/datasets/frozen-human-healthy-brain-tissue-3-k-1-standard-1-0-0. All analyses can be explored in SCope (https://scope.aertslab.org/#/scenic-v2) and UCSC in the following sessions: PBMC (https://genome-euro.ucsc.edu/s/Seppe%20De%20Winter/scenicplus_pbmc), ENCODE cell lines (https://genome.ucsc.edu/s/cbravo/SCENIC%2B_DPCL), melanoma (http://genome-euro.ucsc.edu/s/Seppe%20De%20Winter/scenicplus_mix_melanoma), mouse and human cortex (https://genome-euro.ucsc.edu/s/cbravo/SCENIC%2B_Cortex), eye-antennal disc (http://genome.ucsc.edu/s/cbravo/SCENIC%2B_EAD) and human cerebellum (https://genome-euro.ucsc.edu/s/cbravo/SCENIC%2B_cerebellum).

## Code availability

PycisTopic is available at https://github.com/aertslab/pycisTopic. Pycistarget is available at https://github.com/aertslab/pycistarget. SCENIC+ is available at https://github.com/aertslab/scenicplus. Detailed tutorials and documentation on the SCENIC+ workflow are available at https://scenicplus.readthedocs.io, while tutorials on pycisTopic and pycistarget (within the SCENIC+ workflow and as standalone packages) are available at https://pycisTopic.readthedocs.io and https://pycistarget.readthedocs.io, respectively. Code to generate custom cisTarget databases is available at https://github.com/aertslab/create_cisTarget_databases. Our implementation of Cluster-Buster is available at https://resources.aertslab.org/cistarget/programs/cbust. Notebooks to reproduce the analyses presented in this manuscript are available at https://github.com/aertslab/scenicplus_analyses.

## Acknowledgements

Computing was performed at the Vlaams Supercomputer Center (VSC). This work is funded by the following grants to S.Ae: ERC Consolidator Grant (724226_cis-CONTROL), ERC Proof of Concept (963884), Special Research Fund (BOF) KU Leuven (grant C14/18/092), Foundation Against Cancer (2020-062), and FWO (grants G0B5619N and G094121N); PhD fellowships from the FWO to C.B.G.-B. (11F1519N) and S.D.W (1191323N), and postdoctoral fellowships from FWO to N.H (1273822N) and Stichting tegen Kanker (Foundation against Cancer) to J.W (2019-100). The authors also thank members of various groups that make curated position weight matrices publicly available, including T. Hughes (cis-bp), M. Bulyk (Uniprobe), A. Mathelier (Jaspar), V. Makeev (Hocomoco) and many others, listed in Table S1. We thank Resolve Biosciences, specially Jeroen Aerts, for performing the Molecular Cartography experiments in the mouse cortex; and Janssen Pharmaceutica, VIB Tech Watch and the VIB single-cell accelerator for their help and funding for generating the mouse cortex data.

## Author contributions

C.B.G.-B., S.D.W and S.Ae conceived the study. C.B.G.-B. developed pycisTopic, C.B.G.-B. and S.D.W co-developed pycistarget and the SCENIC+ modules and workflow, and G.H. developed the code to generate custom cisTarget databases. C.B.G.-B and G.H. made the SCENIC+ motif collection. C.B.G.-B and S.D.W performed the computational analyses, with assistance of G.H., N.H and S.Ai. I.M and S.P generated the mouse cortex multiome data and J.W. performed the single-cell ATAC-seq experiments on the melanoma cell lines, with assistance of V.C. C.B.G.-B., S.D.W and S.Ae wrote the manuscript.

## Supplementary Tables

Table S1. Motif collections included in the SCENIC+ motif collection.

Table S2. ENCODE Chip-seq data sets used for motif enrichment and GRN inference benchmarks.

Table S3. ENCODE TF perturbation data sets used for the GRN inference benchmark.

Table S4. Molecular Cartography probes.

## Supplementary Note 1

### Benchmark of GRN inference methods

While all methods aim to infer gene regulatory networks, there are conceptual differences that result in different types of GRNs and insights (Fig 3). For instance, SCENIC^2^ only uses scRNA-seq, while the remaining methods use (single-cell) multi-omics data as input. CellOracle^3^, like SCENIC, only provides TF-Gene networks (despite using chromatin information internally), infers region-gene links only based on accessibility and compared to other methods does not assess repression, but can predict perturbation effects. Pando^4^ co-optimizes TF-region-gene relationships, resulting in one unique score for each combination, which hinders the assessment of region-gene relationships as most regions are targeted by more than one TF, resulting in several scores for each region-gene pair. FigR^5^ derives DORCs (Domains of Regulatory Chromatin), which consist of sets of regions (50kb from the gene TSS by default) whose accessibility correlates with the gene expression. Motif enrichment is then performed across the whole DORC, which prevents assessing to which region within the DORC a TF is binding. GRaNIE^6^ is the only tool conceptually similar to SCENIC+ (consisting of different steps to infer TF-region and region-gene relationships and a final eGRN compilation step), but it is designed for bulk data. As the original ENCODE Deeply Profiled Cell Lines data are bulk profiles, we tested GRaNIE with these bulk data. However, its performance was very poor, only finding 26 TFs and 11,106 TF-region-gene links. Interestingly, when we applied it to our simulated single-cell data set (used for all the other tools), its recovery increased (finding 39 TFs and 44,666 TF-region-gene links); hence, we report the latter results. Finally, SCENIC+ not only builds enhancer-GRNs (with TF-region and region-gene information), but can also assess repression, regulon specificity, the effect of TF perturbation (as CellOracle) and prioritize eGRNs driving differentiation processes. Importantly, both SCENIC and SCENIC+ use the SCENIC+ motif collection with thousands of motif/clusters, while most of the remaining methods rely on few hundreds of motifs. To assess the effect of clustering in the motif collection, we have also benchmarked SCENIC using the unclustered and the clustered collections (Fig S5, S7), and both at the same time (Fig 3); and SCENIC+ using consensus peaks (642,982 peaks, Fig 3) and SCREEN regions^7^, that overlap with ∼80% of the consensus peaks in this case (Fig S5, S7), finding that results between the variants for each method give similar results.

SCENIC and FigR were excluded from the TF-region benchmark since they perform motif enrichment in a space around the TSS or DORCs rather than individual regions, respectively. SCENIC and Pando were excluded from the region-gene benchmark since SCENIC does not calculate region-gene relationships and Pando reports a score per TF-region-gene triplet. Because generally several TFs can bind to the same region, this results in several scores for the same region-gene pair.

Of note, we found that knocking down different master regulators in K562 (e.g. STAT5A, HOXB9, HOXB4, SFPQ, GATA1, GATA2, ARID3A) can result in similar effects and affect similar regulons. These downstream effects can be largely explained by indirect effects of the TF-knockdown experiment direct interactions between the TFs (i.e the targeted TF activates that other TF) or cooperativity (i.e. these TFs target the same genes) (Fig S7). Interestingly, we also observed that some repressive regulons are upregulated upon knockdown for the corresponding TFs, such as HOXB4, HOXB9 and ARID3A8,9, showing that SCENIC+ is also able to accurately recover repressive interactions (Fig S7).

## Supplementary Note 2

### Predicting repressive interactions using SCENIC+

Transcriptional repression is an important biological mechanism mostly studied in the context of developmental biology^10–13^, in which transcription factors (TFs) of one cell type repress TFs of another (adjacent) cell type thereby locking in the fate of the first cell type. Repression is mostly analyzed using genetic gain and loss-of-function experiments^10^, in which it is difficult to disentangle direct from indirect effects. For this reason, the mechanisms by which TFs induce repression on a molecular level are poorly understood.

The eukaryotic genome is compacted in chromatin which maintains a restrictive ground state/default “off” state. To activate transcription, (combinations of) TFs bind enhancers^14^, displacing nucleosomes and opening up the chromatin thereby lifting the restrictive ground state. In the context of this model in order to induce repression the chromatin has to be closed again. For this, two main molecular mechanisms are described. First, the access of the activator TF(s) to the chromatin can be limited^10^. For this, the cell can simply stop transcribing the activator TF(s); the cell can transcribe a “repressor” TF with a similar DNA binding domain as the activator thereby excluding the activator by going into direct competition for binding the DNA; the activator TF(s) can be sequestered away from the nucleus using protein-protein interactions; or the amount of activator protein can be limited due to repression at the RNA-level. Second, the chromatin can be actively closed using repressor TFs which, upon binding the DNA, recruit repressive co-factors^15^.

To detect this type of repression (repression by chromatin closing) SCENIC+ relies on observing negative correlations between on the one hand TF expression and target gene expression and on the other hand region accessibility and target gene expression. A problem with this approach ensues whenever two or more TFs are expressed in the system of interest for which the motif is the same or very similar but the expression is anti-correlated. In this case it is impossible to deduce whether the chromatin is being closed simply by the absence of the activator TF or it is actively closed by the action of the inferred repressor TF (potentially recruiting co-repressors) (Fig. S12a). This problem is most prevalent in species with many paralogs for which there are many TFs belonging to the same family, for example human^16^.

We illustrate this issue in the melanoma cell line analysis. In this system, the expression of the following pairs of TFs is anti-correlated but their motifs are very similar: SOX10 and SOX9, MITF and TCF4 and TCF4 and MITF (Fig. S12b). Because of this, SCENIC+ predicts repressive eRegulons for SOX9, TCF4 and MXI1 (Fig. S12c) and the predicted target regions of these TFs strongly overlap with the predicted target regions of their activating partners (Fig. S12d). Whether SOX9, TCF4 and MXI1 actively close, in the mesenchymal state, the regions opened up by respectively SOX10, MITF and TCF4 in melanocytic/intermediate state can not be concluded by the SCENIC+ analysis on its own. Using SCENIC+ alone both the scenario in which the regions are passively closed simply by the absence of the activators in the mesenchymal state or actively closed by the presence of the repressors in the mesenchymal state has the same likelihood.

Even though one has to be cautious with the interpretation of repressors predicted by SCENIC+, these predictions can still lead to novel insights. An interesting example in the melanoma cell line analysis is HES1. SCENIC+ predicts a repressive eRegulon for HES1, in line with the function of its ortholog in Drosophila melanogaster^17^. HES1 is a basic helix-loop-helix transcription factor and thus its DNA binding motif is very similar to MITF however its expression is not exactly anti-correlated to MITF (Fig. S12e). MITF and HES1 are co-expressed in MM031, MM011 and all cell lines of the intermediate state (Fig. S12e) while MITF but not HES1 is expressed in MM001. Given that the expression of HES1 is a better prediction for the region accessibility of predicted MITF target regions compared to the expression of MITF itself (Fig. S12f) and the predicted target regions and genes of HES1 strongly overlap with those of MITF (Fig. S12g) we hypothesize that HES1 represses the action of MITF in melanoma cell lines which co-express both factors thus restricting MITF target gene expression to MM001 (Fig. S8d). This is in line with recent reports where Notch signaling is shown to counteract the effect of MITF in melanoma^18^ and where HES1 is shown to enhance epithelial-to-mesenchymal transitions^19^.

**Fig S1.**
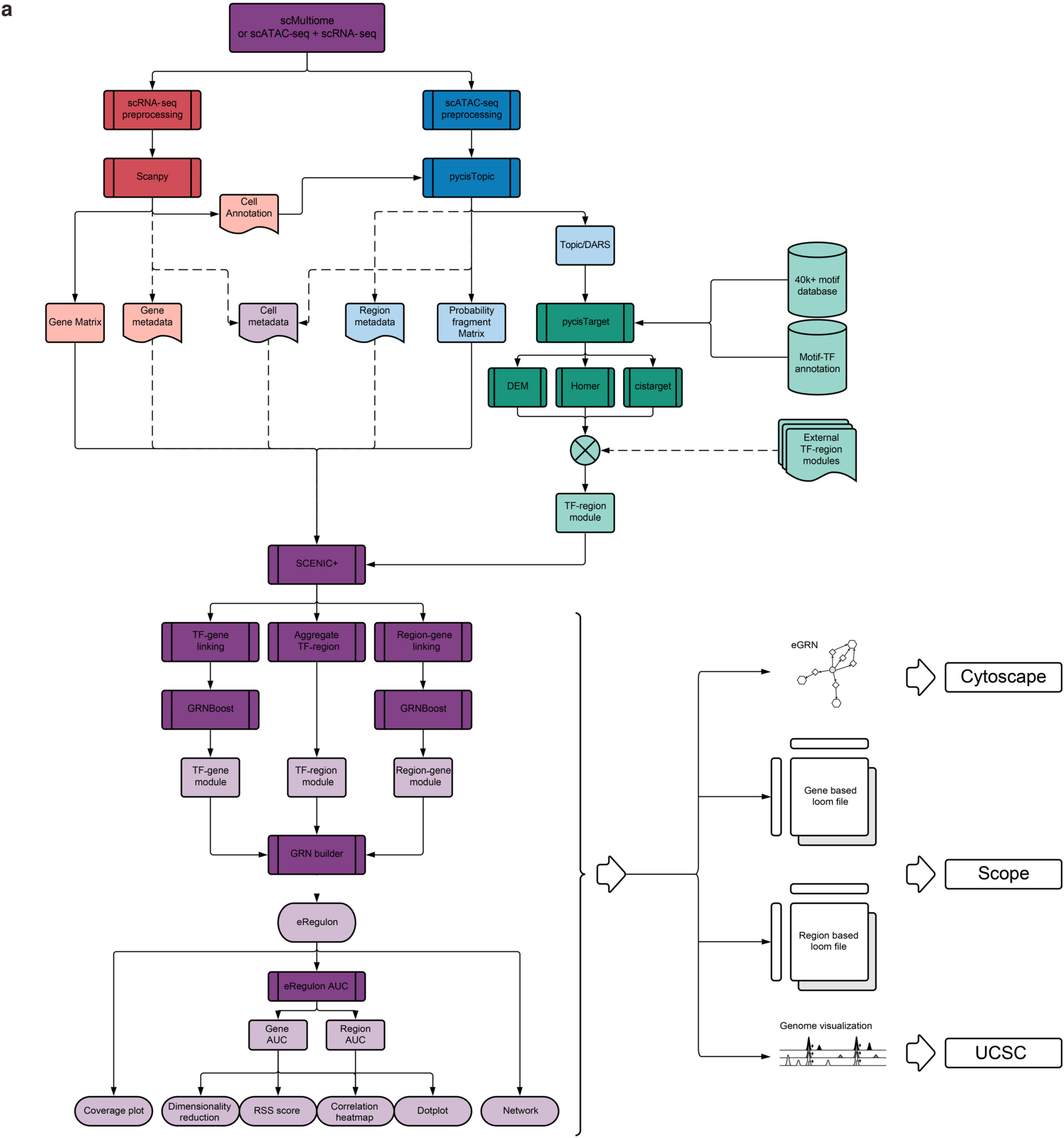
The SCENIC+ workflow. Diagram showcasing the main steps and outputs performed during the SCENIC+ workflow. Single cell RNA-seq processing steps (e.g. Scanpy) are shown in red; pycisTopic steps, in blue; pycisTarget, in green; and SCENIC+, in purple.

**Fig S2.**
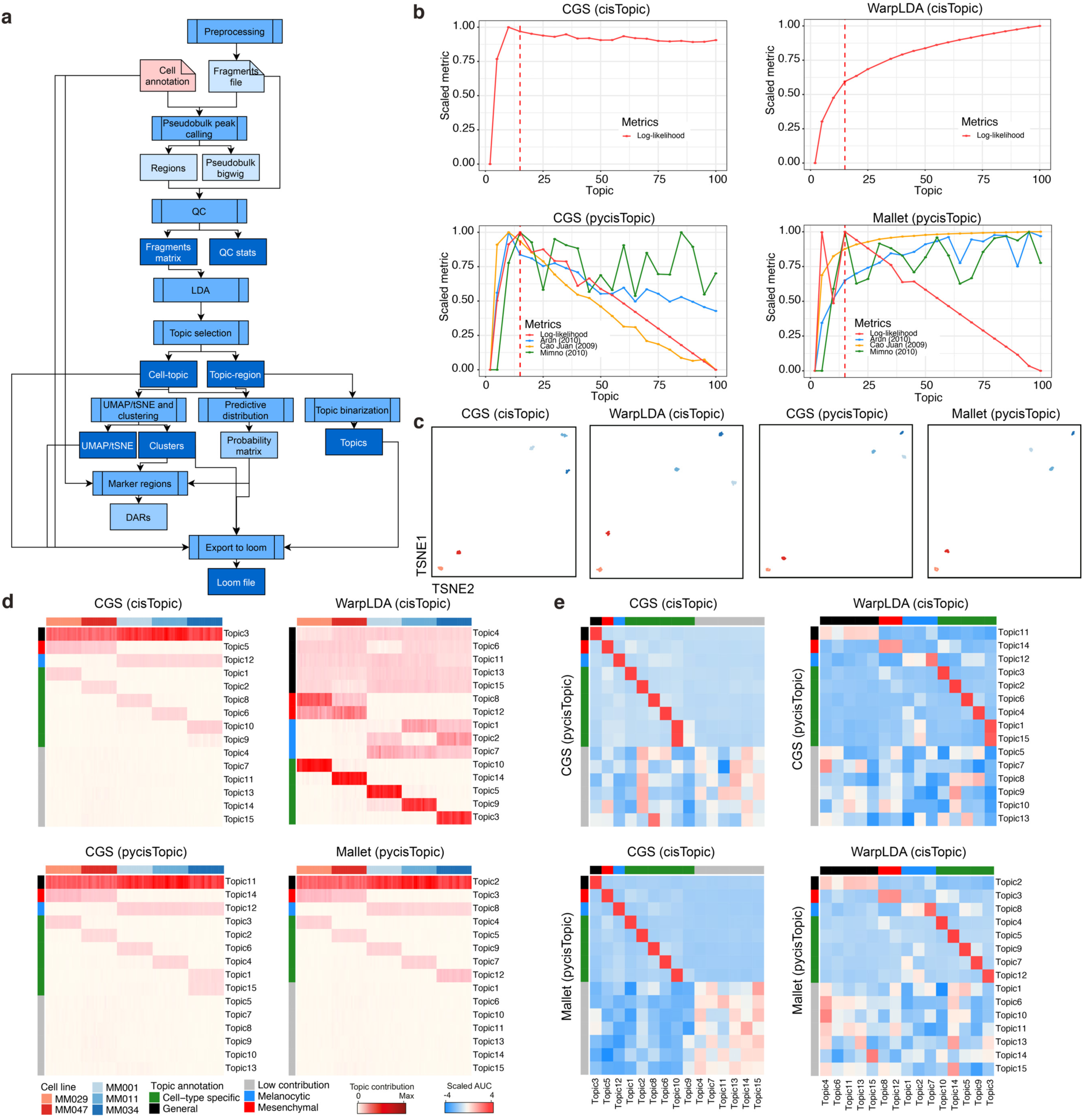
PycisTopic provides faster and equally accurate results compared to cisTopic in a simulated single-cell epigenomics data set from melanoma cell lines. **a.** pycisTopic workflow with key steps. **b.** Model selection for models with different parameter optimization methods, namely Collapsed Gibbs Sampler (CGS) and WarpLDA with cisTopic, and CGS and Mallet with pycisTopic. cisTopic relies on the log-likelihood per model; while pycisTopic incorporates additional measurements including coherence (Minmo (2010)), a density based metric (Cao Juan (2009) and a divergence based metrics (Arun (2010)). **c.** Cell-topic dimensionality reduction for each of the models (100 cells). Red clusters denote the 2 mesenchymal cell lines, blue clusters depict the 3 melanocytic cell lines. **d.** Cell-topic enrichment heatmap for each of the models. General topics are shown in black; mesenchymal, in red; melanocytic, in blue; cell line specific in green; and low contributing in grey. **e.** AUCell enrichment of topics between different models.

**Fig S3.**
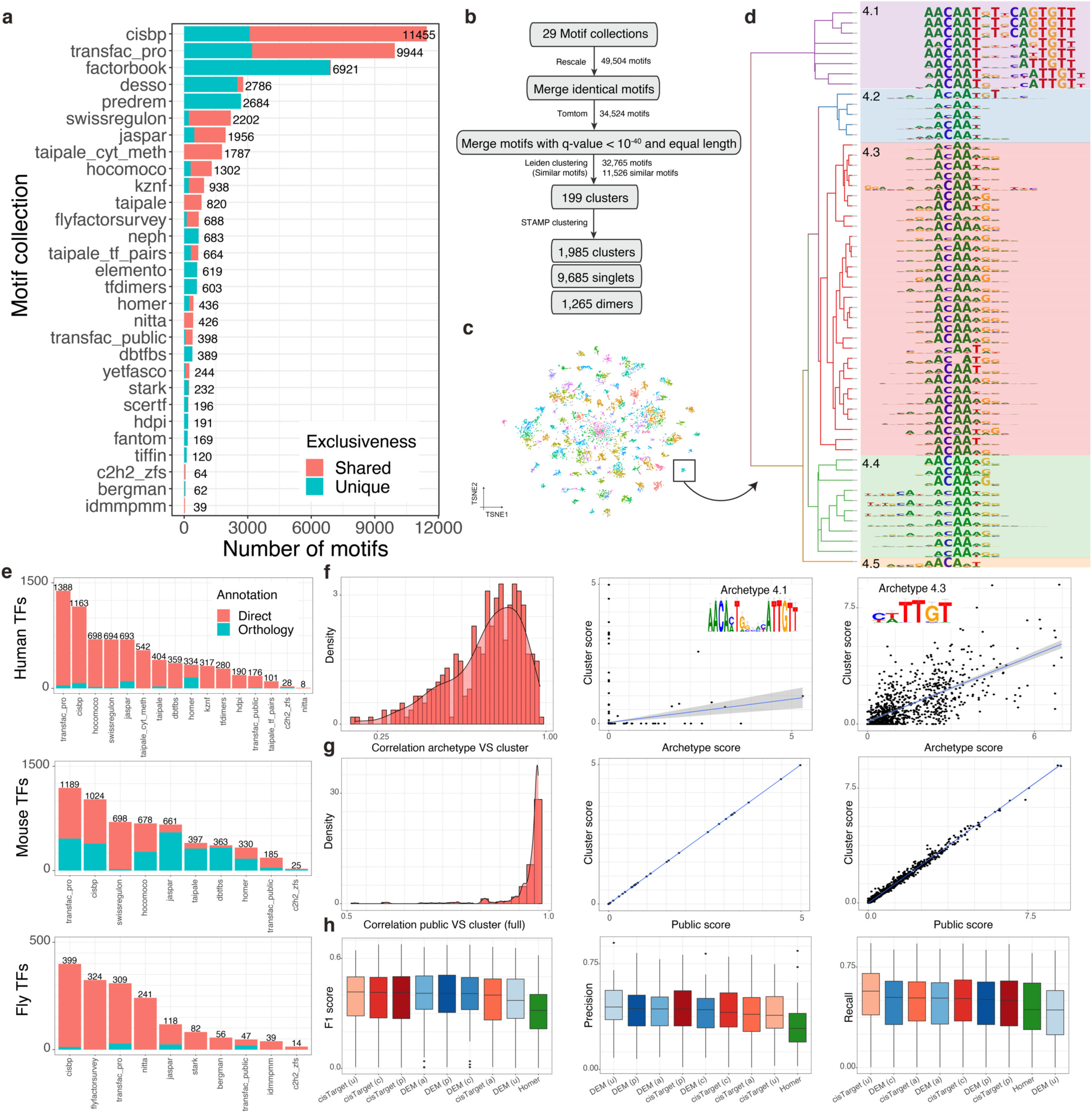
The SCENIC+ motif collection. **a.** Number of motifs per motif collection in the SCENIC+ motif collection. Some motifs are uniquely included in one collection, while others can be found in several**. b.** Workflow depicting the motif collection cluster strategy **c.** tSNE based on Tomtom motif similarity values (11,526 motifs) colored by Leiden clustering (res 25). **d.** STAMP clustering tree for cluster 4. Subclusters are indicated with different colors. **e.** Number of motifs annotated directly or by orthology in each motif collection. **f.** Left: Histogram showing the correlation between scores in chr19 regions using the archetype or all motifs in a cluster for scoring with Cluster-Buster. Middle: Correlation between scores for cluster 4.1 using the archetype or all motifs in chr19 regions. Right: Correlation between scores for cluster 4.3 using the archetype or all motifs in chr19 regions. **g.** Left: Histogram showing the correlation between scores in chr19 regions using all motifs in a cluster for scoring with Cluster-Buster or all motifs except for Transfac Pro PWMs. Middle: Correlation between scores for cluster 4.1 using all motifs in chr19 regions or all motifs except for Transfac Pro PWMs. Right: Correlation between scores for cluster 4.3 using the archetype or all motifs in the cluster in chr19 regions. **h.** F1 score (left), precision (middle) and recall (right) distributions upon comparison of TF cistromes derived from motif enrichment from 309 ChIP-seq data sets from ENCODE with different methods.

**Fig S4.**
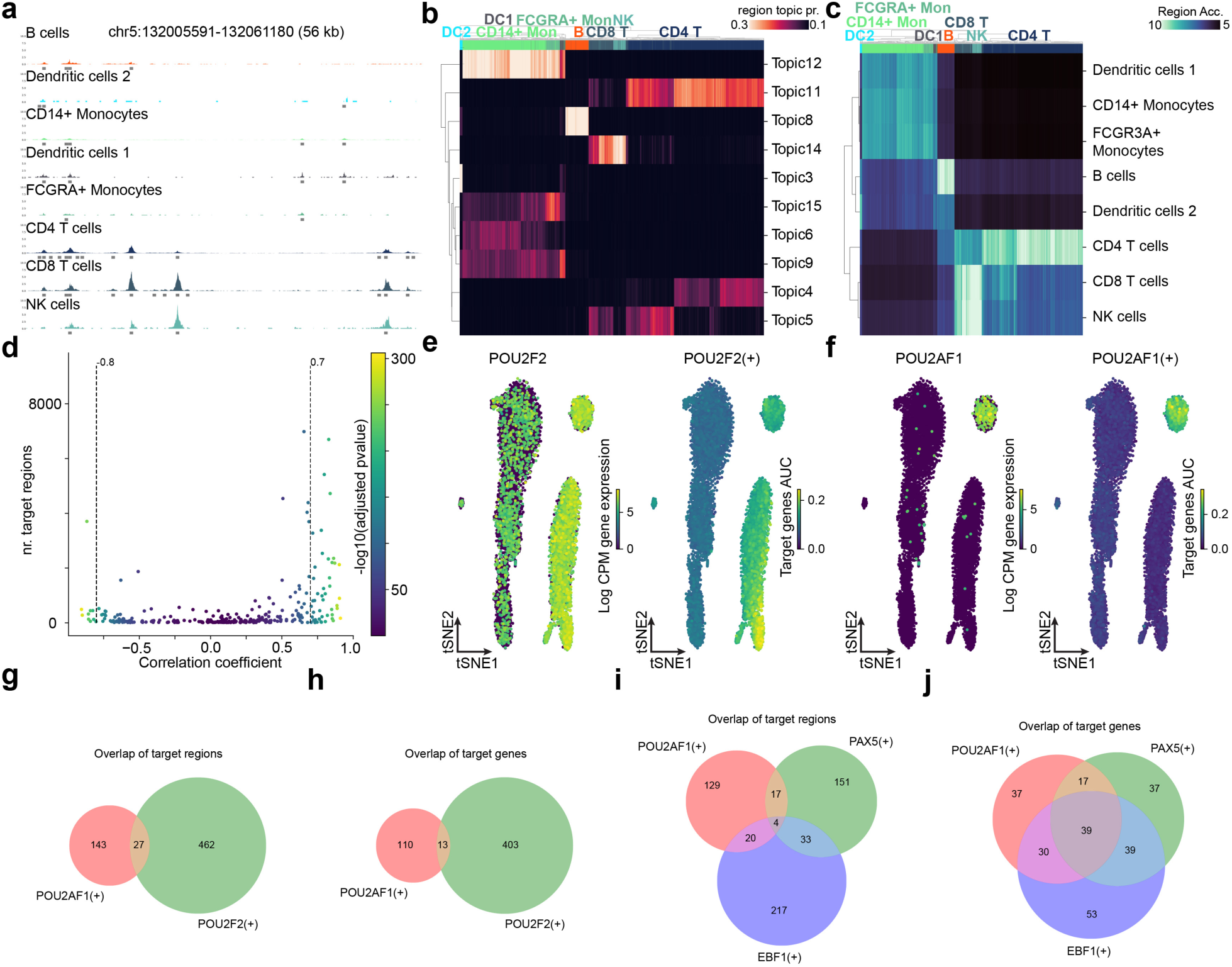
SCENIC+ analysis on peripheral blood mononuclear cells. **a.** Coverage track illustrating pseudobulk coverage generated by pycisTopic for different cell types. Gray lines represent consensus peaks across cell types. **b.** Heatmap of cell topic probabilities (cell topic prob.). **c.** Heatmap of imputed accessibility (Region Acc.) of differentially accessible regions across cell types. **d.** Scatter plot showing number of target regions vs TF expression-to-region AUC Pearson correlation coefficients for each eRegulon. Color scale represents -log10(adjusted p value) of the Pearson correlation coefficients. eRegulons are selected based on a threshold on the correlation coefficient, indicated by dotted line. **e.** Expression and predicted target gene AUC values of POU2F2. **f.** Expression and predicted target gene AUC values of POU2AF1. **g.** Venn diagram of POU2AF1 and POU2F2 target regions. **h.** Venn diagram of POU2AF1(+) and POU2F2(+) target **i.** Venn diagram of POU2AF1, EBF1 and PAX5 target regions. **j.** Venn diagram of POU2AF1, EBF1 and PAX5 target genes.

**Fig S5.**
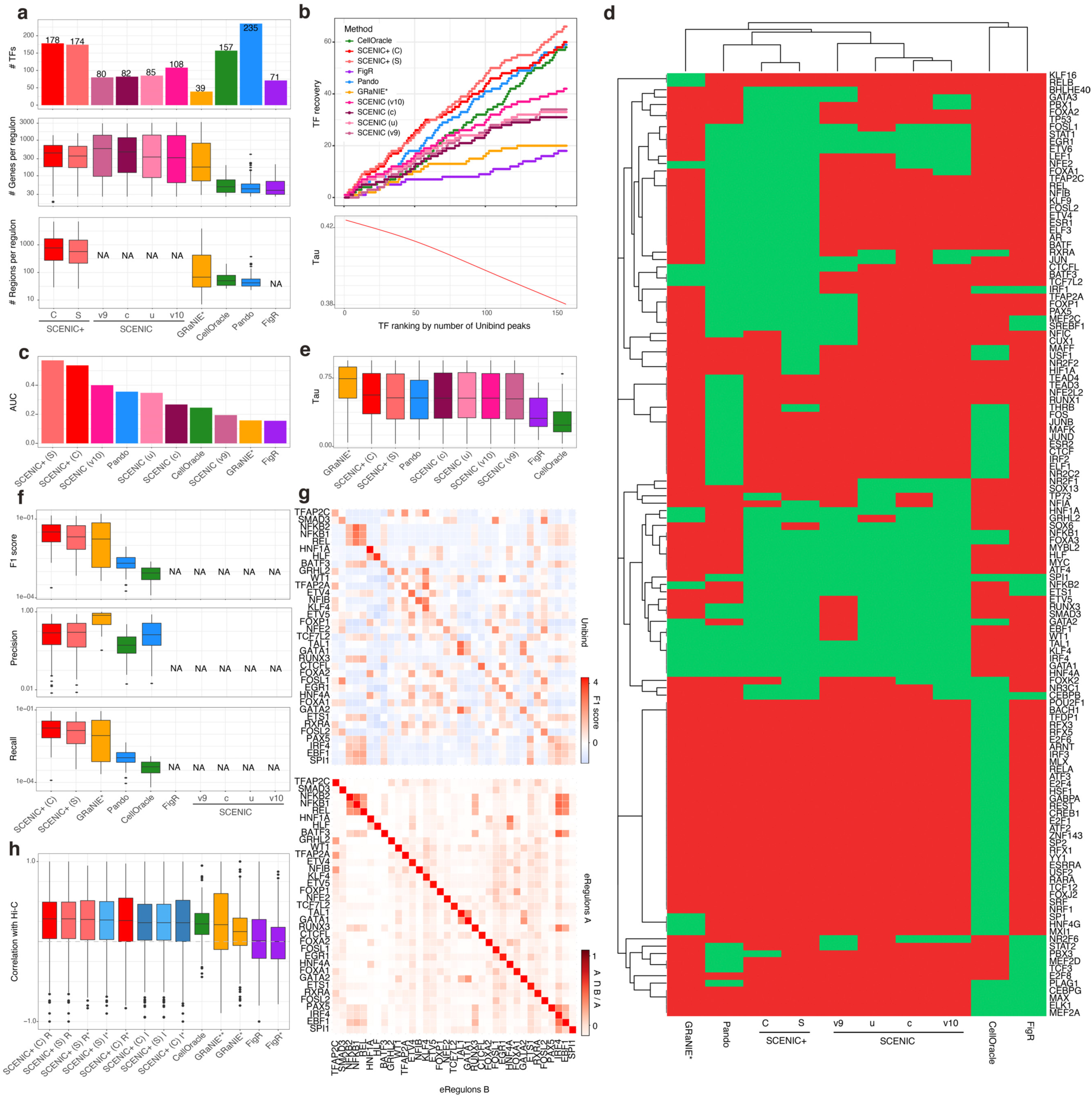
Benchmarking TF recovery, TF-to-region and TF-to gene inferred relationships with different approaches. **a.** Number of TFs recovered per method (top), and distributions showing the number of genes and regions found in each regulon per method (middle and bottom, respectively). **b.** Cumulative TF recovery for each method using as x-axis TFs ranked based on the number of Unibind peaks and GAM fitted Tau value across the ranking. **c.** AUC values per method using the top 40 TFs. **d.** Heatmap indicating whether a TF is found (green) or not (red) by each method. The TFs shown correspond to those found in Unibind and by at least one method. **e.** Tau values distributions for the TFs recovered by each method. **f.** F1 score, precision and recall distributions resulting from the comparison of the target regions found by the methods and Unibind. **g.** Top: Heatmap showing F1 score values for each SCENIC+ regulon and the target regions for that TF in Unibind. Bottom: Heatmap showing the overlap between the regions of the regulons indicated by the rows and columns, divided by the size of the regulons in the columns. **h.** Correlation between Hi-C links for the top 100 markers genes for each of the cell lines where Hi-C is available (namely IMR90, GM12878, HCT116, HepG2 and K562) and the region-gene scores from different methods. * indicates if only region-gene pairs found in the final eGRN are used for the calculations. SCENIC+ (C) or (S) indicates that SCENIC+ was run using consensus peaks or SCREEN regions for the pycisTarget database; SCENIC v9, v10c, v10u, and v10 indicate whether SCENIC was run using the previous version of the motif collection, the clustered SCENIC+ motif collection, the unclustered SCENIC+ motif collection or both the clustered and unclustered motif collection, respectively. NA is used in the panels when that information is not available when using that method.

**Fig S6.**
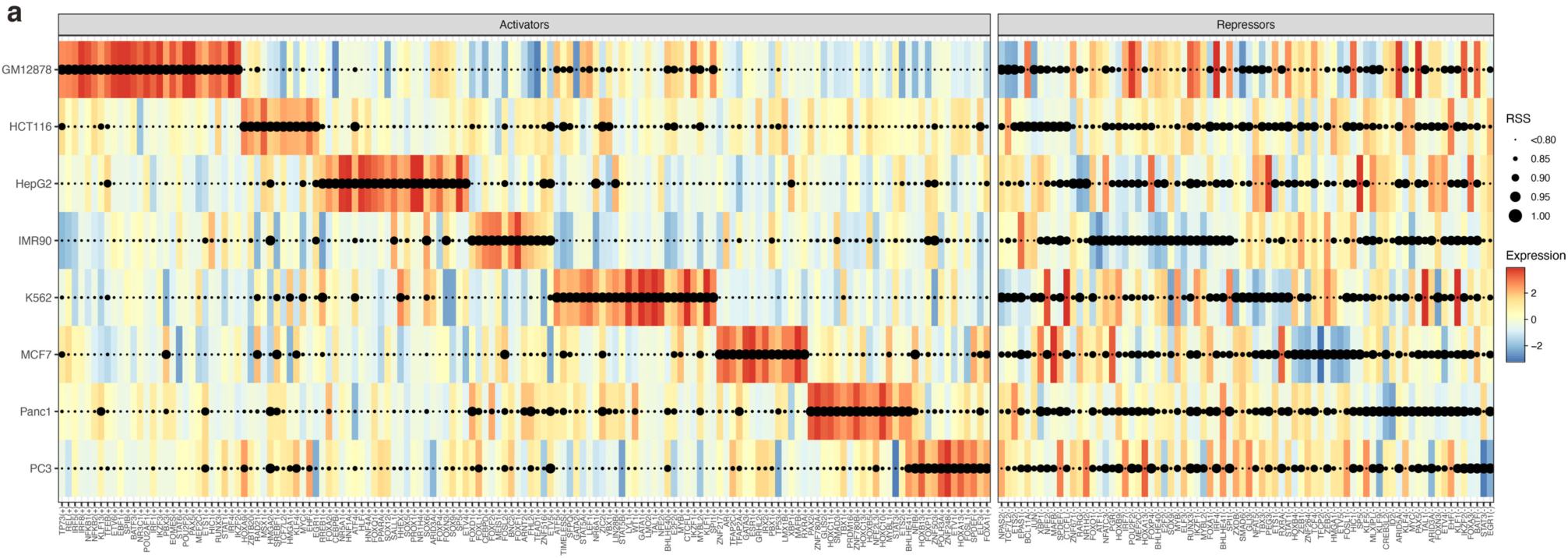
Inferred regulons by SCENIC+ in the ENCODE Deeply Profiled Cell Lines. **a.** Dotplot showing TF expression of the regulon on a color scale and cell type specificity of the regulon (RSS) on a size scale.

**Fig S7.**
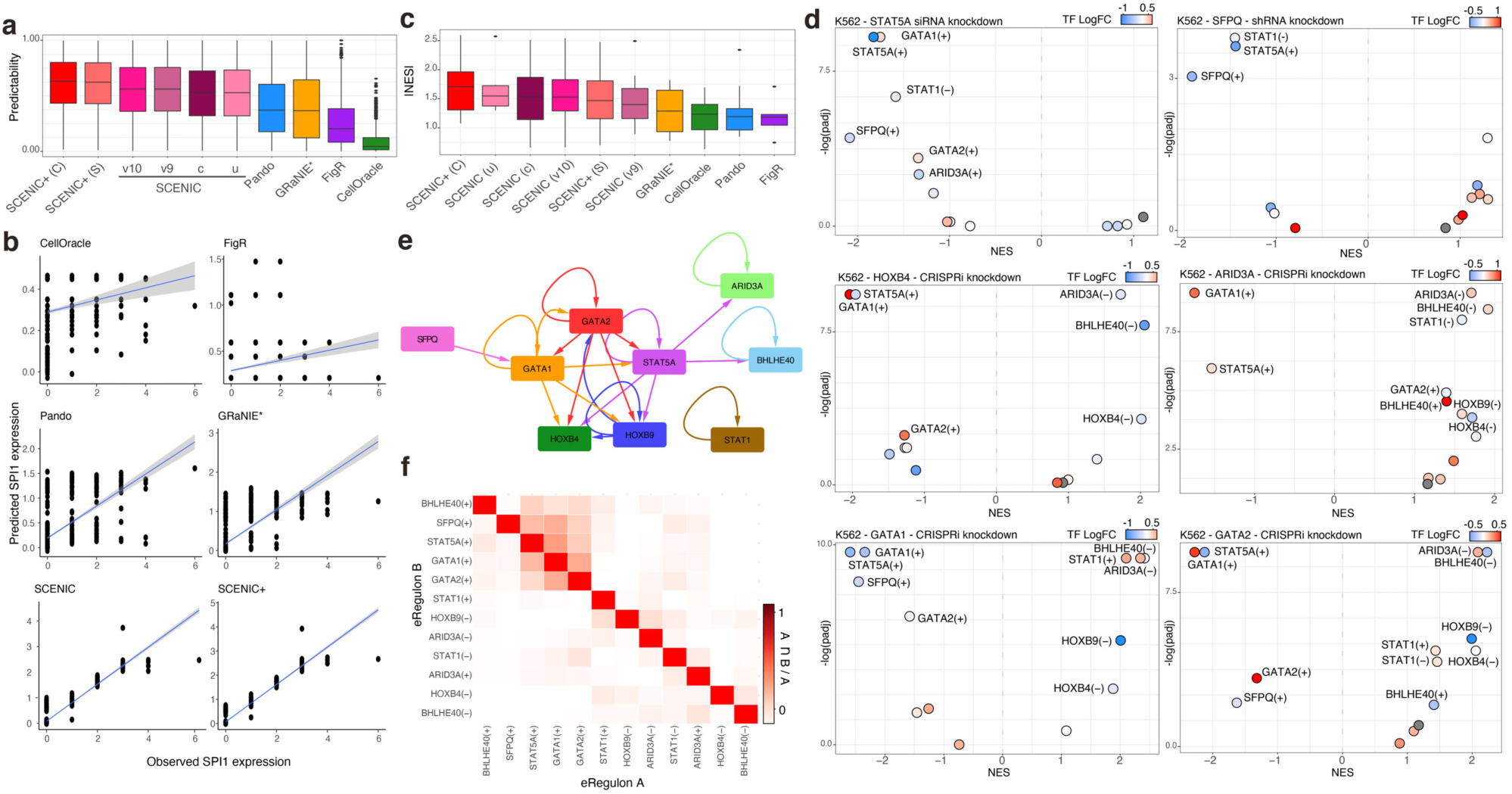
Validation of predicted target genes by different methods. **a.** Box plot depicting the correlation between observed and predicted gene expression values using the eGRNs inferred from each method. **b.** Comparison between observed and predicted values for SPI1. **c.** NES distribution based on GSEA analysis using TF knockdown data as ranking and target genes derived by each method as gene sets. **d.** Examples on K562 different TF knockdowns showing GSEA -log10 adjusted p-value and NES for different eGRNs found by SCENIC+. **e.** Network showing TF-target gene interactions for selected genes. **f.** Heatmap showing the overlap between the regions of the regulons indicated by the rows and columns, divided by the size of the regulons in the columns.

**Fig S8.**
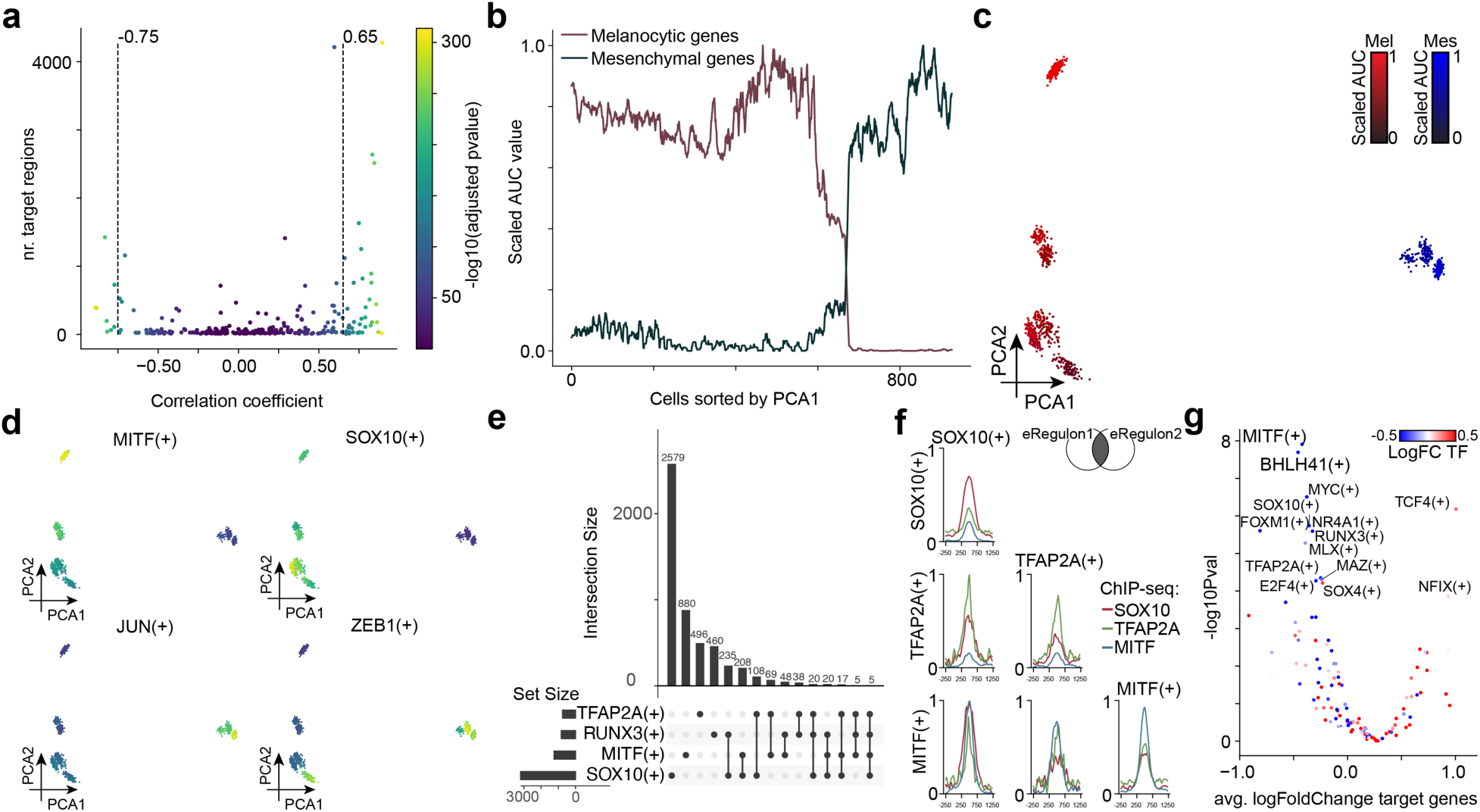
SCENIC+ analysis on cohort of melanoma cell lines. **a.** Scatter plot showing number of target regions vs TF expression-to-region AUC Pearson correlation coefficients for each eRegulon. Color scale represents -log10(adjusted p value) of the Pearson correlation coefficients. eRegulons are selected based on a threshold on the correlation coefficient, indicated by dotted line. **b.** Line plot of scaled AUC values of melanocytic and mesenchymal marker gene sets^1^ in cells sorted by PCA1. **c.** PCA dimensionality reduction of cells by eRegulon enrichment scores colored by scaled AUC values of melanocytic and mesenchymal marker gene sets. **d.** PCA dimensionality reduction of cells by eRegulon enrichment scores colored by scaled regulon AUC values of MITF, SOX10, JUN and ZEB1. **e**. Upset plot showing size and overlap of target regions of TFAP2A, RUNX3, MITF and SOX10. **f**. ChIP-seq enrichment of SOX10, MITF and TFAP2A in target regions of SOX10, MITF and TFAP2A and all combinations of two. Signal is scaled across all comparisons between 0 and 1. **g**. -log10 p-value (t-test) and average log2 fold change of target genes of eRegulons after SOX10 knockdown in MM001. Color scale encodes log2 fold change of the expression of the TF corresponding to each eRegulon after SOX10 knockdown.

**Fig S9.**
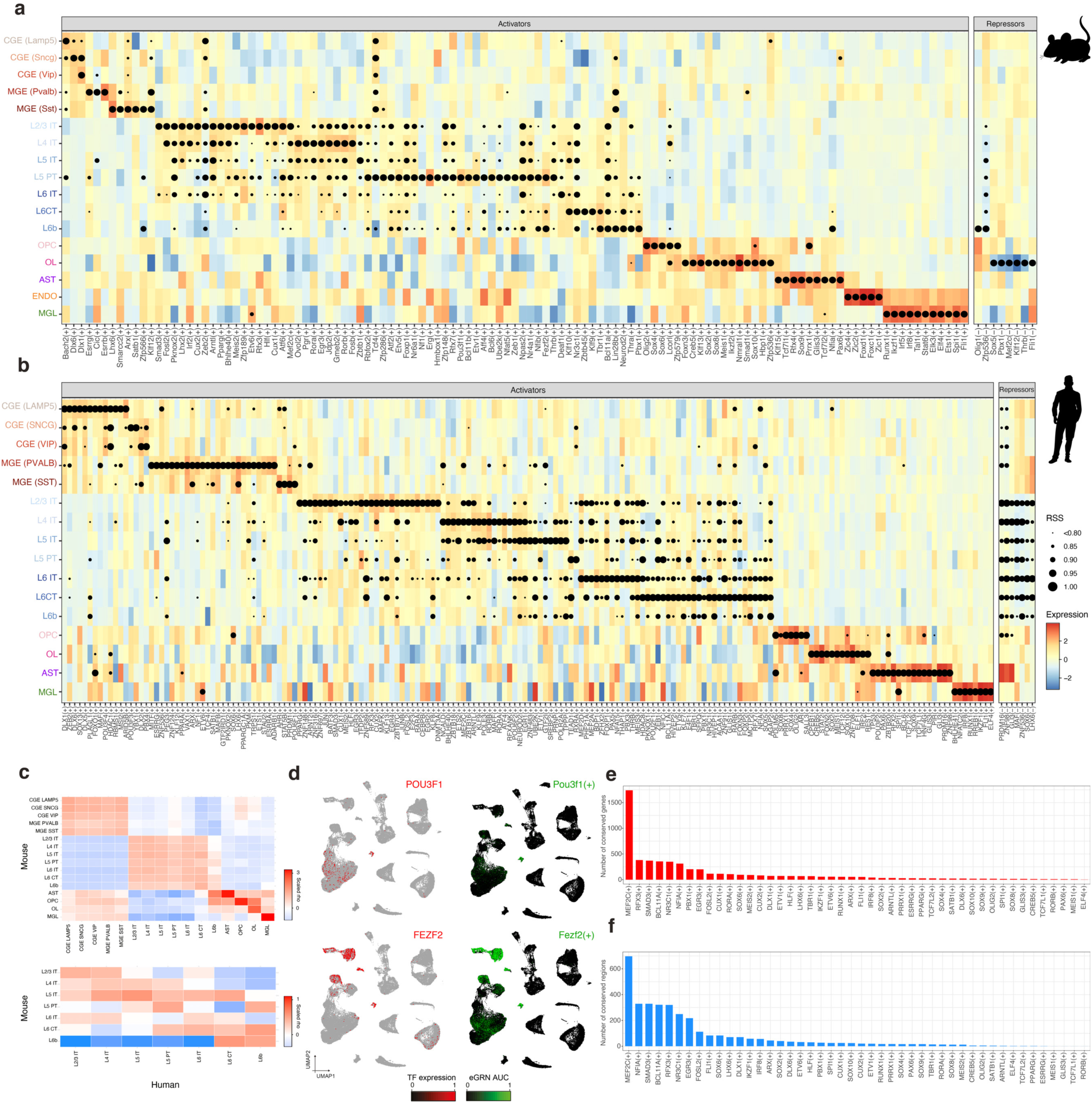
Cross-species analysis with SCENIC+ in the mouse and the human brain. **a.** Dotplot including high quality regulons in the mouse cortex showing TF expression of the regulon on a color scale and cell type specificity of the regulon (RSS) on a size scale. **b.** Dotplot including high quality regulons in the mouse cortex showing TF expression of the regulon on a color scale and cell type specificity of the regulon (RSS) on a size scale. **c.** Heatmap showing the scaled correlation between the RSS values for each regulon in each cell type. **d.** Human cortex UMAP (84,159) showing TF expression (red) and AUC enrichment of the mouse regulon (converted to human genes). **e.** Barplot showing the number of conserved genes between the matching human and mouse regulons. **f.** Barplot showing the number of conserved regulons between the matching mouse and human regulons.

**Fig S10.**
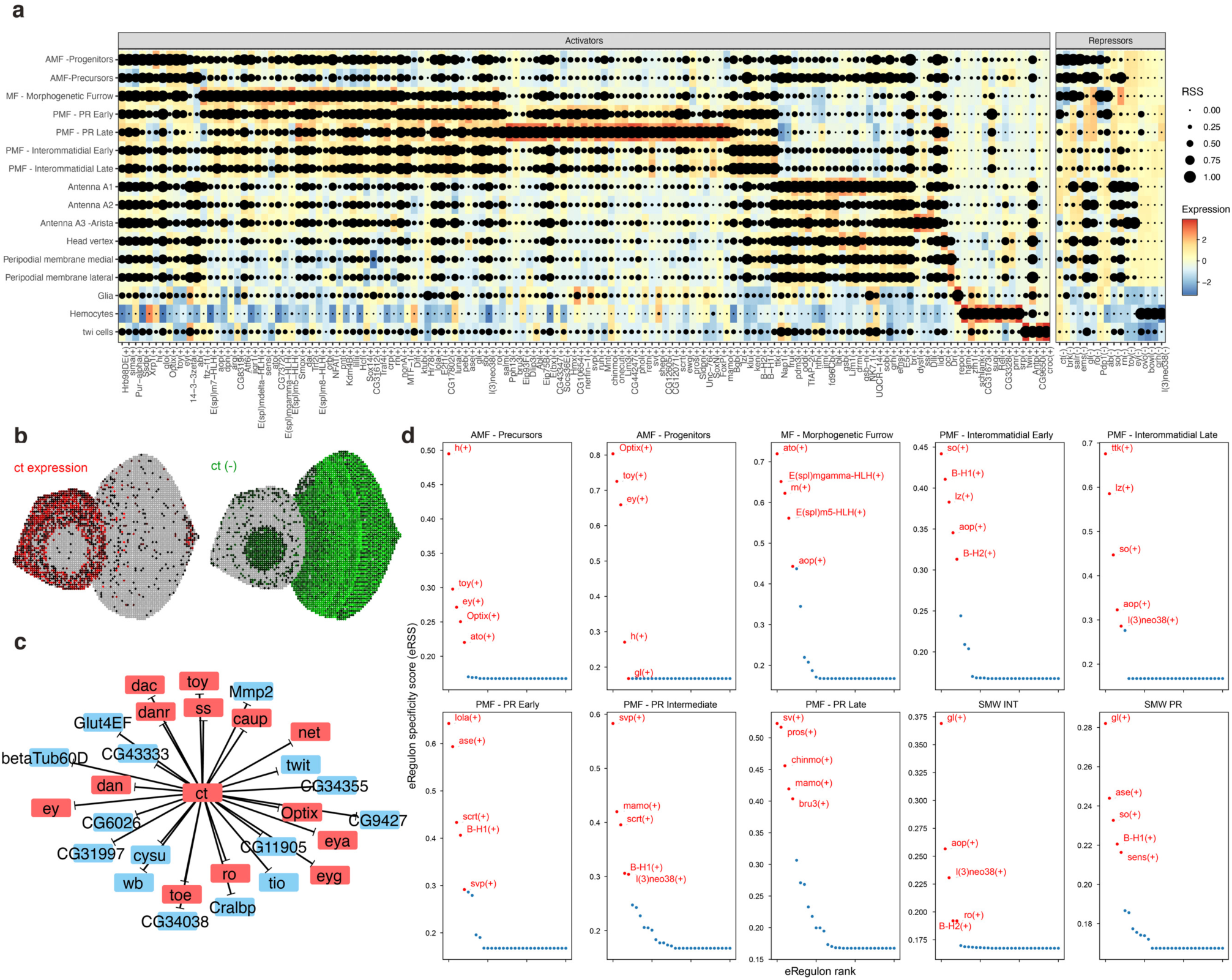
SCENIC+ analysis on the fly eye-antennal disc. **a.** Dotplot including high quality regulons in the eye-antennal disc showing TF expression of the regulon on a color scale and cell type specificity of the regulon (RSS) on a size scale. **b.** Virtual eye-antennal disc with 5,058 pseudocells colored by Ct expression and AUC values of the repressive Ct regulon. **c.** Targets of the Ct repressive regulon, showing in red targets that are transcription factors. **d.** Prioritization of differentiation forces in different stages of the eye disc differentiation.

**Fig S11.**
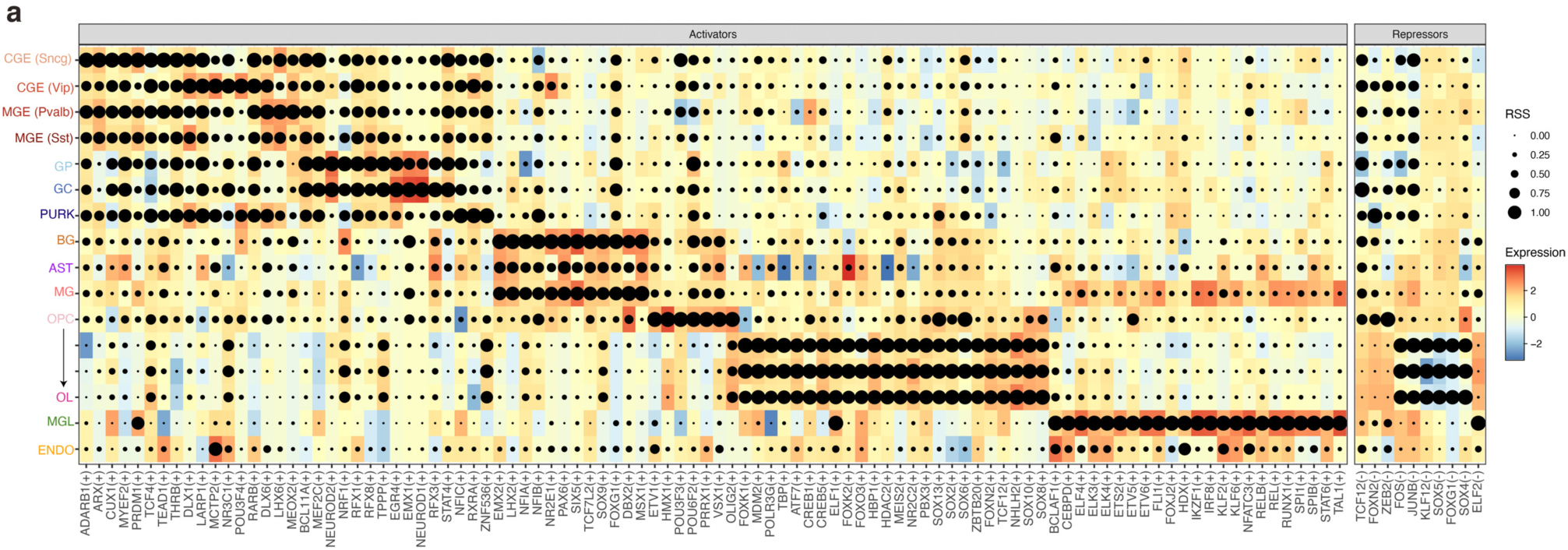
Inferred regulons by SCENIC+ in the human cerebellum. **a.** Dotplot showing TF expression of the regulon on a color scale and cell type specificity of the regulon (RSS) on a size scale.

**Figure S12.**
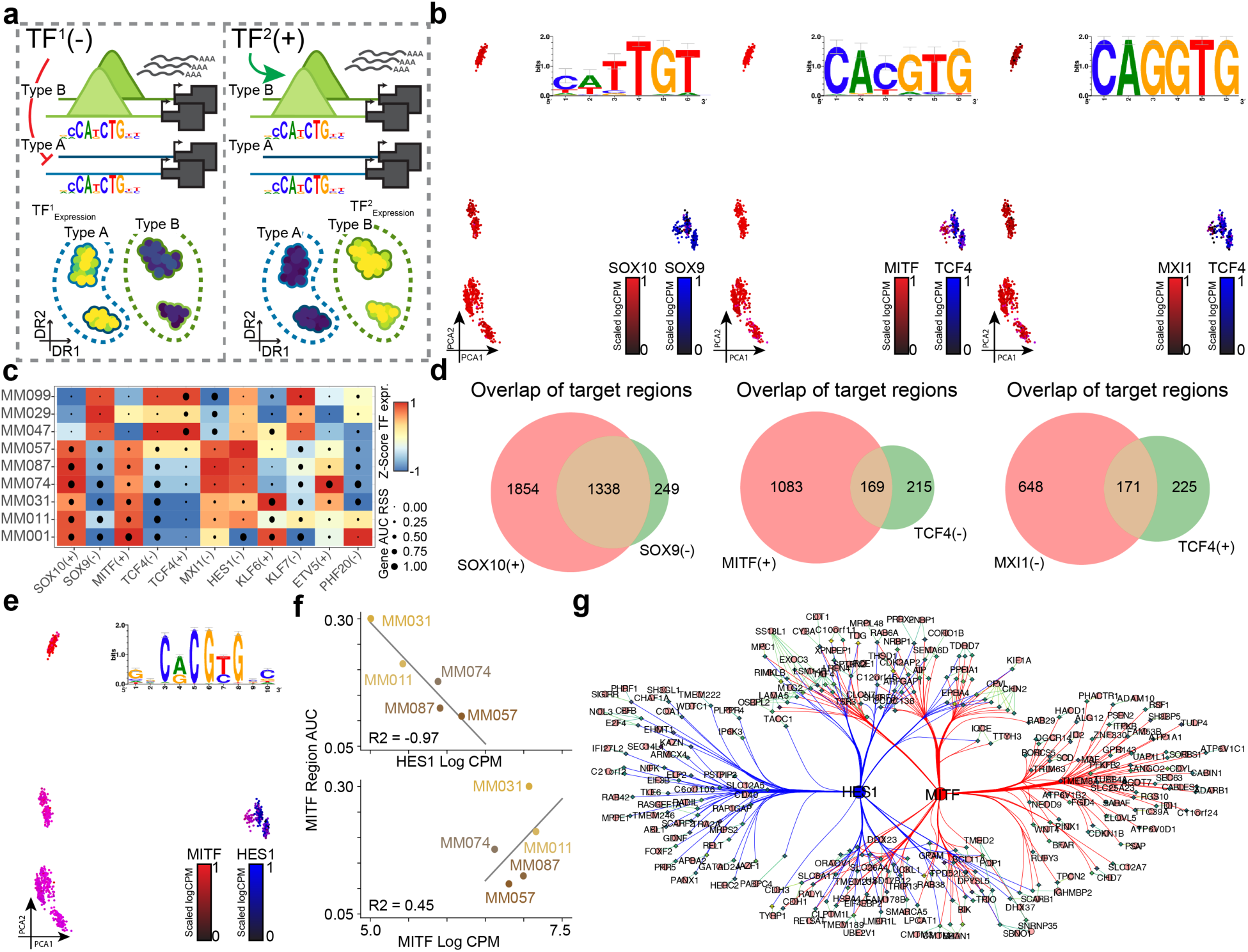
Repressor predicitons of SCENIC+ in melanoma. **a.** Illustration how TFs of the same family of which the expression is anti-correlated in system can cause spurious repressor predictions. Scenario 1 (left): TF^1^ is a potential repressor which is expressed in cell type A and actively closes chromatin in that cell type. Scenario 2 (right): TF^2^ is a potential activator of the same TF family as TF^1^ which is expressed in cell type B and opens the chromatin in that cell type. Both scenarios lead to the same gene-expression and chromatin-accessibility measurements and can thus not be disentangled if both TF^1^ and TF^2^ are present in the same system. **b.** Principal Component Analysis (PCA) projection of 936 pseudo mutli-ome cells based on cellular enrichment (AUC scores) of predicted target genes and regions from SCENIC+ eRegulons colored by the expression of SOX10 and SOX9, MITF and TCF4, MXI1 and TCF4. Shared motif used by the pair of TFs in each plot is shown on the top right. **c.** Heatmap-dotplot showing TF expression of the eRegulon on a color scale and cell type specificity (RSS) of the eRegulon on a size scale. **d.** Venn diagram showing overlap of predicted target regions of SOX10 and SOX9, MITF and TCF4; and MXI1 and TCF4. **e.** Principal Component Analysis (PCA) projection of 936 pseudo mutli-ome cells based on cellular enrichment (AUC scores) of predicted target genes and regions from SCENIC+ eRegulons colored by the expression of MITF and HES1. Shared motif used by the pair of TFs in each plot is shown on the top right. **f.** Log(CPM) expression of HES1 (top, x-axis) and MITF (bottom, x-axis) versus MITF target region AUC value (y-axis). Line fit using linear regression, least squares method. **g.** Nework showing subset of MITF and HES1 target regions. Diamonds represent regions circles represent genes. Diamonds are color coded by the average accessibility log2 Fold Change of corresponding regions in the melanocytic state.

